# Optimus: a general purpose adaptive optimisation engine in R

**DOI:** 10.1101/2022.01.18.476810

**Authors:** Nicholas A. G. Johnson, Liezel Tamon, Xin Liu, Aleksandr B. Sahakyan

## Abstract

**Motivation:** Many calculations in computational biology necessitate a use of a probabilistic optimisation protocol to determine a set of parameters that capture the system at a desired state in the configurational space. Here, we developed a flexible optimisation engine in R that can be plugged to any, simple or complex, modelling initiative through a few lucid interfacing functions, to perform a seamless optimisation with rigorous parameter sampling.

**Results:** Optimus features an acceptance ratio simulated annealing, acceptance ratio replica exchange, and adaptive thermoregulation, thus driving a Monte Carlo optimisation process in a flexible manner, through constrained acceptance frequency but unconstrained adaptive pseudo temperature regiments. We show the applicability of our R optimiser to a wide variety of problems spanning data analyses and computational biology tasks.

**Availability and Implementation:** Optimus is written and implemented in R, and is freely available from the http://github.com/SahakyanLab/Optimus repository.

**Supplementary Information:** Supplementary information with more details, tutorials, and developer instructions is available.

**Graphical Abstract:** The Optimus software logo depicting two gears (“Op”) that drag the system “s”, trapped in a “u” minimum, through a rough solution landscape into a more favourable solution with a deeper pseudo-energy minimum (“p”).

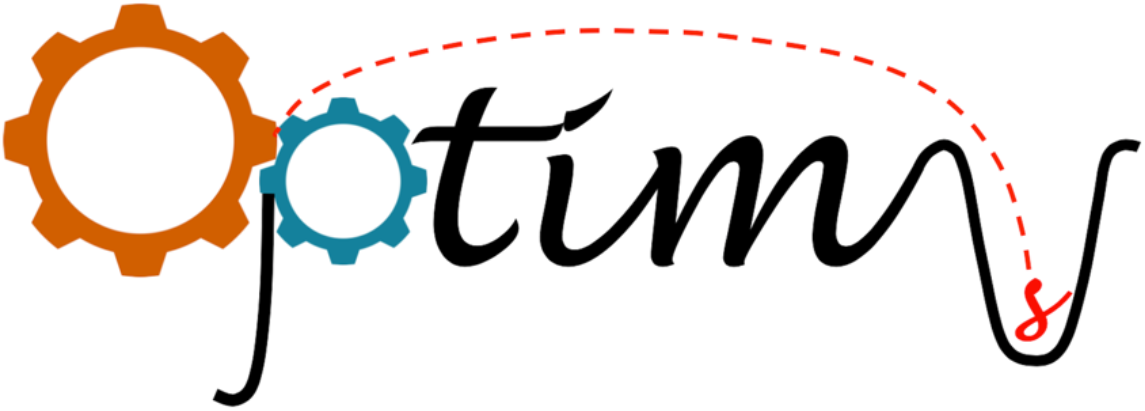

## 1. Introduction

For a complex model, where the unknown parameters cannot be determined by conventional linear or non-linear fitting techniques (Bethea *et al*., 1985; Freedman, 2009), methods for optimisation based on biased random sampling of the parameter space are the methods of choice (Cowless and Cartin, 1996). In such cases, the quality of the solution found, following some optimisation protocol, depends on that protocol’s ability to effectively explore the parameter space (Gilks *et al*., 1996) and be drawn to more favourable areas therein. Monte Carlo based algorithms are the methods of choice for such biased sampling of the parameter space, generalisable for many complex optimisation problems where the specialised algorithms are impractical to use. To meet the need of a transferrable Monte Carlo-based optimisation engine, where the flexibility of the usage is prioritised, and no major prior or runtime intervention of the core optimisation regiment is needed by a user, here we present Optimus - a universal and flexible optimisation engine in R (R Core Team, 2022) that can interface with a wide variety of modelling initiatives through a small R scripting interface. Optimus does not need any situational tunning of the pseudo-temperature scaling while defining the acceptance probabilities, even if the same system is trapped into a significant local minimum. This is achieved by driving the optimisation process through the annealing or replica exchange of the constrained factual acceptance ratio, while automatically and freely adapting the unconstrained pseudo temperature of the system (**Fig. 1**). Below is the brief description of the package and its major distinctions, all further detailed in the **Supplementary Information** that also brings five example tutorials to showcase the usage of the engine in a variety of usage scenarios.

**Figure 1.**
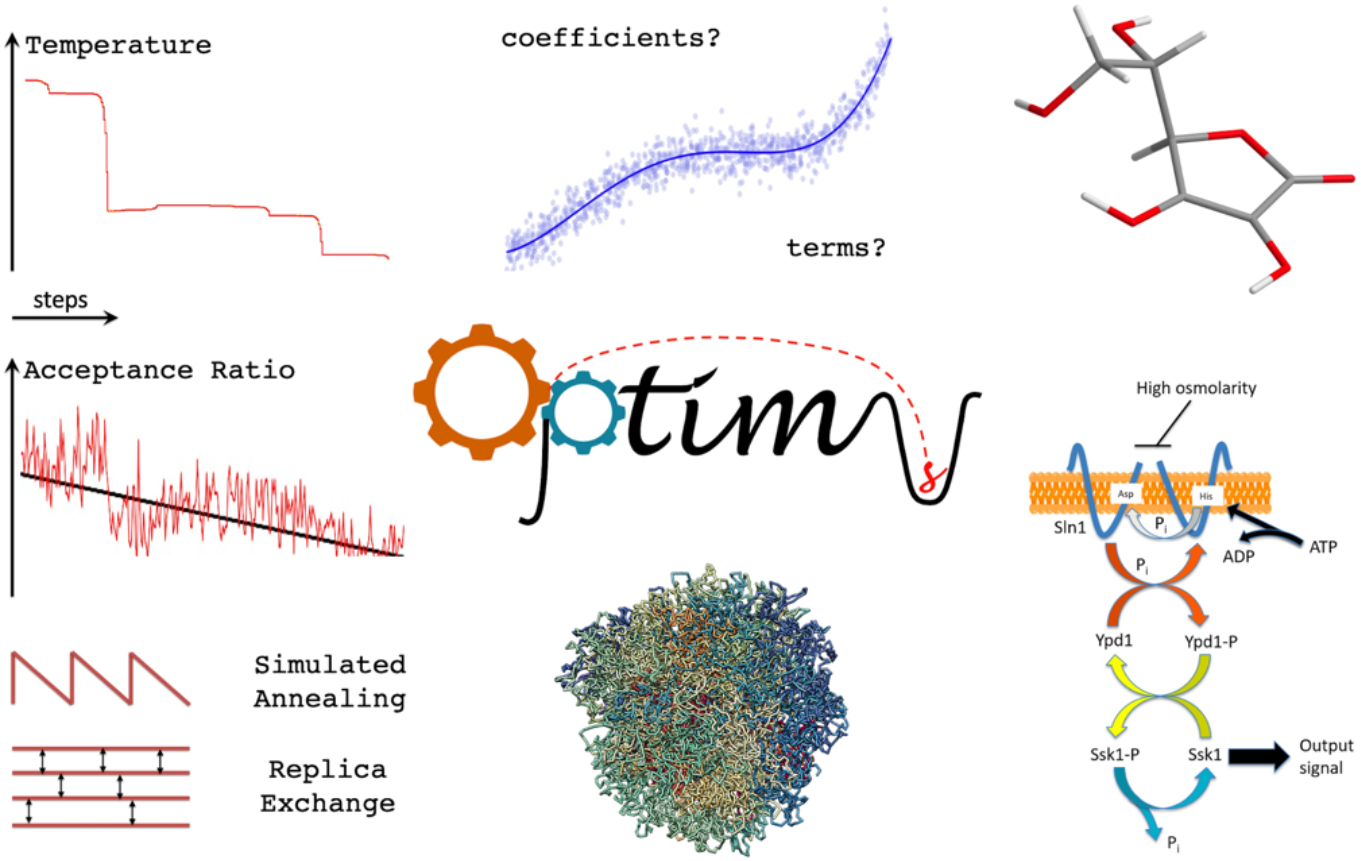
Optimus logo and a schematic illustration of its capabilities and a number of computational biology optimisation objectives, where Optimus engine is showcased in the **Supplementary Information**. The reduced plots on the left side represent the abrupt adaptive changes in pseudo temperature, automatically performed in Optimus as necessitated by the evolving system to keep up with the desired gradual decrease in acceptance ratio during the last acceptance ratio annealing cycle of a representative run.

## 2. Software description

If using the acceptance ratio (AR) simulated annealing (Kirkpatrick, 1984) (AR-SA) mode of Optimus, in a given annealing cycle, Optimus constructs a linear target AR regiment across each step. This is done based on an initial target AR, a final AR, and the number of iterations in each cycle for a given optimisation run (all of which can be modified from defaults as advanced inputs). Once the optimisation process begins, Optimus calculates an observed AR at the end of each window of a fixed number of steps by calculating the fraction of the accepted moves from all the past trials in the current window at a given initial pseudo temperature. Thereafter, Optimus compares the observed AR with the pre-defined target AR based on the annealing schedule, and determines whether and how to alter the system pseudo temperature (adaptive thermoregulation) to align the observed AR with the target ratio at the end of the following statistics window (**Fig. 1**). Thus, by employing AR-SA and adaptive thermoregulation, Optimus is able to methodically explore the parameter space even when no smooth relationship exists between the parameters and system pseudo energy, avoiding a danger to fall into traps with no crossing with conventional methods under constrained temperature range (Ingber, 1993).

Optimus additionally supports acceptance ratio replica exchange (AR-RE) as an optimisation mode, provided that the user has access to multiple processors (preferably 8 or more). This additional mode was adapted from the parallel tempering/replica exchange methodologies (Swendsen and Wang, 1986; Sugita and Okamoto, 1999). Optimus modifies the Monte Carlo flavour of the replica exchange approach to apply to any arbitrary optimisation problem with two primary modifications: the use of varying replicas of fixed AR optimisations, and necessitated unconditional exchange after two candidate configurations in those replicas are selected for an exchange (Ballard and Jarzynski, 2009). These changes are only acceptable if relaxing the equilibrium sampling criteria, whereby the parameter space can be more extensively explored for the sole reason of finding an optimal solution. This approach produces good results, as illustrated in the tutorials, and is a viable alternative to the AR-SA mode in Optimus.

## 3. Conclusion

In cases where *i*) we do not deal with energies and temperatures that emulate real physical systems; *ii*) we are only interested in final optimised configuration of the system, and do not need to characterise the statistical ensemble of reachable states/solutions, we can then cross barriers in the solution energy landscape by annealing or replica exchanging based on a metric (AR) that is more transferable to different volatile system configurations. We demonstrate the usability and flexibility of Optimus in 5 example tutorials (**Fig. 1** and **Supplementary Information**): **1)** optimising coefficients to a known equation (Tutorial 1 in **Supplementary Information**), **2)** optimising an equation itself by attempting various configurations of its functional form (Tutorial 2 in **Supplementary Information**), **3)** optimising a geometry of a molecule while interfacing with an external quantum chemical program (Tutorial 3 in **Supplementary Information**), **4)** optimising parameters to coupled ODEs from systems biology (Tutorial 4 in **Supplementary Information**), and **5)** optimising a highly specific constrained shuffling problem from 3D genome biology (Tutorial 5 in **Supplementary Information**). All these are showcased in **Supplementary Information**, with the full walkthrough of both the AR-SA and AR-RE variants. We successfully used this program in a number of research objectives, including in a recently published tuning of a model for the inference of iDNA stabilities (Cheng *et al*., 2021), and we hope it will be equally useful and accessible for other researchers.

## Acknowledgement

The Sahakyan Laboratory has been supported by the UK Medical Research Council (MRC), MRC Strategic Alliance Funding (MC_UU_12025). NAGJ is grateful to the Princeton University International Internship program for supporting his internship in Oxford, LT is grateful to the Jardine Foundation for supporting her DPhil studies.

## Supplementary Information

### Introduction

#### Motivation

For a complex model, where the unknown parameters cannot be determined by conventional linear or non-linear fitting techniques, methods for optimisation based on biased random sampling of the parameter space, are the methods of choice (Cowles and Carlin 1996). In such cases, the quality of the solution found, following some optimisation protocol, depends on that protocol’s ability to effectively explore the parameter space (Gilks, Richardson, and Spiegelhalter 1996) and be drawn to more favourable areas therein. Many existing methods perform well for certain modelling scenarios, but fail in others due, in part, to an inefficient exploration of the parameter space and easy trapping into local minima with regard to the metrics for the quality of the found solutions. In this manual, we present Optimus, a universal Monte Carlo optimisation engine in R with acceptance ratio annealing, replica exchange and adaptive thermoregulation. It can universally interface with any modelling initiative through its interfacing functions, and optimise the model parameters by effectively exploring the parameter space. Controlled by a user, Optimus can execute either an annealing or a replica exchange procedure, however, both driven by acceptance ratio adjustment, rather than by altering pseudo temperature.

This manual will begin with a basic overview of Monte Carlo optimisation and the common temperature-based simulated annealing framework. It will then proceed with a presentation of the acceptance ratio annealing procedure of Optimus, its adaptive thermoregulation feature, and its replica exchange procedure. After an explanation of how to download the Optimus R package from GitHub and install it locally, five stand-alone tutorials will be presented to illustrate how users should employ Optimus and to demonstrate its flexibility as an optimisation engine across a wide variety of optimisation scenarios. Finally, an “Advanced User Manual” section is included, in which all possible and tested input parameters to Optimus are outlined and the output formats are detailed.

#### Briefly on Monte Carlo and Temperature-Driven Simulated Annealing Procedure

Let us assume that our model is a certain function *m*() that performs operations on the inputted **K** coefficient set and returns the observable object *O* = *m*(**K**). Our task is to optimise **K**, the set of coefficients, so that the error metric *u*(*O*) that measures the violations from the target *O^trg^* set by the model-generated *O* is minimal. In a Monte Carlo optimisation procedure, we can define a pseudo-energy *e*(*u*) of the system as a function of *u*(), where lower values of the pseudo energy *e* correspond to better candidate solutions **K**. In order to find a better set of **K**, we need to alter it by a certain rule *r*() that, if repeated many times, enables the sampling of the parameter space for **K**. One can then evaluate the pseudo energies before and after the alterations:

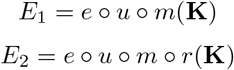

We then accept or reject the alteration move, meaning we accept the new set of *r*(**K**) coefficients as the new **K** or revert back to the previous **K**, guided by the following acceptance probability, as postulated in a Metropolis criterion (Chib and Greenberg 1995), (Chen and Roux 2015):

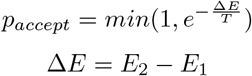

where T is the pseudo temperature, that should be always greater than 0. For a given Δ*E* > 0 energy difference, one would have a certain stringency for accepting the alteration move, depending on the value of the pseudo temperature *T*. Therefore, if the Monte Carlo simulation was to modify the **K** set to a state where any further move would increase the pseudo energy at a great enough amount for the moves to be almost always rejected, then one could overcome that state and further sample the other values in the parameter space by increasing the value of the pseudo temperature.

To this end, one way to overcome the barriers is by using the technique known as simulated annealing, where we gradually anneal the temperature from a higher value to lower during the course of the simulation (Kirkpatrick 1984). The parts of the simulation where the pseudo temperature is higher, allows relatively unconstrained exploration of the parameter search space, whereas those parts with a lower pseudo temperature limit the search to a more local area of the energy landscape and associated parameter space. Multiple cycles of this annealing procedure can thus be executed to increase the overall sampling.

#### Acceptance Ratio Simulated Annealing Procedure

A significant limitation of the pseudo-temperature-based simulated annealing, while used only in an optimisation objectives, is that a given scheme of temperature annealing might be efficient for some models or pseudo-energy metrics, but not efficient for others (Ingber 1993). A temperature for a given system at a given point of the annealing cycle that is designated to be quite permissive in terms of accepting the moves, can actually not be permissive for some other models or systems simulated. This depends on the value and scale of the Δ*E* energy difference, as can be seen in the equation for *p_accept_*, and can be affected by any of the *e*(), *u*(), *m*() components, and the system configuration **K**. This necessitates a pre-adjustment of the pseudo-temperature values from one modelling objective to another. Furthermore, even within a single model optimisation procedure on a defined system, the pseudo-energy metric can shift into a scale (due to a significant alteration/shift in the system as per the alteration rule *r*()) that does not match with the selected temperature scheme anymore, leading to a poor sampling of the parameter space when optimisation is at stake within restricted time and computational capacity. This can often be the case when the pseudo energy of the system does not exhibit a smooth dependency on **K**, loosely meaning that close states of **K** do not necessarily produce close pseudo-energy values (for instance, due to an alteration rule that randomly expands or trims the system, such as when equation terms are randomly turned on or off).

To this end, in general cases where

A. we do not deal with energies and temperatures that display the smoothness or roughness emulating real physical systems, such as while simulating molecular systems;
B. we are only interested in final optimised configuration of **K**, and do not need to characterise the statistical populance as an ensemble of reachable states/solutions,

we can anneal a metric for crossing different barriers that is more transferable to different systems and system configurations. As such a metric, Optimus uses the acceptance ratio, expressed as a fraction of accepted moves within certain number of past steps (`STATWINDOW`).

In a given annealing cycle, Optimus constructs a linear target acceptance ratio regiment for each step. This is done based on an initial target acceptance ratio, a final acceptance ratio, and the number of iterations in each cycle for a given optimisation run (all of which can be specified as inputs). Once the optimisation process begins, Optimus calculates an observed acceptance ratio at the end of each `STATWINDOW` (a fixed number of steps which can be specified by the user) by calculating the fraction of the accepted moves from all the past trials in the current `STATWINDOW`. Thereafter, Optimus compares the observed acceptance ratio with the pre-defined target acceptance ratio based on the annealing schedule and determines whether and how to alter the system pseudo-temperature (adaptive thermoregulation) to align the observed acceptance ratio with the target ratio at the end of the following `STATWINDOW`. Thus, by employing acceptance ratio annealing and adaptive thermoregulation, Optimus is able to methodically explore the parameter space for **K** even when no smooth relationship exists between **K** and the system pseudo energy.

#### Adaptive Thermoregulation

All decisions governing the adaptive pseudo-temperature alterations on the system are made by a Temperature Control Unit (TCU) in Optimus. Note, that the TCU is completely encapsulated as a backend unit, such that modifications can be easily made by advanced users if needed. This section articulates the exact protocol followed by the default state of the TCU.

The initial system temperature is specified as an input argument. At the end of each `STATWINDOW`, if the observed acceptance ratio is within a fixed range `T.DELTA` (specified as an input argument) of the target acceptance ratio based on the annealing schedule, the TCU will make no change to the current system pseudo temperature. If the observed acceptance ratio is less than the ideal ratio and outside the range of `T.DELTA`, the TCU will increase the system pseudo temperature by a value `T.ADJSTEP` (the initial value of `T.ADJSTEP` is specified as an input argument). Similarly, if the observed acceptance ratio is greater than the ideal ratio and outside the range of `T.DELTA`, the TCU will reduce the temperature by a value `T.ADJSTEP`.

If the observed acceptance ratio keeps being below the ideal acceptance ratio for `TSCLnum` (an integer input argument) number of subsequent `STATWINDOW`s, `T.ADJSTEP` will be increased by a factor `T.SCALING` (an input argument). Similarly, `T.ADJSTEP` will also be decreased by a factor `T.SCALING` if the observed acceptance ratio is greater than the ideal acceptance ratio for `TSCLnum` subsequent `STATWINDOW`s. `T.ADJSTEP` is reset to its original input value whenever a series of subsequent observed acceptance ratios being greater than/less than ideal acceptance ratios is broken. If ever the TCU subtracts `T.ADJSTEP` from the current temperature and the result is a negative value, the system pseudo temperature is set to `T.MIN` (an input argument). The final feature of the TCU is that although the initial system pseudo temperature is specified by the user, if multiple annealing cycles are employed, the initial pseudo temperature for acceptance ratio annealing cycles, after the first cycle, is inferred from the decisions of the TCU on previous cycles.

The above collection of TCU decision rules with their default values result in pseudo-temperature modulations that assure the observed acceptance ratios during Optimus runs to follow the ideal acceptance ratios remarkably well. Moreover, as will be highlighted in the five tutorials below, large temperature alterations are often required to align the observed acceptance ratios with the ideal ratios, a task which Optimus excels at, whereas other protocols would have difficulties and result in poor parameter sampling because of fixed or gradually changing temperature regiments.

#### Acceptance Ratio Replica Exchange Procedure

Optimus additionally supports acceptance ratio replica exchange as an optimisation mode, which can be selected in place of acceptance ratio annealing, provided that the user has access to multiple processors (ideally at least 4, and preferably 8 or more). The inspiration for this additional mode was taken from the parallel tempering Monte Carlo techniques and Replica Exchange Molecular Dynamics (REMD) simulations (Sugita and Okamoto 1999). In particular, REMD simulation is a technique employed to obtain equilibrium sampling of a molecular system (for instance, a protein) at a certain fixed (usually the lowest in a range) temperature. Let *T* = {*T*_1_, *T*_2_, …, *T_n_*} be a set of *n* distinct temperatures for which *T*_1_ < *T*_2_ < … < *T_n_*. In REMD, *n* replicas of molecular dynamics simulations for a given system are initialised at each *T_i_* ∈ *T*. Note, that each temperature *T_i_* corresponds to a slightly different potential of the examined system to roam the energy landscape and cross the barriers. States possible to populate at a temperature *T_i_* can become a bit more accessible at a temperature *T*_*i*+1_ and so on. If the state configurations in adjacent replicas are allowed to exchange, the simulation will be able to overcome energy barriers at various temperature replicas and thoroughly explore the parameter space. Moreover, for configuration *x_n_* in replica *T_i_*, and configuration *x_m_* in replica *T*_*i*+1_, equilibrium sampling is achieved by selecting appropriate exchange probabilities (Sugita and Okamoto 1999), generally expressed as:

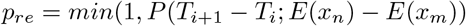

Thus, overall, replica exchange simulations can be executed by simulating *n* replicas of a simulation at distinct temperatures simultaneously and independently, next randomly selecting two configurations in adjacent replicas and exchanging them with probability *p_re_*.

Optimus modifies the Monte Carlo analogue of the replica exchange approach to apply to any arbitrary optimisation problem with two primary modifications. Firstly, due to the aforementioned transferability of utilising acceptance ratio as a controlling metric rather than pseudo temperature in arbitrary systems, Optimus initialises *n* replicas with different fixed target acceptance ratios as opposed to fixed temperatures, and uses the previously described TCU for the adaptive thermoregulation to keep up with the fixed acceptance ratio values for varying **K** within a given replica. Secondly, since Optimus is only concerned with finding an optimal solution and is not compliant with any equilibrium sampling of the parameter space (see the A and B criteria described above), after two candidate configurations are selected for exchange, they are necessarily exchanged (this can be viewed as setting *p_re_* =1 in the above procedure) (Ballard and Jarzynski 2009). The exchange regiment is controlled through the input, with a default optimal value that works well in most cases. By relaxing the equilibrium sampling criterion, the parameter space can be more extensively explored for the sole reason of finding an optimal solution. This approach produces good results, as illustrated in the tutorials, and is a viable alternative to the acceptance ratio annealing mode in Optimus optimisations, when the user has access to better computational resources.

### How to Cite

The usage of Optimus should be accompanied with the following citations:

- Nicholas A.G. Johnson, Liezel Tamon, Xin Liu and Aleksandr B. Sahakyan, *Optimus: a general purpose adaptive optimisation engine in R, bioRxiv*, this deposition, **2022**.
- Nicholas A.G. Johnson, Liezel Tamon, Xin Liu and Aleksandr B. Sahakyan, http://github.com/SahakyanLab/Optimus, **2018**.

### Installation Instructions

Installing Optimus locally for immediate use requires only R and a connection to the Internet. After opening an R client, execute the commands below to install Optimus. Note, that the latest version of Rtools is required for this installation to work. If it is not available locally and the installation is attempted from within RStudio, a prompt will appear to download and install Rtools. After following those instructions, restart the RStudio session before reattempting the Optimus installation.

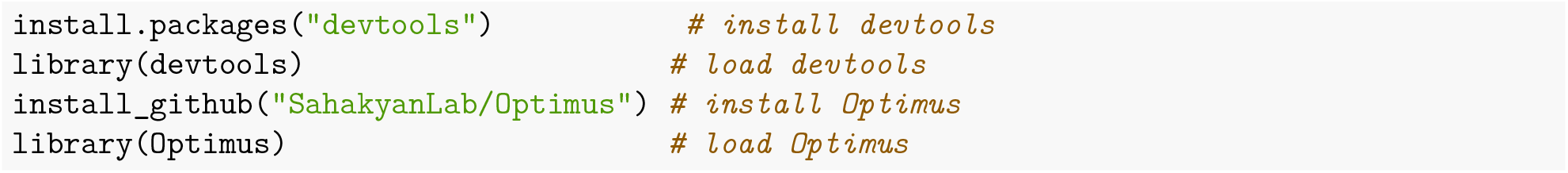

### Execution Instructions

Optimus optimisation engine works seamlessly and requires little to no intervention from the user while applied to a wide variety of optimisation tasks. The part where the user must intervene is the specification of **K**, and the R functions corresponding to *r*(), *m*(), and the combined *e* ∘ *u*(). Those objects serve as the application interface of the Optimus engine to any optimisation task. The port through which additional user data or objects can be supplied to Optimus, in case those are needed in the specification of a model, is implemented through the `DATA` argument that is passed to the model function `m()` and `u()`. All the interfacing functions and objects have a minimal and well defined internal compliance rules in Optimus, and otherwise can be flexible and as simple or complicated as the user requires, to properly plug Optimus into a desired modelling objective. The whole optimisation procedure with acceptance ratio annealing or replica exchange is then fully taken care of by the Optimus engine.

The five tutorials below showcase the usage of Optimus in five, broadly different scenarios, with a particular focus on how to write the interfacing functions to plug the Optimus engine, and how the acceptance ratio annealing mode performs in comparison to the acceptance ratio replica exchange one. The examples also cover a usage scenario where an external, more complicated, program is linked to the R procedure. Furthermore, we provide a built in function `OptimusExamples()`, which can generate the necessary inputs for any of the mentioned five tutorials, ready for modifications to tailor those to particular needs. The `OptimusExamples()` function requires, at minimum, the tutorial identifier (1 to 5) along with the methodology specification (simulated annealing - SA; replica exchange - RE), in order to generate the interfacing Optimus input, which by default is saved as an `example.R` file within the current directory (all alterable).

The recommended way to use Optimus would thus be by exploring and understanding the below tutorials, finding the example closest to one’s objective, generating the input for that example using `OptimusExamples()`:

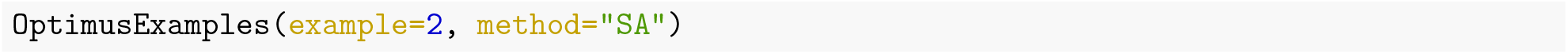

then modifying the input objects to your specific needs and workflow.

### Tutorial 1: Finding Coefficients for a Polynomial Function

#### Problem Statement

In this example, we shall use Optimus to find the coefficients of the polynomial function that is known to represent the observations *y* the best. This, of course, is a simple task that can be addressed more robustly by least-squares linear model fitting. However, by starting with this example, we shall focus our attention on the organisation of the Optimus input, rather than the complexity of the task.

First of all, let us create some data for the example.

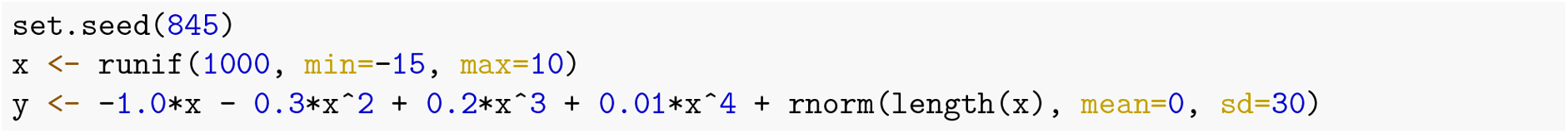

The good side of this noisy data generation is that we know the original function that describes it: *y* = – 1.0*x* – 0.3*x*^2^ + 0.2*x*^3^ + 0.01*x*^4^. Hence, we can check how well Optimus performs at finding the correct coefficients. The synthetic “real world” noisy data that we generated looks like this:

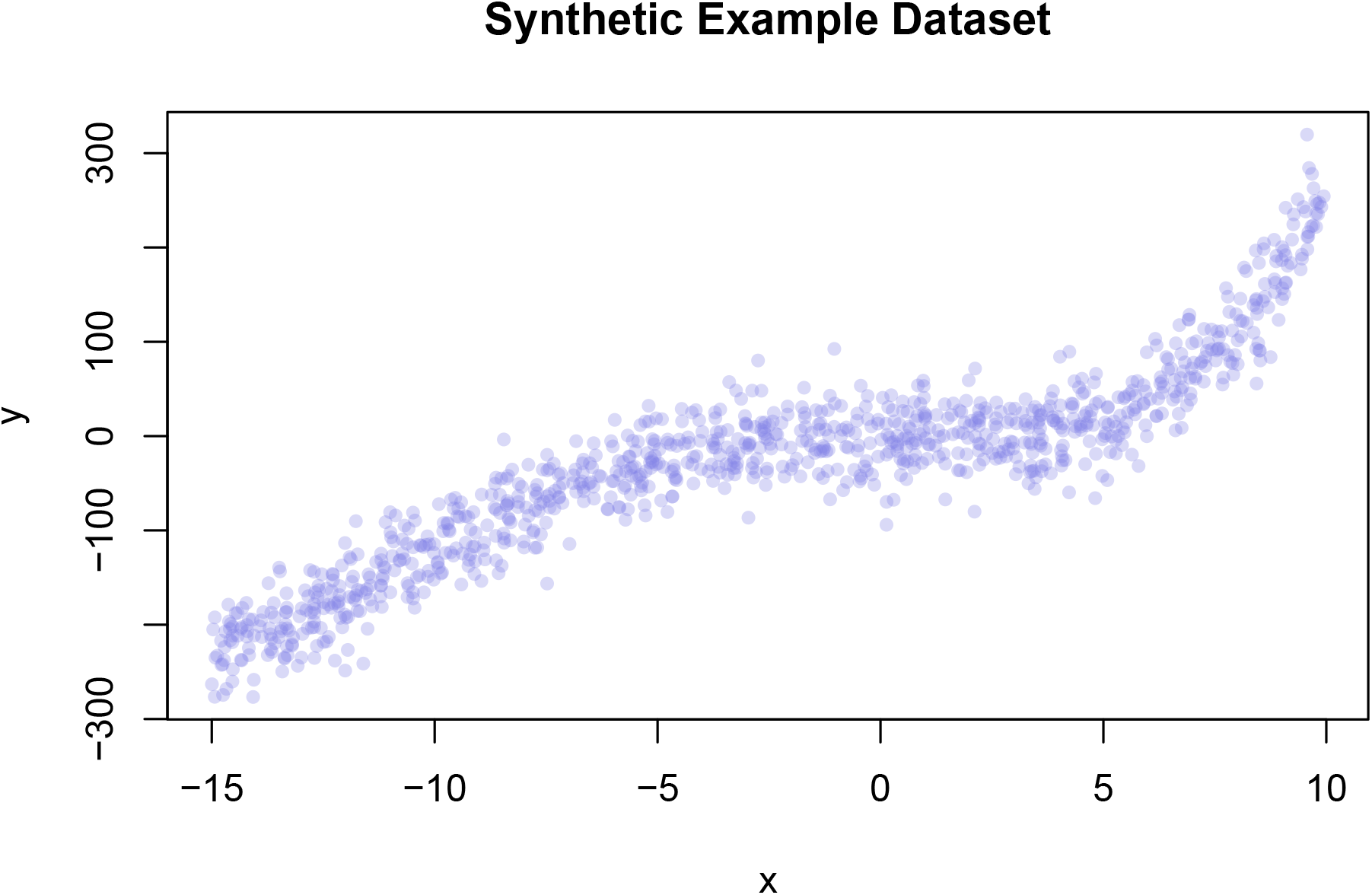

Before we turn to Optimus, let us see how the proper linear model fitting will perform using this data.

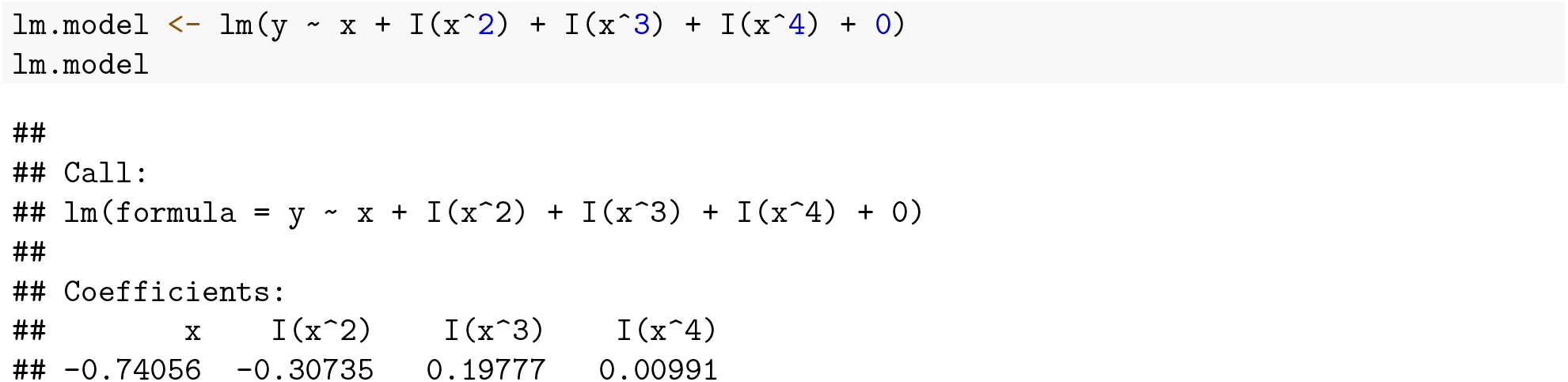

The least-squares linear model fitting for the coefficients to the known functional form is, not surprisingly, rather close to the original equation *y* = –0.741*x* – 0.307*x*^2^ + 0.198*x*^3^ + 0.010*x*^4^:

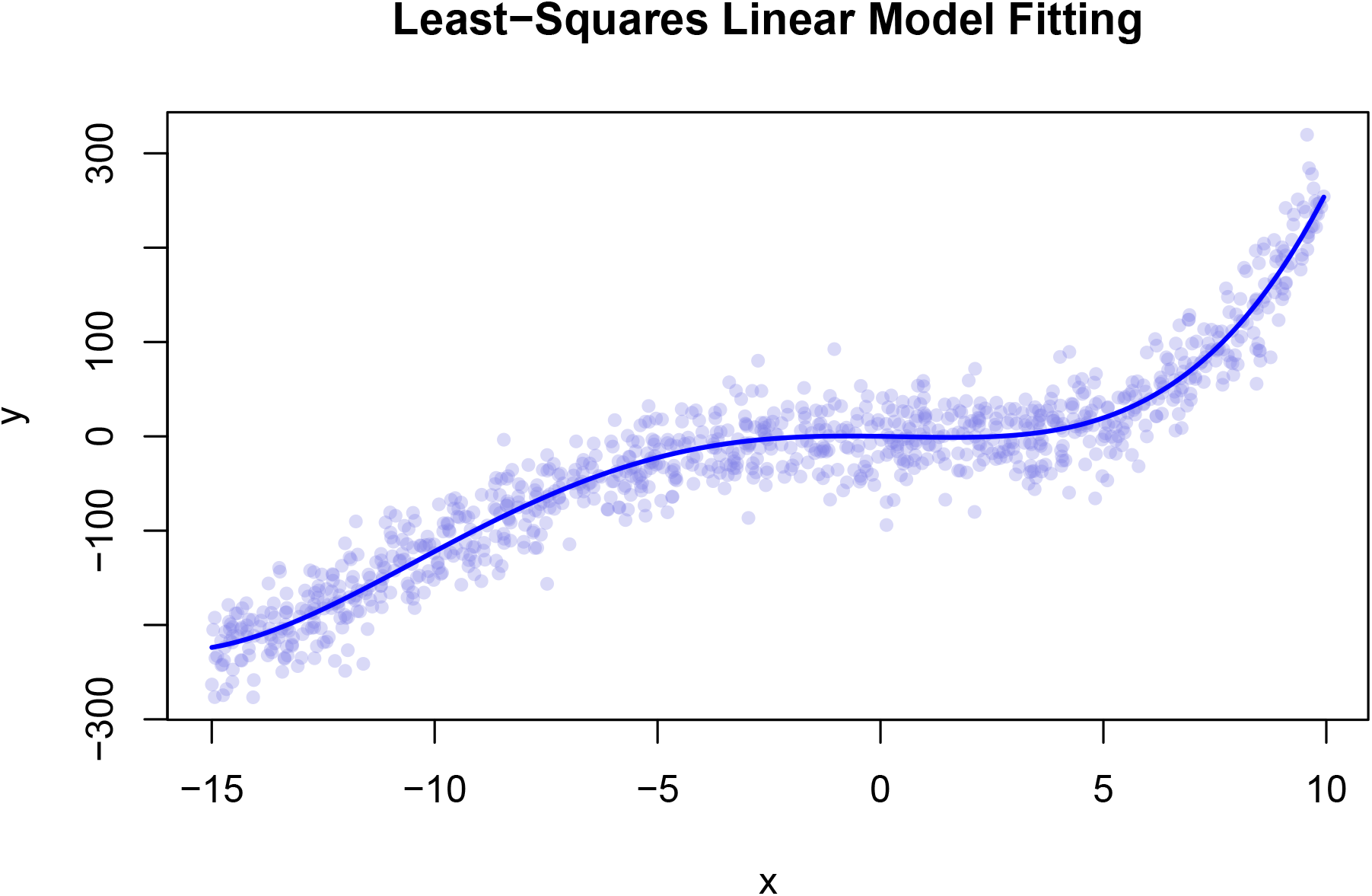

The root mean squared deviation (RMSD) between the observed data *y* and the linear model fitting outcome is:

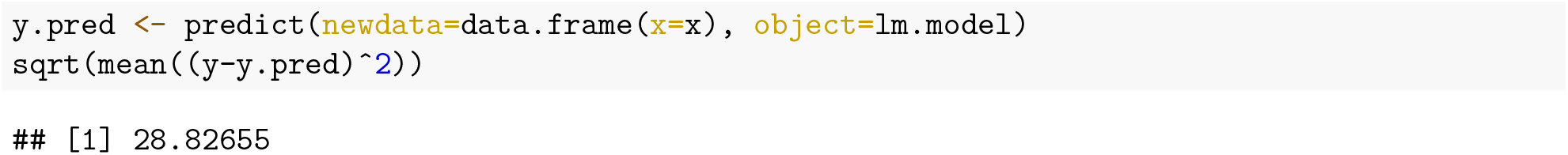

which is even slightly better in describing the noisy data, as compared to the maximum possible RMSD based on the de-noised data:

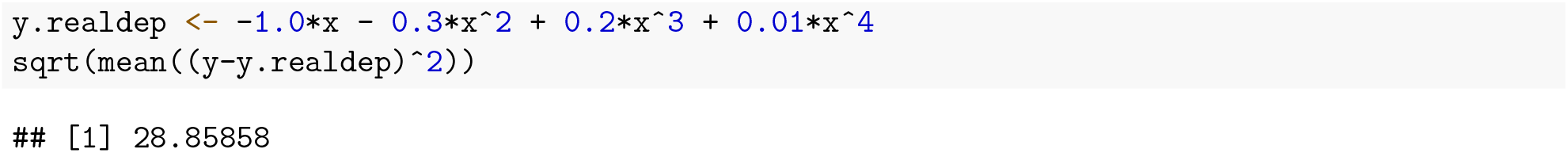

#### Defining Optimus Inputs

Now we can set up the inputs for the Optimus run. We shall use the model *k*_1_*x* + *k*_2_*x*^2^ + *k*_3_*x*^3^ + *k*_4_*x*^4^ to fit the *y* observables based on the values for *x*. The dependent functions that are needed for setting up an Optimus run, are given as inputs in the Optimus function call.

First, we need to create the essential object `K`, which stores the initial values for the parameter(s) to be optimised. `K` can be an object of any type. From a single numeric or character value to a vector of values or a data frame holding, say, Cartesian coordinates of a molecule to be optimised. The only Optimus requirement from `K` is that it should be something alterable (*via* a rule function `r()`, see below) and should influence the outcome of another required model function `- m()` (see below). In this example, we have 4 coefficients to optimise from some random initial state. We can thus make `K` be a numeric vector of size 4. Let us start from all the components being 1.0, which, as entries in `K`, can be both named and unnamed. Though not the case here, the entry-named data for `K` can be essential for some models that specifically use coefficient names, for instance when a system of ODEs is used in the model function `m()` in one of the tutorials.

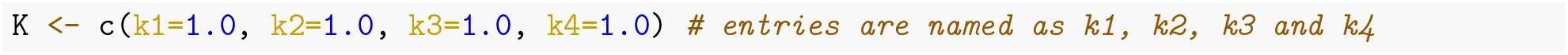

Second, we should create the function `m()` for the model. The function `m()` should be designed to operate on the whole set of parameter snapshot `K` and return the corresponding observable object `O`. Please note that the size and shape of `K` and `O` are not necessarily to match, depending on the nature of the model used. Operating on `K` is **one of the hard conditions** on `m()`, which can optionally operate on other data as well. In our situation, the function `m()` should operate on the provided instance of four coefficients (in the object `K`), and, additionally on the values *x*. It should then return a vector of observations `O` (to be compared with *y* target observations) of the same size as vector *x*. Any additional data required by the model, in our case an object with the set of 1000 *x* values, must be provided to the function in an input variable `DATA`, a list holding the additional data that must be accessed by `m()` and `u()` (see below). The variable `DATA` **must be provided** to Optimus in the function call, and `m()` **must take it** as an input. In the case that neither `m()` nor `u()` require additional data, the two functions should still be created such that they take a variable `DATA` as an input, and the variable `DATA` passed to Optimus will be set to `NULL`).

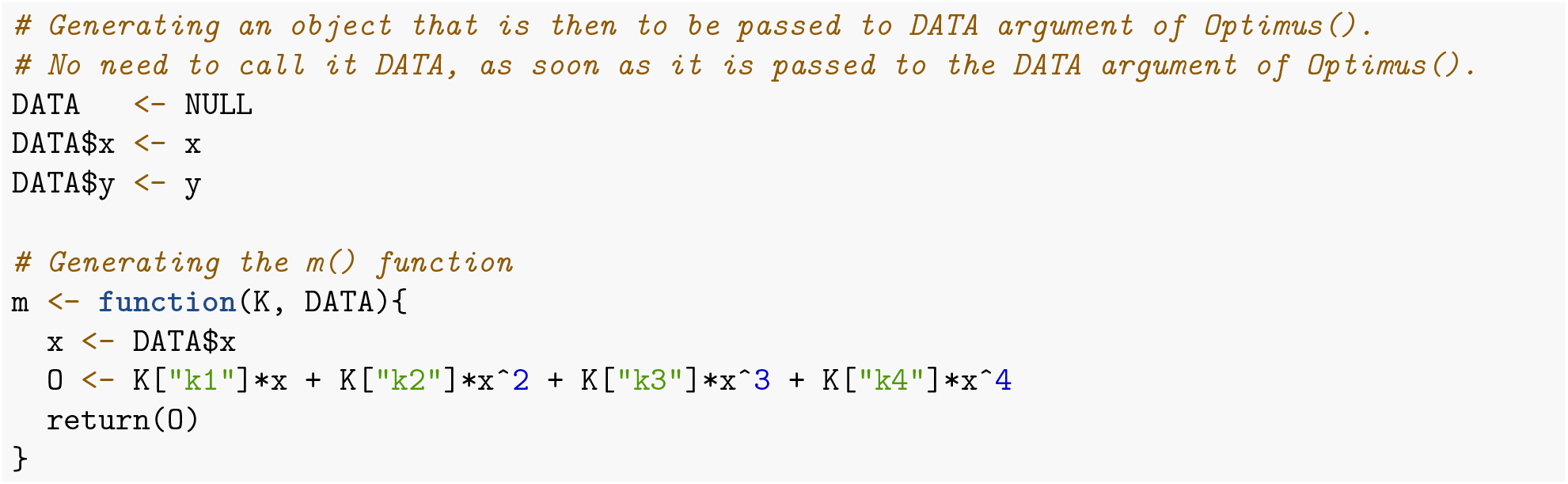

At this point, calling `m(K=K, DATA=DATA)` from within Optimus will return the predicted `O` set from the initial, non-optimal values for `K`, hence rather far from the target *O^trg^* = *y*.

In this example, the optimisation goal is for the `O` model outcomes to come as close as possible to the target observations *y*, to be achieved by optimising the coefficients `K`. The object *y* holding the target values therefore also needs to be specified and given as an input to the main `Optimus()` function (as a constituent entry in the `DATA` argument), just like *x* was supplied, as required, in this example, by the function `m()`.

Now, we need to define how the performance of a given snapshot of coefficients `K` is to be evaluated. For Optimus, this is done by specifying a function `u()`, which should **necessarily** take as inputs `O` (the output of `m()`) and the variable `DATA`. The output should have two components, `Q` holding a single number of the quality of the `K` coefficients, and `E` holding a (pseudo) energy for the given snapshot `K`. It is important that the returned (pseudo) energy value **must be lower for better performance/version** of `K`, never vice versa. The `Q` component of the `u()` function output is only used for plotting the optimisation process, and, if desired, can just repeat the value of the `E` component.

For our example, the `u()` function will assess the agreement between the snapshot of predictions `O` and the complete set of real observables (target) *y*. Here, we can use RMSD between `O` and *y* as a measure of `K` snapshot quality (`Q`). Since better agreement means better RMSD, it can be directly used as a pseudo energy (`E`), without putting a negative sign or performing some other mathematical operation on `Q`.

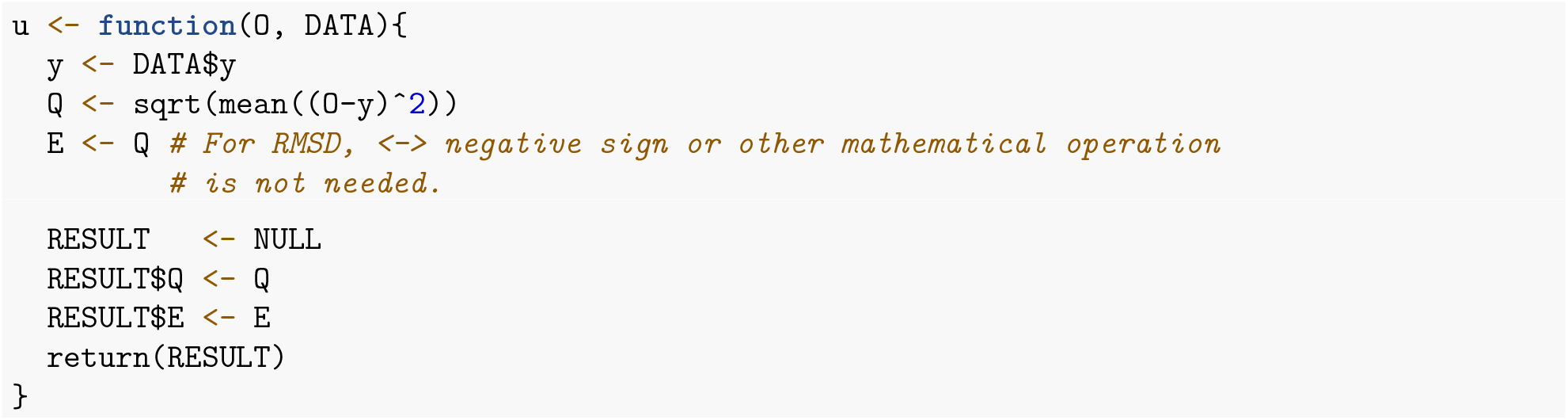

And finally, we need to define the rule, by which the `K` coefficient vector is to be altered from one step to another. This is done by defining a rule function `r()` that **must take** `K`, and **return an object analogous to** `K`, but with some alteration(s). In this example, for each snapshot of `K`, we shall randomly select one of its four coefficients, then either increment or decrement (chosen randomly) it by 0.0005, returning the altered set of coefficients.

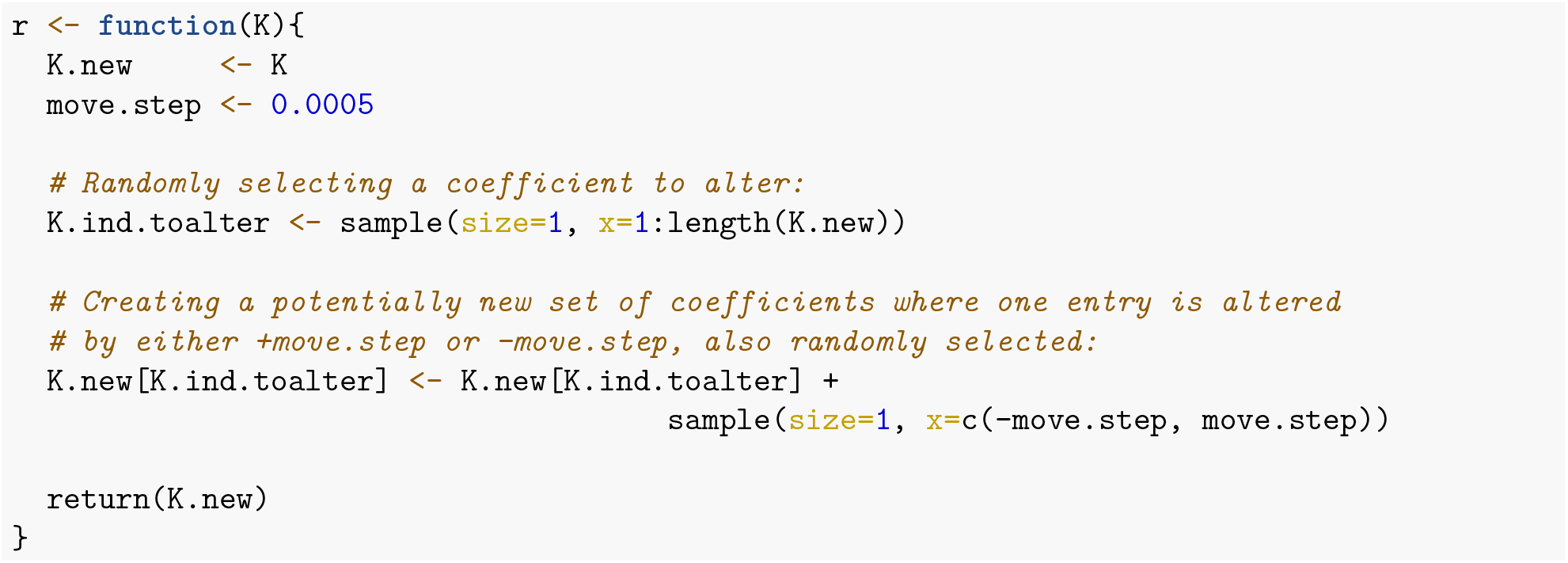

All the constructed objects (`K`) and functions (`m, u, r`), as well as the data required by `m()` and `u()` (stored in the variable to be passed to `DATA`) should be defined in an R session and given to Optimus as inputs. The users are free to define some dependencies as additional files (for example: initial protein geometry for a Monte-Carlo optimisation), which should be called from within the function definitions.

#### Acceptance Ratio Simulated Annealing Optimus Run

Having constructed `K`, dependent data for `DATA` argument of `Optimus()`, `m()`, `u()` and `r()`, we are now ready to call Optimus. Let us first investigate the Acceptance Ratio Annealing (SA) version of Optimus on 4 CPUs (the vast majority of personal computers currently have at least 4 CPUs), which can be executed as follows:

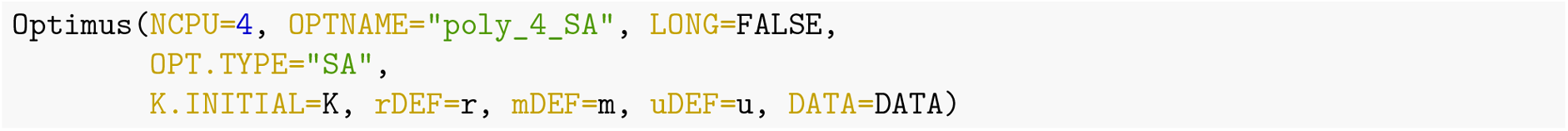

Note that the field `LONG=FALSE` is included in the function call so that all data from the optimsation process is saved. Calling Optimus with `LONG=TRUE` will result in a memory saving optimisation process (more details in the Advanced Usage section in this document). Of the 4 optimisation replicas, the second and fourth CPUs found the best parameter configuration (lowest RMSD) in our trial:

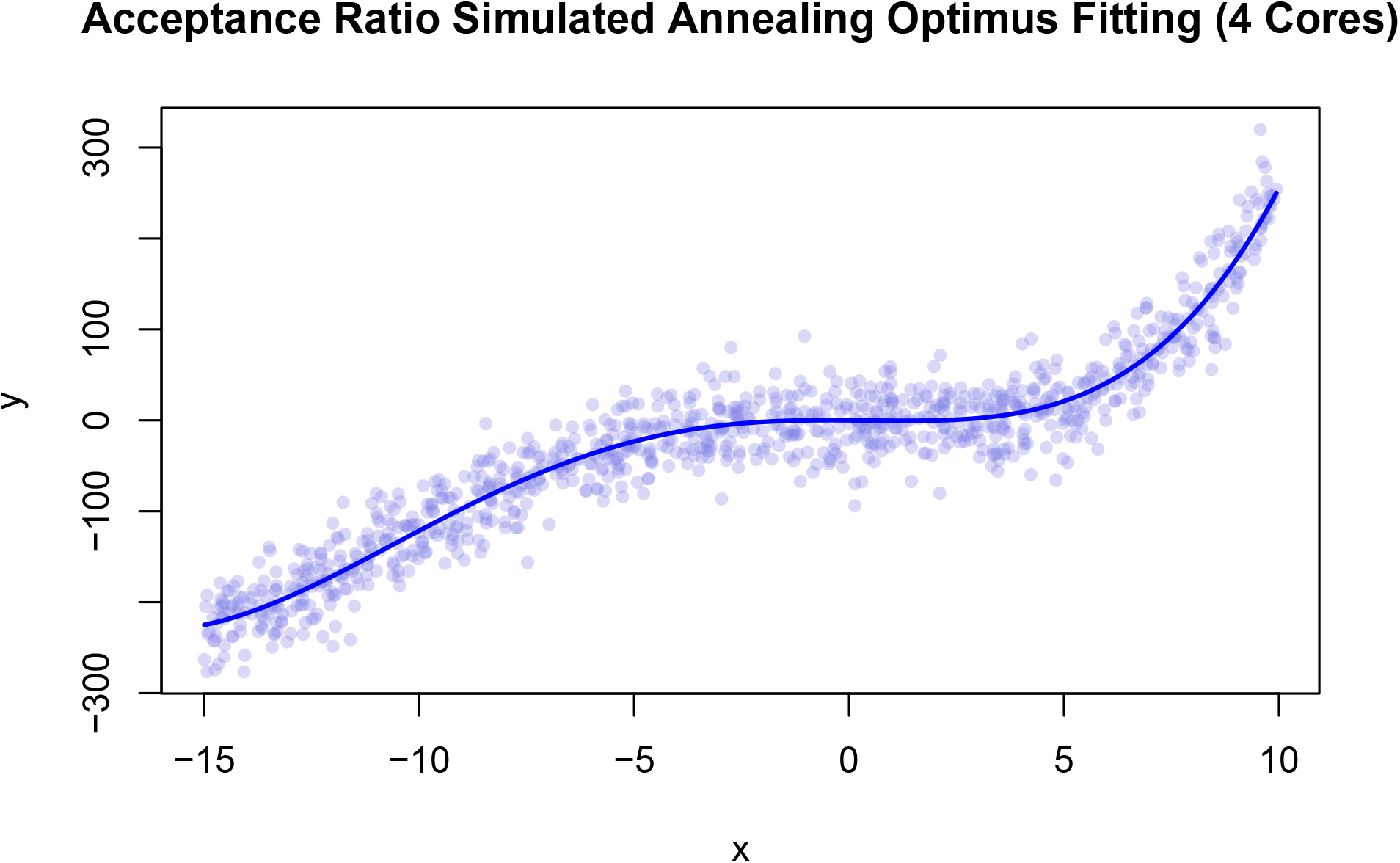

**Table 1:** 4-core Acceptance Ratio Simulated Annealing run results from Optimus.

|  | E (RMSD) | K1 | K2 | K3 | K4 |
| --- | --- | --- | --- | --- | --- |
| CPU 1 | 28.857 | -0.1560 | -0.2850 | 0.1905 | 0.0095 |
| CPU 2 | 28.841 | -0.3760 | -0.2825 | 0.1920 | 0.0095 |
| CPU 3 | 28.864 | -0.1045 | -0.2820 | 0.1905 | 0.0095 |
| CPU 4 | 28.841 | -0.3760 | -0.2825 | 0.1920 | 0.0095 |

The equation recovered by CPU 2 (and 4) is *y* = –0.3760*x* – 0.2825*x*^2^ + 0.1920*x*^3^ + 0.0095*x*^4^.

Notice that although the RMSD of this solution, 28.841, is greater than the RMSD of the least squares solution, 28.827, it is less than the RMSD of the de-noised data found above, 28.859.

The two graphs below illustrate i) the evolution of the system pseudo temperature, in response to alterations made by the Temperature Control Unit (TCU), as a function of the optimsation step; and ii) the observed acceptance ratio as a function of the optimisation step, respectively. The graphs show data from the last 20 000 steps of the optimisation executed by CPU 2.

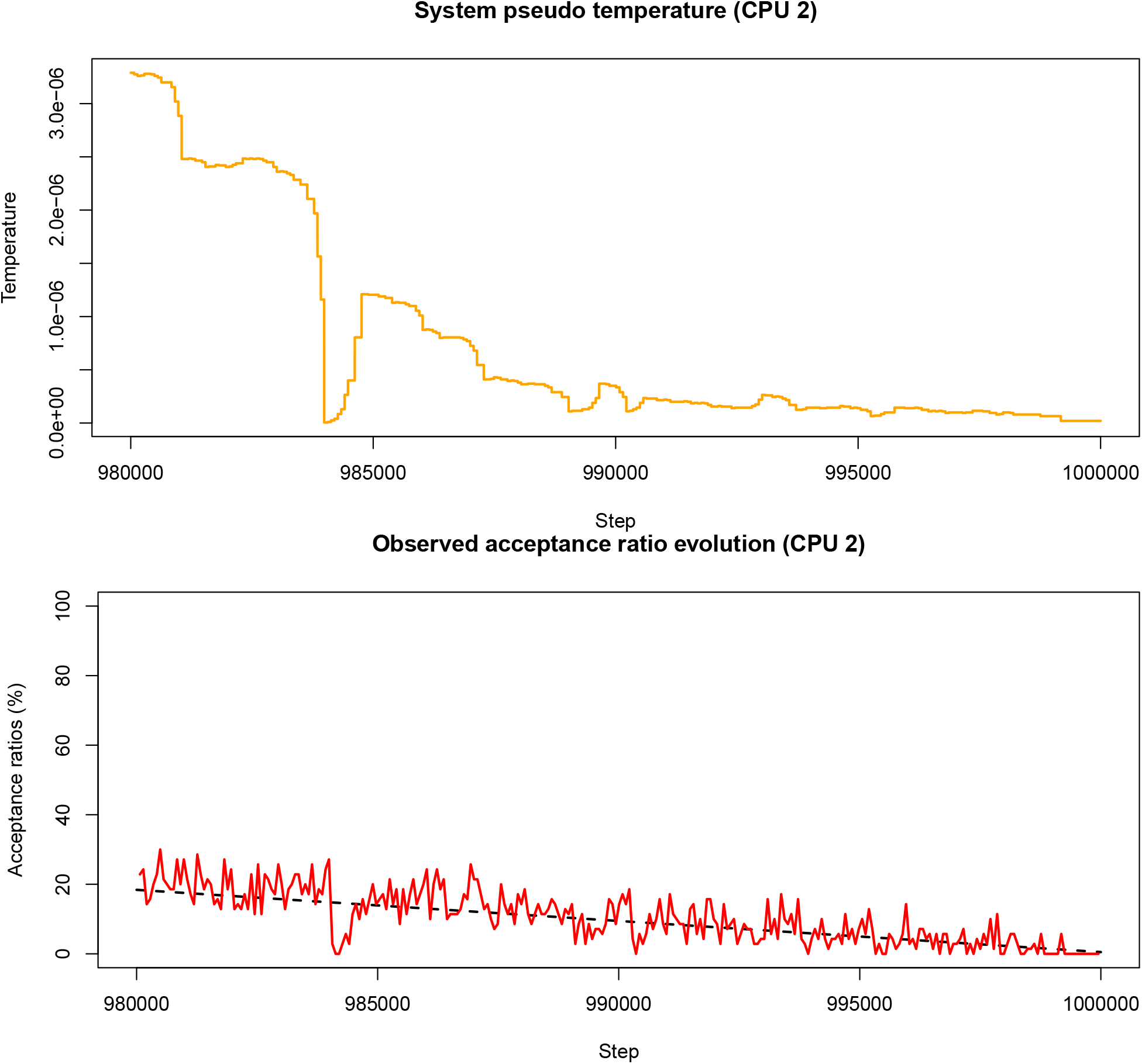

In the first plot, the solid red line tracks the observed acceptance ratios calculated by Optimus at the end of each `STATWINDOW` and the dashed black line tracks the target acceptance ratio based on the annealing schedule. From the above two graphs, notice that while the observed acceptance ratio tracks the target acceptance ratio closely, the system pseudo temperature changes significantly and non-monotonically. This illustrates that the adaptive thermoregulation allows Optimus to effectively anneal the system acceptance ratio.

#### Acceptance Ratio Replica Exchange Optimus Run

Let us now consider the Replica Exchange version of Optimus on 12 CPUs. The purpose here is to illustrate how to run an optimisation using the Replica Exchange version of Optimus; this method is of course an overkill for solving this simple task.

In addition to the arguments specified above, the Replica Exchange version of Optimus also requires an input variable `ACCRATIO`, which is a vector that defines the acceptance ratios to be used for each of the replicas initiated, 12 in this case. Note that the length of `ACCRATIO` must always be equal to the argument `NCPU`.

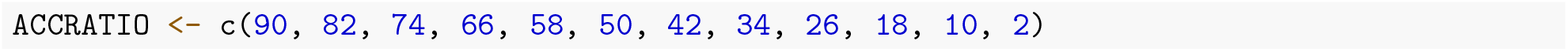

Having defined the acceptance ratios for each level, the optimisation can be executed as follows:

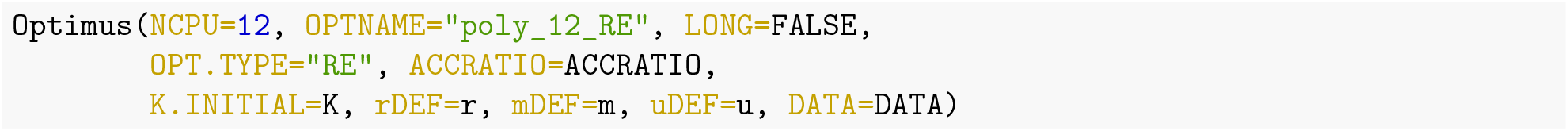

Of the 12 optimisation replicas, replica 8 finds the best parameter configuration (lowest RMSD) in this trial:

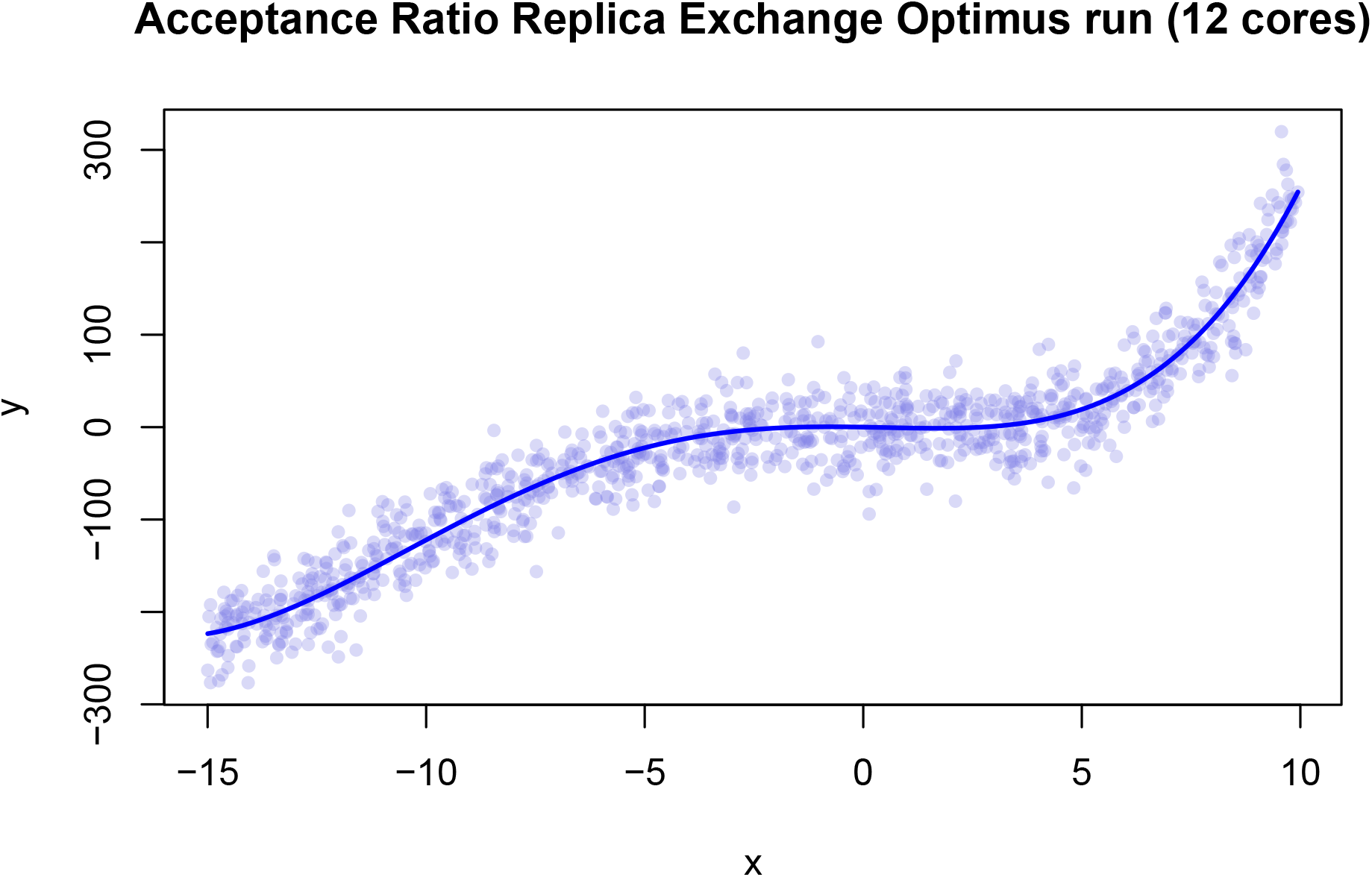

**Table 2:** 12-core Replica Exchange run results from Optimus.

|  | Replica Acceptance Ratio | E (RMSD) | K1 | K2 | K3 | K4 |
| --- | --- | --- | --- | --- | --- | --- |
| CPU 1 | 90 | 29.71934 | -3.2760 | -0.5760 | 0.2395 | 0.0135 |
| CPU 2 | 82 | 29.51819 | -3.4445 | -0.3755 | 0.2300 | 0.0115 |
| CPU 3 | 74 | 28.88707 | -0.0340 | -0.2375 | 0.1870 | 0.0090 |
| CPU 4 | 66 | 30.10499 | -3.8095 | -0.6630 | 0.2435 | 0.0140 |
| CPU 5 | 58 | 29.06293 | -2.3575 | -0.4180 | 0.2200 | 0.0115 |
| CPU 6 | 50 | 29.70906 | -3.7360 | -0.3785 | 0.2320 | 0.0115 |
| CPU 7 | 42 | 28.85095 | -1.1370 | -0.3515 | 0.2045 | 0.0105 |
| CPU 8 | 34 | 28.82721 | -0.8175 | -0.3130 | 0.1990 | 0.0100 |
| CPU 9 | 26 | 28.84057 | -0.5760 | -0.2740 | 0.1940 | 0.0095 |
| CPU 10 | 18 | 29.48785 | -2.7805 | -0.5420 | 0.2325 | 0.0130 |
| CPU 11 | 10 | 28.85095 | -1.1370 | -0.3515 | 0.2045 | 0.0105 |
| CPU 12 | 2 | 29.41377 | 1.2255 | -0.0895 | 0.1645 | 0.0070 |

The equation recovered by CPU 8 is *y* = –0.8175*x* – 0.313*x*^2^ + 0.199*x*^3^ + 0.01*x*^4^.

Notice that the RMSD of this solution, 28.8272, is less than the RMSD of the Acceptance Ratio Simulated Annealing solution, 28.841, and only slightly greater than the RMSD of the least squares solution, 28.8266.

Let us now briefly examine the evolution of the system pseudo temperature and acceptance ratio compliance in response to the adaptive thermoregulation. The following two graphs represent data from the last 20 000 steps of optimisation replica running on CPU 8 (fixed 34% target acceptance ratio).

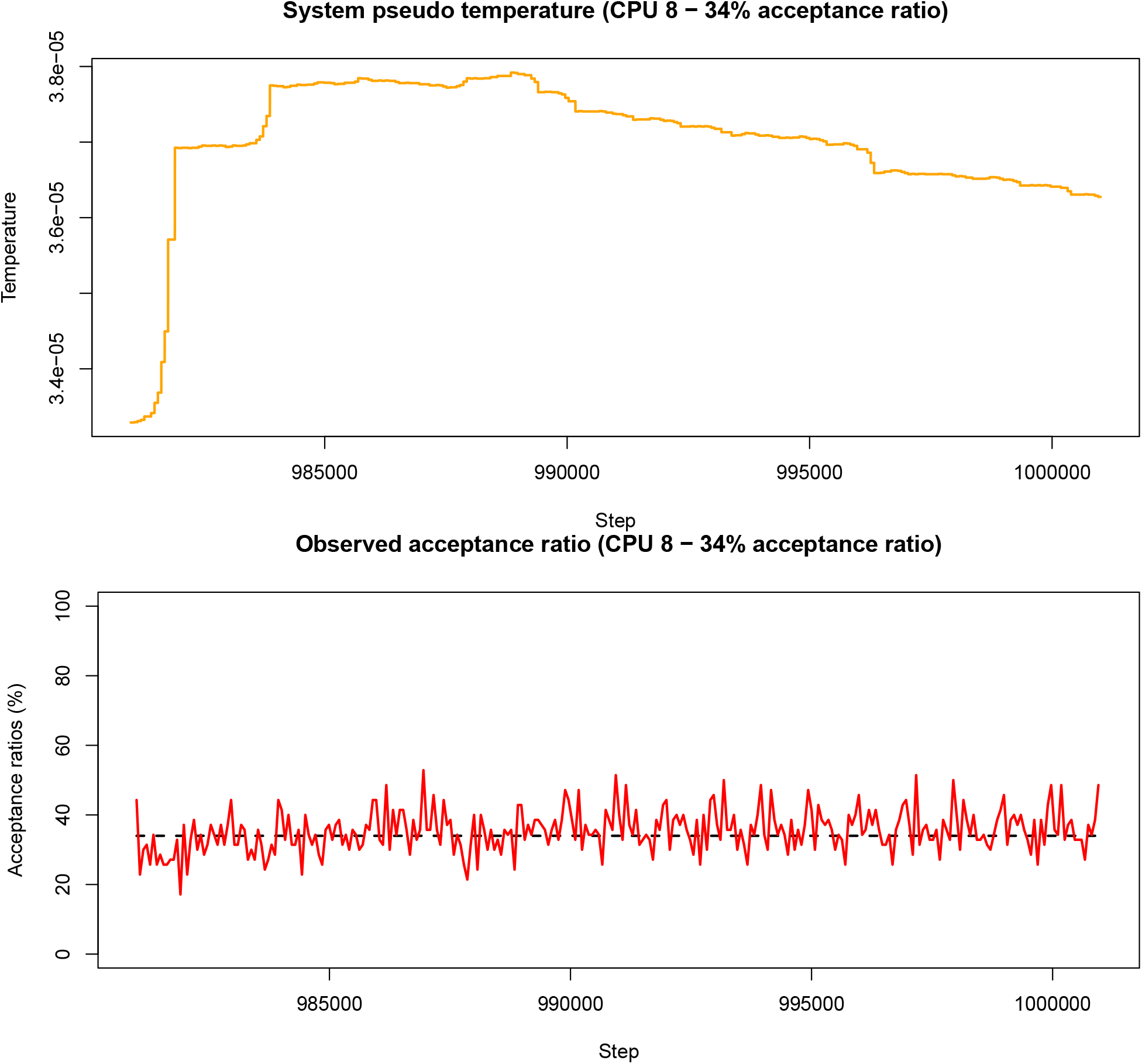

Notice that in the observed acceptance ratio graph, the dashed line indicating the target acceptance ratio is constant (as opposed to linearly changing as in acceptance ratio annealing). This is because each processor in the replica exchange mode has a single target acceptance ratio, as described above. Here again, adaptive decisions on the pseudo temperature to maintain the desired acceptance ratio result in non-monotonic, and non-uniform pseudo temperature adjustments, while the observed acceptance ratios fluctuate relatively closely around the set target value.

#### Summary

We now understand the input requirements to interface with the Acceptance Ratio Simulated Annealing and Replica Exchange versions of Optimus. In this example, both versions retrieved solutions having a lower RMSD than the de-noised data, and only a slightly greater RMSD than the optimal least squares solution. Replica Exchange resulted in a better solution than Simulated Annealing, at the cost of greater computing resources.

**Table 3:** Summary of solutions.

|  | E (RMSD) | K1 | K2 | K3 | K4 |
| --- | --- | --- | --- | --- | --- |
| De-noised Function | 28.85858 | -1.0000 | -0.3000 | 0.2000 | 0.0100 |
| Optimus (AR Simulated Annealing) | 28.84100 | -0.3760 | -0.2825 | 0.1920 | 0.0095 |
| Optimus (AR Replica Exchange) | 28.82721 | -0.8175 | -0.3130 | 0.1990 | 0.0100 |
| Least Squares | 28.82655 | -0.7406 | -0.3074 | 0.1978 | 0.0099 |

### Tutorial 2: Finding an Optimal Equation by Navigating Through a Term Space

#### Problem Statement

Consider again the synthetic data that was created in **Tutorial 1**. Suppose that we were only provided with the data and, unlike in **Tutorial 1**, had no knowledge of the best terms to be included in a functional representation of said data. In this example, we shall use Optimus to determine which terms should be used in a least squares fitting of the data to achieve a representation with low RMSD while avoiding overfitting the data with too complicated equation.

Let us start by generating the same data which was used in **Tutorial 1**:

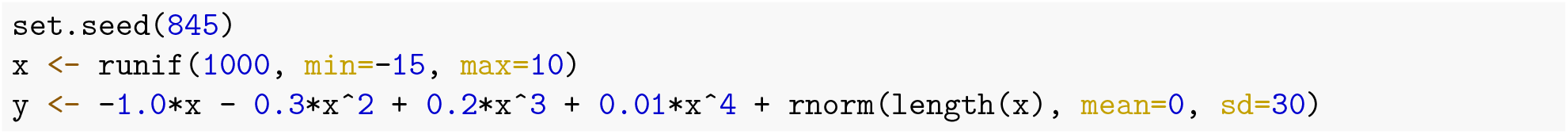

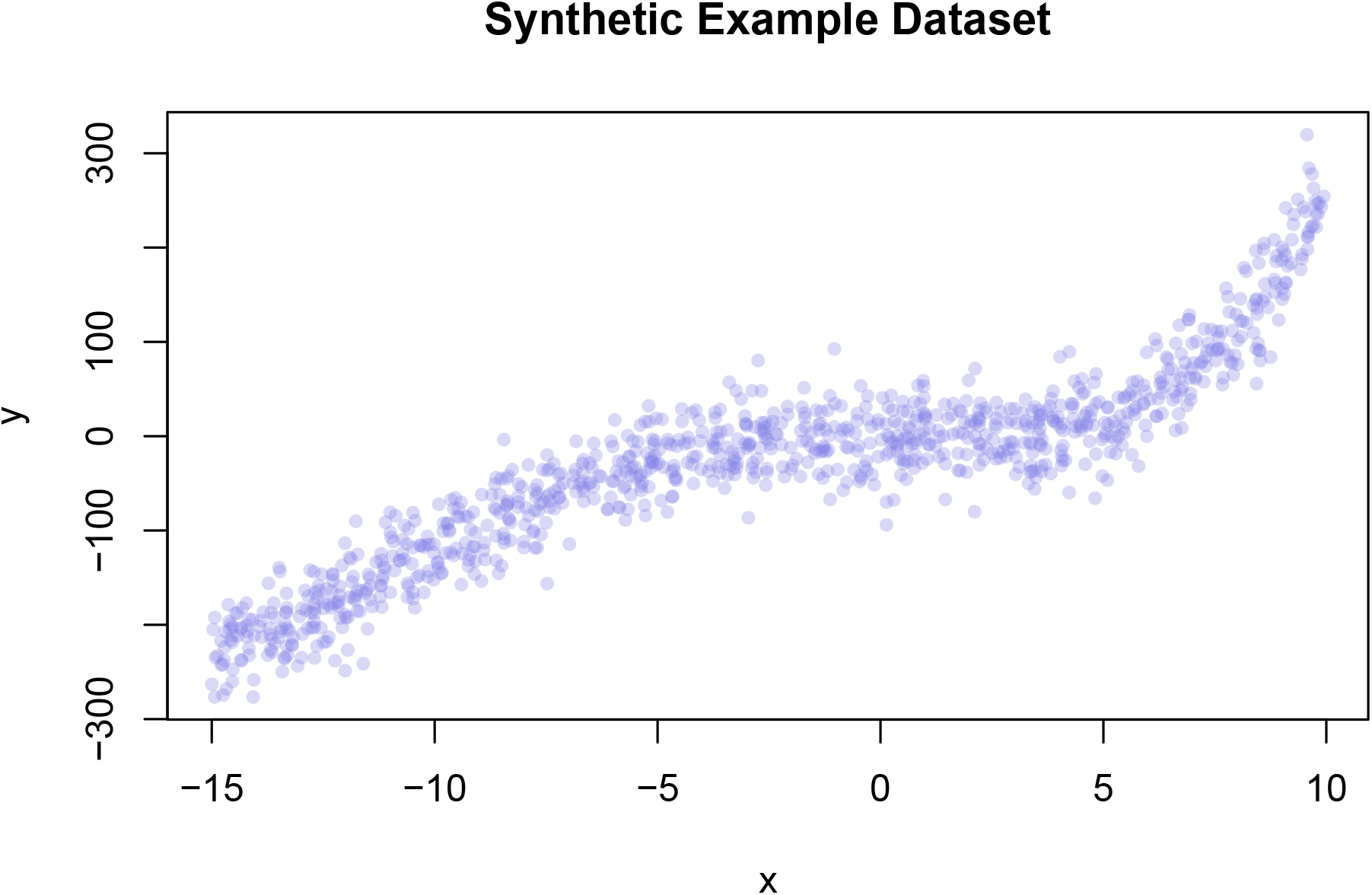

From **Tutorial 1**, we know that if presented with this data and under the assumption that the most appropriate model to describe the data is *k*_1_*x* + *k*_2_*x*^2^ + *k*_3_*x*^3^ + *k*_4_*x*^4^, the Least Squares model fitting is *y* = –0.741*x* – 0.307*x*^2^ + 0.198*x*^3^ + 0.010*x*^4^. We also know that the RMSD between the observed data *y* and the linear model fitting outcome is:

~~~
## [1] 28.82655
~~~

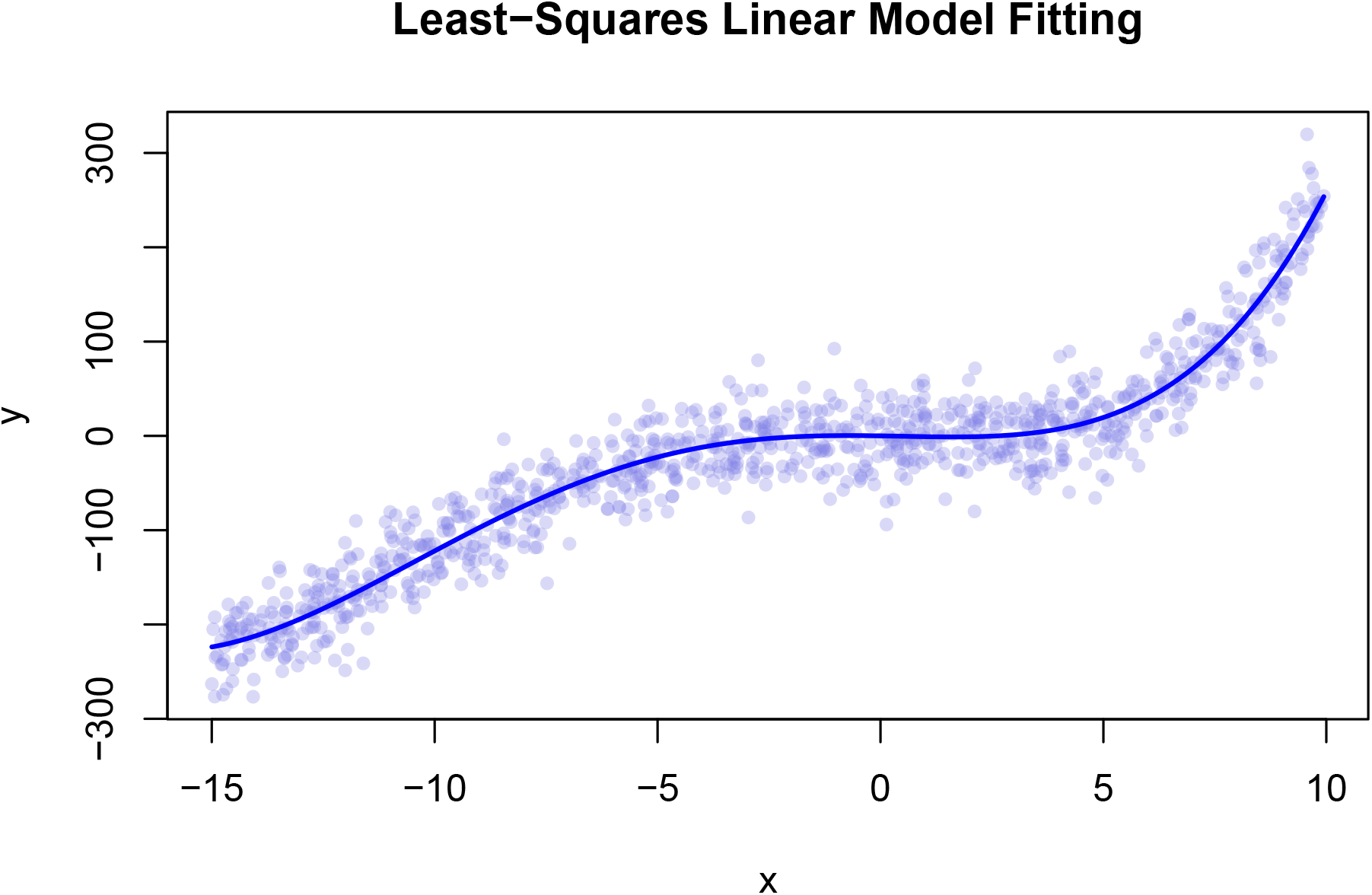

#### Defining Optimus Inputs

Let us first define an ordered set *terms* that is a collection of candidate terms to include in the representation of the data:

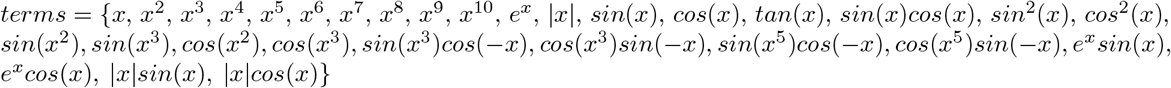

Let *terms_i_* denote the *i^th^* term in the set *terms* (for example, *terms*_14_ = *cos*(*x*)). We shall use the model:

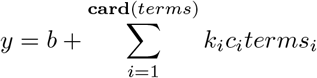

where each *k_i_* is a binary variable (meaning a variable taking a value of either 0 or 1) indicating whether the *i^th^* term is included in the representation, each *c_i_* is a non-zero coefficient for the *i^th^* term and *b* is a real number (the intercept). In our case, **card**(*terms*) = 30 so explicitly, our model is:

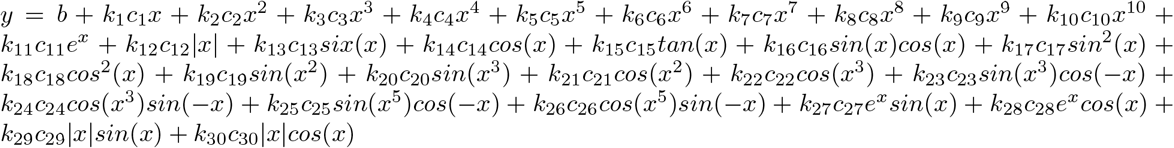

Formally, `K` will be a numeric vector of length **card**(*terms*) whose *i^th^* entry is *k_i_*. `K` uniquely specifies a set *activeTerms* = {*terms_i_* ∀ *i* |*k_i_* = 1}. Note that *activeTerms* ⊆ *terms*. Each binary variable *k_i_* should be initialized randomly as below:

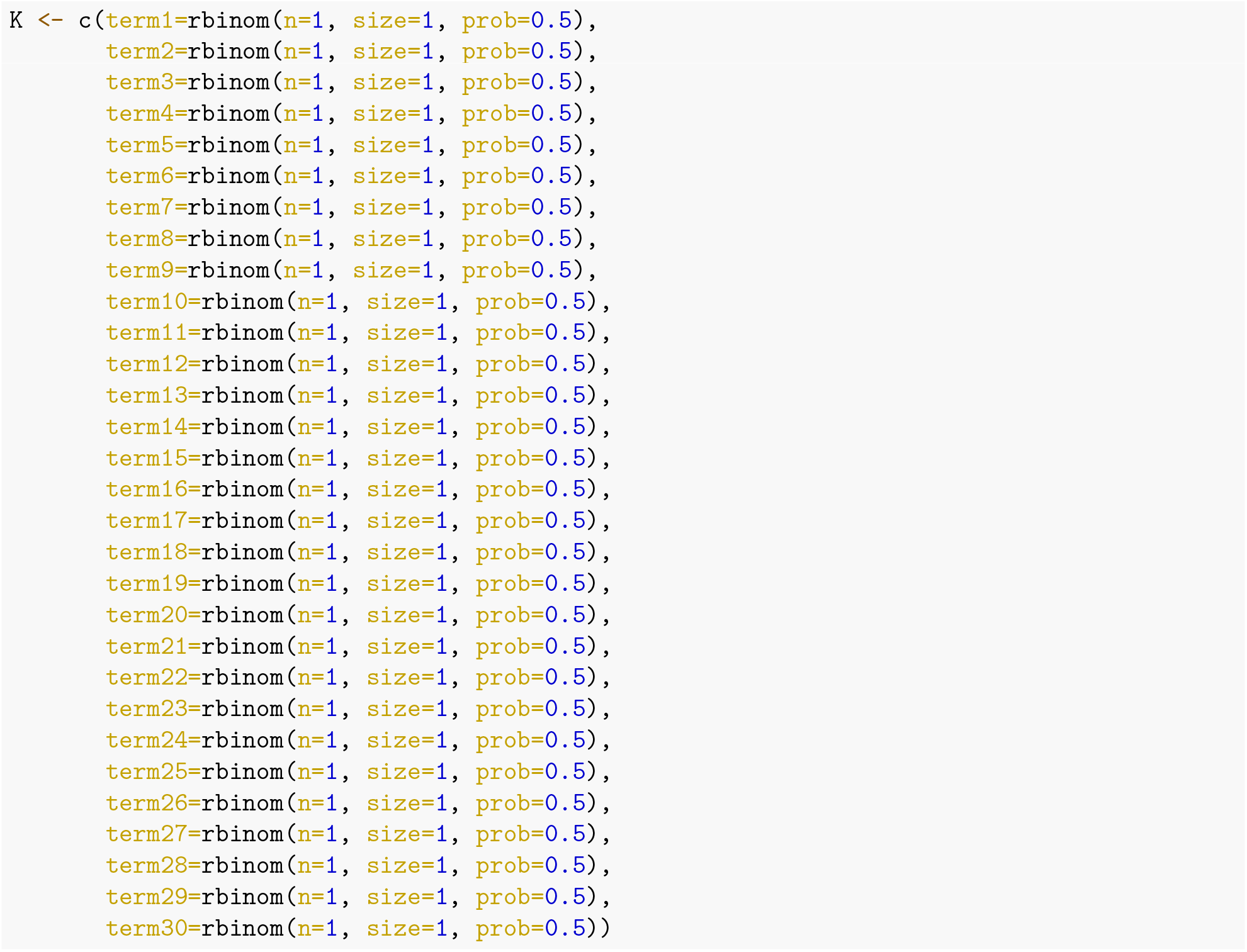

Next, we must define the model function `m()` that will operate on the parameter snapshot `K` and return an observable object `O`. For a given set *activeTerms* specified by `K`, `m()` will fit a linear model to the data using the entries in *activeTerms* and using the built in generalised linear model (`glm()`) function in R, thereby determining values for the variables *c_i_* and *b*. Accordingly, `m()` will require access to the variables *x* and *y*, which will be provided as entries in `DATA`, a variable of type list, as in **Tutorial 1**. The object `O` will be the corresponding output of the function `glm()`. In the case that the set *activeTerms* is the empty set (meaning that all entries in `K` are 0), `m()` will fit a model using the relationship *y*~*x*.

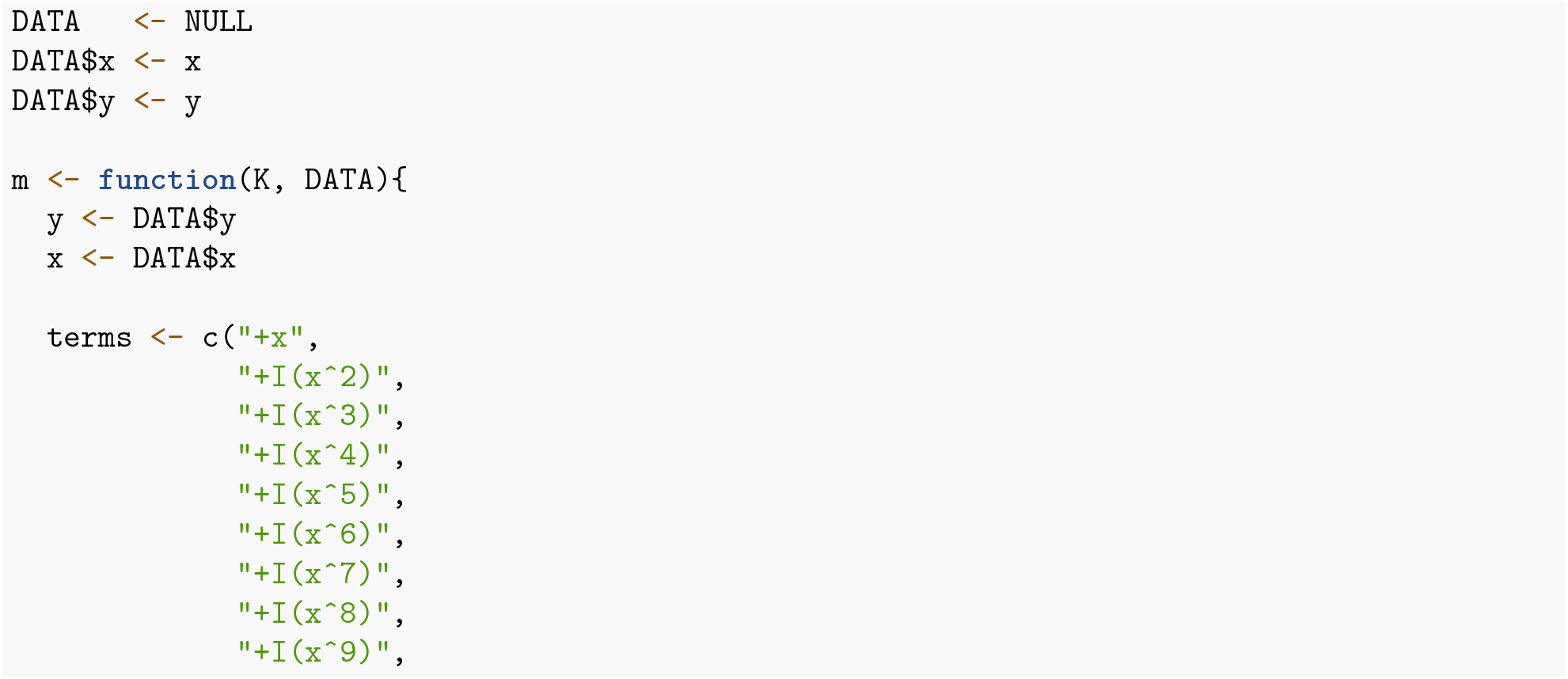
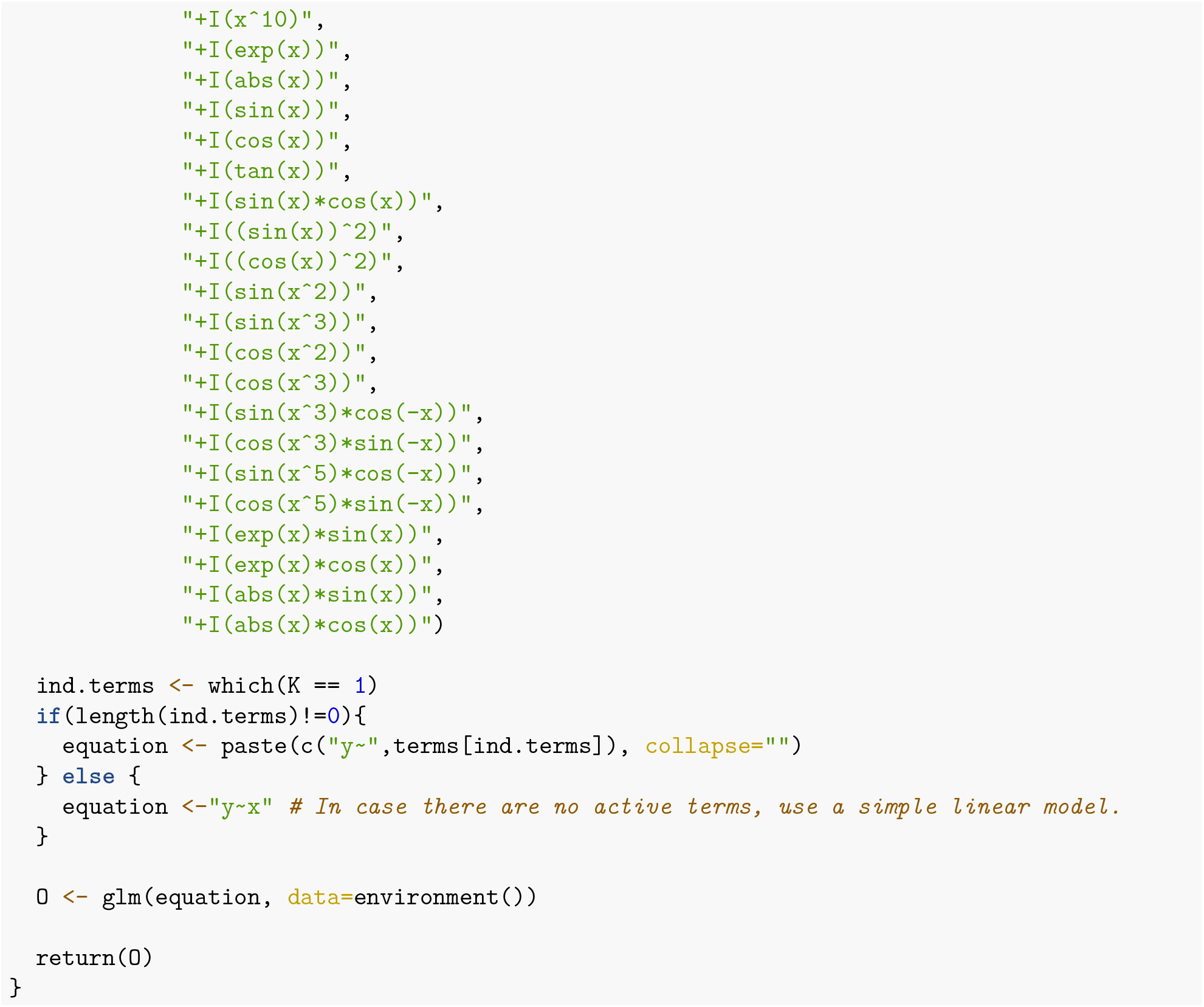

Having defined `m`, we can now proceed to define the function `u`, which will determine how well a given configuration of parameters `K` is performing by operating on the observable object `O` outputted by `m()` and on the variable `DATA`. Here, to quantify (and thus be able to compare) the desirability of a given model for the data, we will employ the Akaike Information Criterion (*AIC*) from information theory, defined as follows:

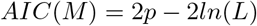

where *p* is the number of parameters in the fitted model, *L* is the maximum likelihood of the model *M* given the data.

The target representation will be the fitted model *M* (whose terms are elements of *terms*) that minimises the *AIC*. It is important to note that the 2*p* term in the *AIC* penalises overfitting by increasing *AIC* as function of the number of parameters, while the −2*ln*(*L*) term rewards models that better represent the data by decreasing *AIC* as a function of the likelihood of the model.

As articulated in **Tutorial 1**, the output of `u()` should have a component `E` holding a pseudo energy for the parameter snapshot `K`, and a component `Q` that can be used for plotting the optimisation process. In this case, `E` will be equal to the value of *AIC* (implemented using the built in `AIC()` function in R) and `Q` will be equal to the RMSD between the predicted values of *y* from the fitted model and the actual *y* values, used for plotting purpose. Consequently, `u()` will need access to the variable *y*. The definition of `u()` is below:

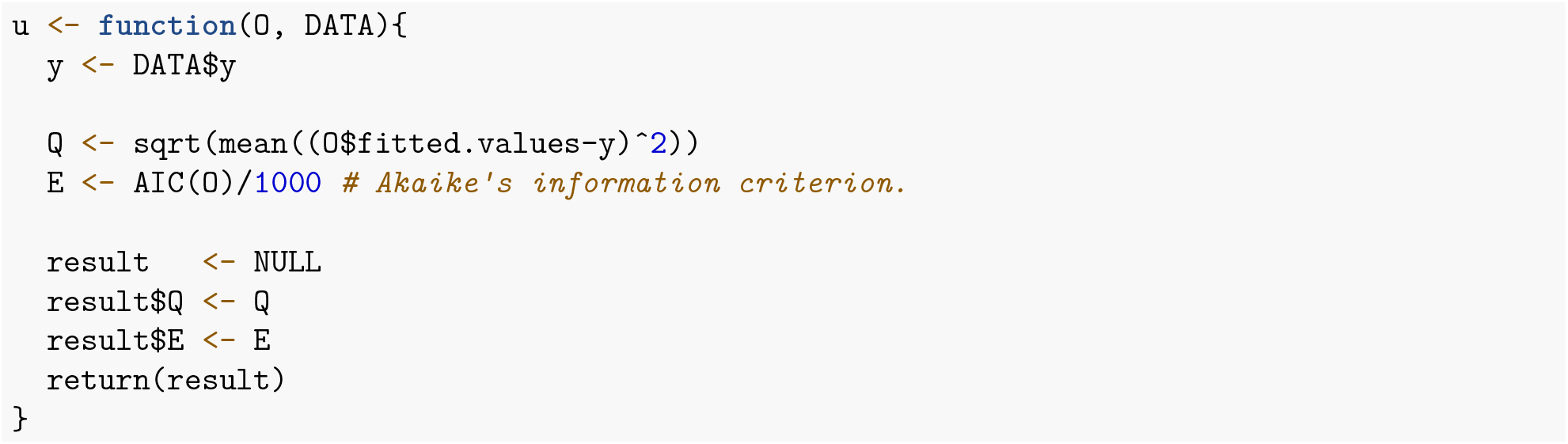

Finally, we need to define the rule function `r()`. We will adopt the following simple procedure: randomly select an equation entry from `K` and switch its value to the other binary value (on to off, or off to on).

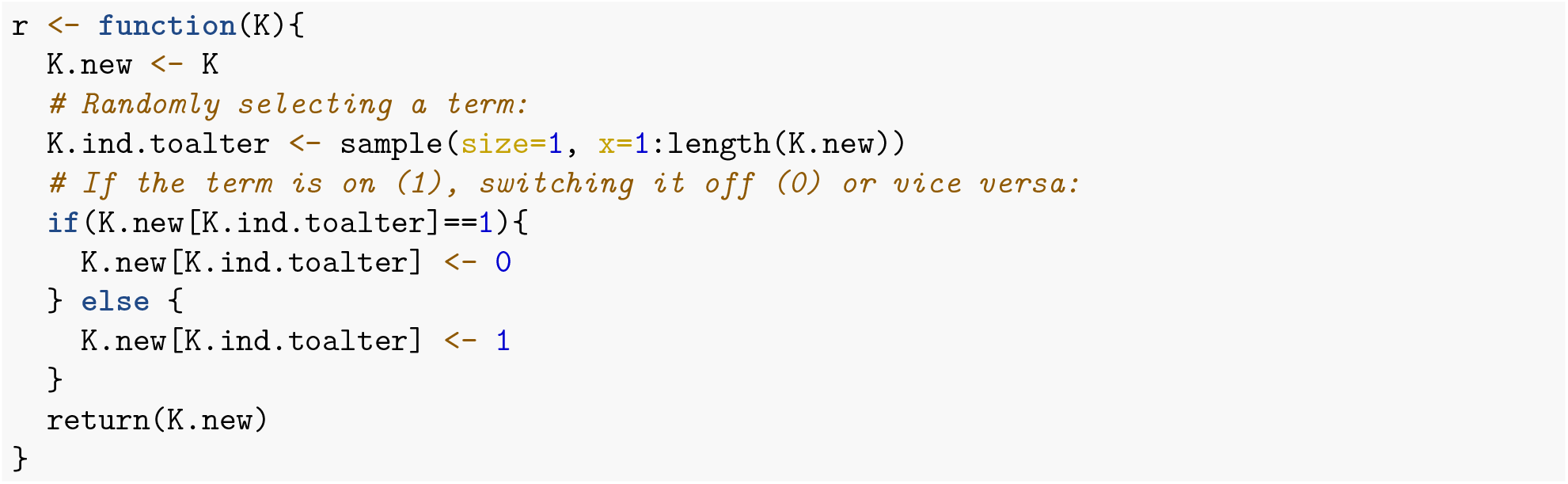

Having defined all the necessary inputs, we are now ready to call Optimus.

An important remark is that modelling this problem in this manner results in an objective function (*AIC*) that is not smooth because small changes in the parameter set `K` (as defined by `r()`) can produce significantly large changes in the objective value. The equation, i.e. the system itself, changes from one step to another. Therefore, an entirely different model is being used to fit the data at each step. Optimisation procedures in such cases are in danger to be trapped in certain minima due to a major change in pseudo temperatures necessary to overcome barriers while the system abruptly evolves. Despite this, we will see that Optimus will get the job done by arriving at good solutions largely as a consequence of its adaptive thermoregulation and acceptance-ratio-guided optimisation procedure.

#### Acceptance Ratio Simulated Annealing Optimus Run

In addition to the inputs defined above, Optimus can optionally take other inputs to dictate the optimisation process (see the Advanced User Manual), all of which have built in default values and some of which will be altered in this example due to the increased computational complexity of the model defined in this tutorial compared to that of **Tutorial 1**. The variable `NUMITER` represents the number of iterations of the optimisation process (per core) and has a default value of 1 000 000. For this example, 200 000 iterations will be used to reduce the running time of Optimus given that each iteration is more computationally demanding than in **Tutorial 1**. The variable `CYCLES` (unique to Acceptance Ratio Annealing Optimus runs) denotes the number of acceptance ratio annealing cycles. Its default value is 10, however it will be set to 2 in this example so that each annealing cycle has 100 000 iterations just as in **Tutorial 1** (the number of steps per cycle is calculated as `NUMITER`/`CYCLES`). Lastly, the variable `DUMP.FREQ`, the frequency (in steps) with which the best found model is assessed and outputted by the function, will be set to 100 000 (its default value is 10 000).

Let us again investigate the Simulated Annealing (SA) version of Optimus on 4 processors, which can be executed as follows:

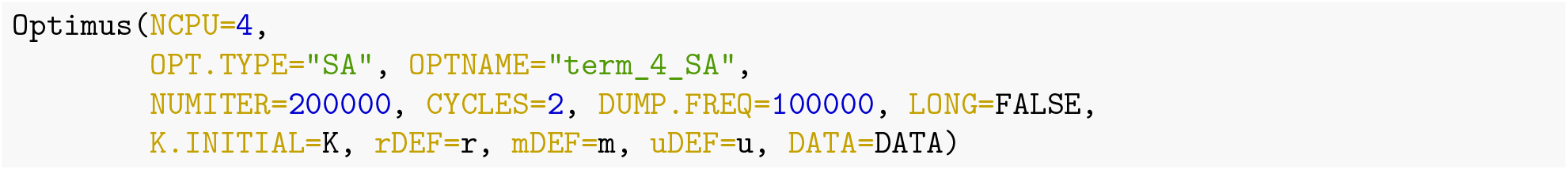

Interestingly, each of the 4 computing cores arrive at the same solution in this instance.

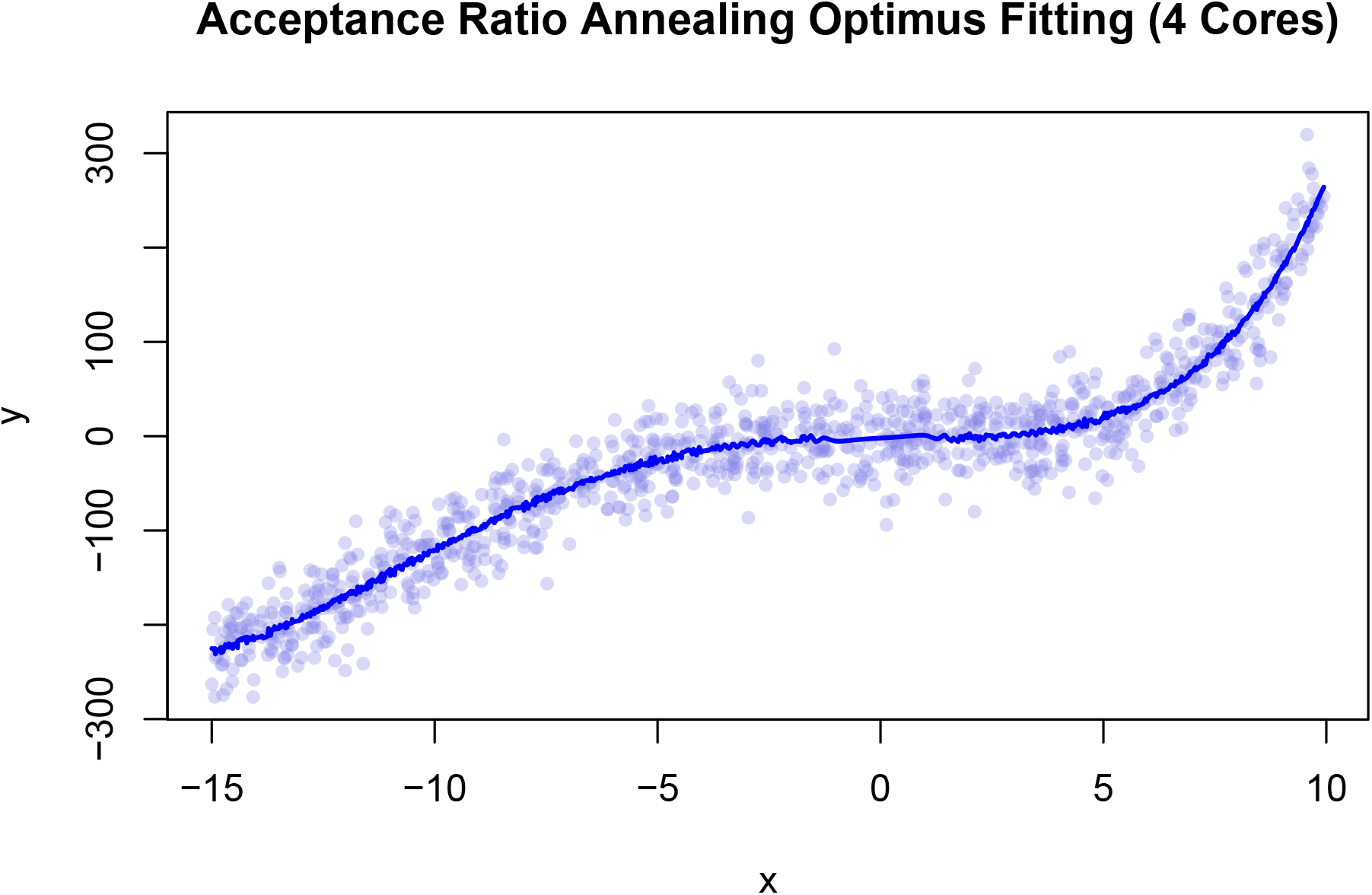

**Table 4:** 4-core Acceptance Ratio Simulated Annealing results from Optimus.

|  | CPU 1 | CPU 2 | CPU 3 | CPU 4 |
| --- | --- | --- | --- | --- |
| E (AIC) | 9.567 | 9.567 | 9.567 | 9.567 |
| Q (RMSD) | 28.693 | 28.693 | 28.693 | 28.693 |
| Term 1 | 0.000 | 0.000 | 0.000 | 0.000 |
| Term 2 | 1.000 | 1.000 | 1.000 | 1.000 |
| Term 3 | 1.000 | 1.000 | 1.000 | 1.000 |
| Term 4 | 1.000 | 1.000 | 1.000 | 1.000 |
| Term 5 | 0.000 | 0.000 | 0.000 | 0.000 |
| Term 6 | 0.000 | 0.000 | 0.000 | 0.000 |
| Term 7 | 0.000 | 0.000 | 0.000 | 0.000 |
| Term 8 | 0.000 | 0.000 | 0.000 | 0.000 |
| Term 9 | 0.000 | 0.000 | 0.000 | 0.000 |
| Term 10 | 0.000 | 0.000 | 0.000 | 0.000 |
| Term 11 | 1.000 | 1.000 | 1.000 | 1.000 |
| Term 12 | 0.000 | 0.000 | 0.000 | 0.000 |
| Term 13 | 0.000 | 0.000 | 0.000 | 0.000 |
| Term 14 | 0.000 | 0.000 | 0.000 | 0.000 |
| Term 15 | 0.000 | 0.000 | 0.000 | 0.000 |
| Term 16 | 0.000 | 0.000 | 0.000 | 0.000 |
| Term 17 | 0.000 | 0.000 | 0.000 | 0.000 |
| Term 18 | 0.000 | 0.000 | 0.000 | 0.000 |
| Term 19 | 0.000 | 0.000 | 0.000 | 0.000 |
| Term 20 | 1.000 | 1.000 | 1.000 | 1.000 |
| Term 21 | 0.000 | 0.000 | 0.000 | 0.000 |
| Term 22 | 0.000 | 0.000 | 0.000 | 0.000 |
| Term 23 | 0.000 | 0.000 | 0.000 | 0.000 |
| Term 24 | 0.000 | 0.000 | 0.000 | 0.000 |
| Term 25 | 0.000 | 0.000 | 0.000 | 0.000 |
| Term 26 | 1.000 | 1.000 | 1.000 | 1.000 |
| Term 27 | 0.000 | 0.000 | 0.000 | 0.000 |
| Term 28 | 0.000 | 0.000 | 0.000 | 0.000 |
| Term 29 | 0.000 | 0.000 | 0.000 | 0.000 |
| Term 30 | 0.000 | 0.000 | 0.000 | 0.000 |

Thus, the optimal functional representation found by Optimus has the following form:

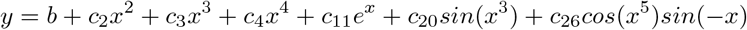

Below is the explicit representation after determining the coefficients *c_i_* and *b*:

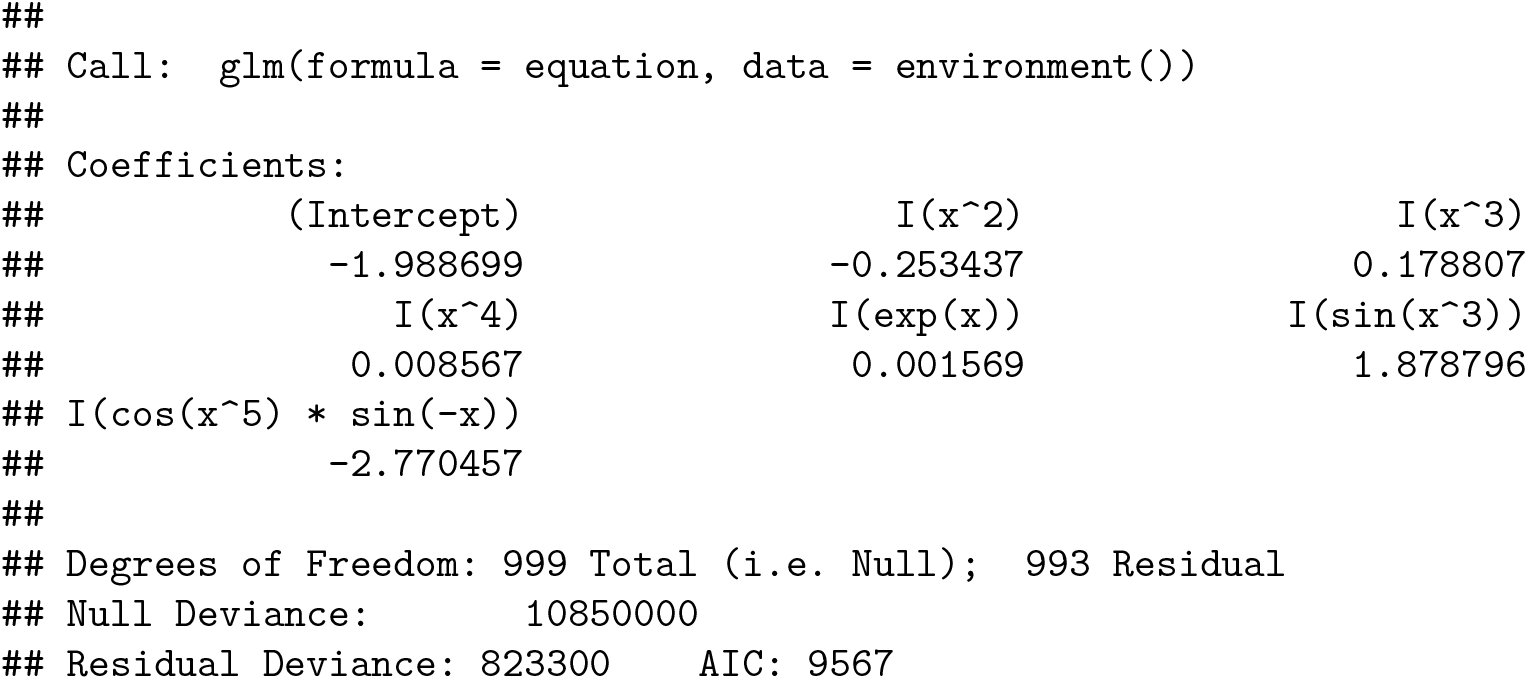

Notice that the solution selected by Optimus results in an RMSD of 28.693 which is lower than the RMSD of the Least Squares Solution (28.827) that assumes the appropriate model is *k*_1_*x* + *k*_2_*x*^2^ + *k*_3_*x*^3^ + *k*_4_*x*^4^. Optimus selected a model which does not include all terms from the form used to generate the data. If a user were concerned by the fact that the model Optimus selected contains more terms (6) than are used in the representation of the de-noised data (4), the user could either increase the multiplicative factor associated with the term *p* in the *AIC* to more strongly penalise representations involving a greater number of parameters. Alternatively, the user could also modify the function `r()` to ensure that only a fixed number of terms are ever active.

Let us now take a look at how the adaptive thermoregulation performed given this highly non-smooth objective. The graphs below should now feel very familiar, they represent data taken from the last 20 000 iterations of the optimisation protocol executed by CPU 1.

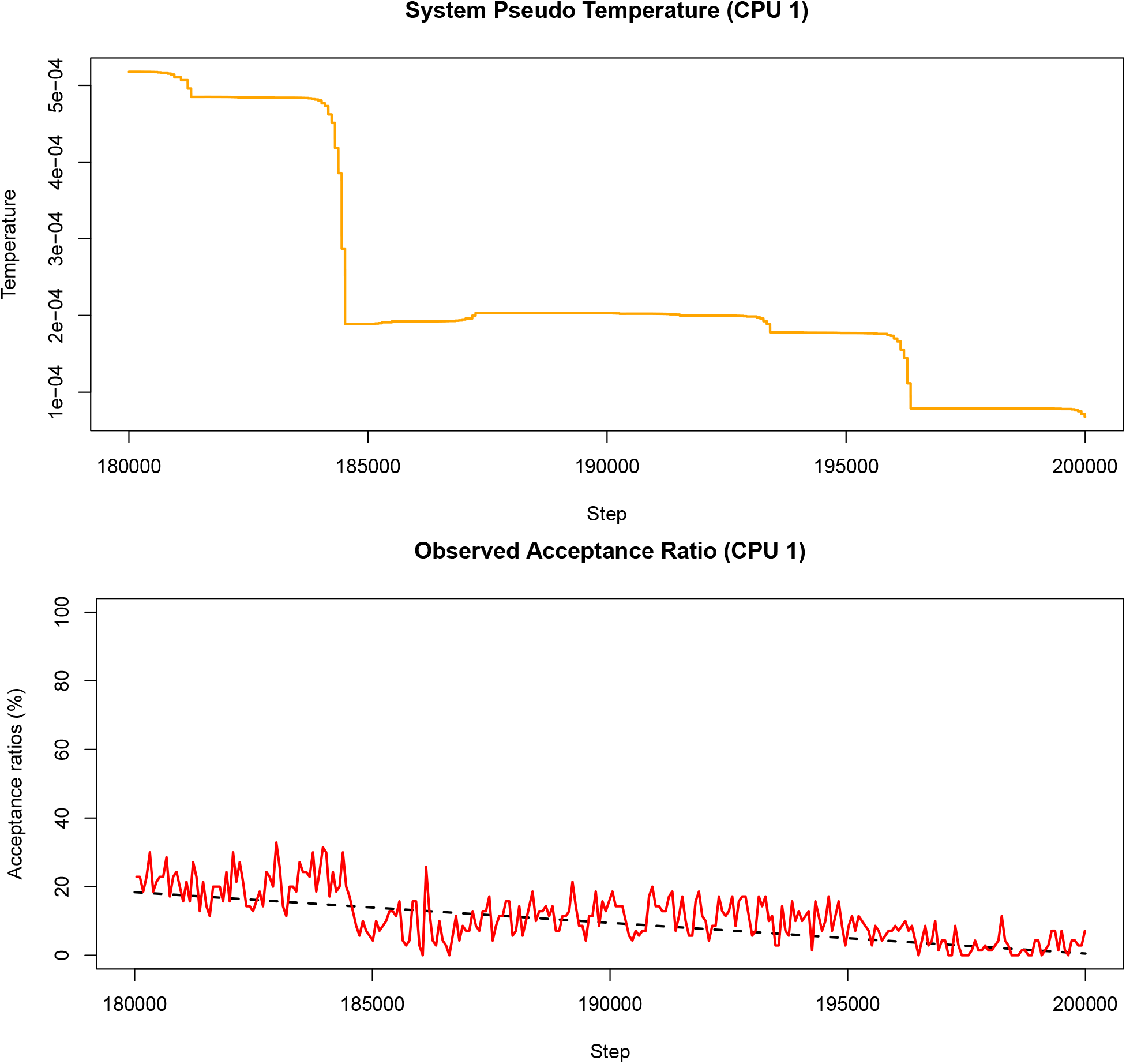

Despite optimising a completely different model with a non-smooth objective function, the TCU succeeds in dynamically adjusting the system pseudo-temperature such that the observed acceptance ratio follows the annealing schedule rather well.

#### Acceptance Ratio Replica Exchange Optimus Run

Let us now consider the Acceptance Ratio Replica Exchange version of Optimus on 12 CPUs with the variable `ACCRATIO` defined as in **Tutorial 1**.

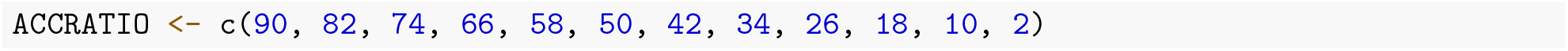

As in the Acceptance Ratio Simulated Annealing run above, we will again execute the optimisation procedure for 200 000 iterations. The Replica Exchange version of Optimus takes an input argument `EXCHANGE.FREQ` (default value 1000) which specifies the total number of exchanges that will occur during the optimisation process. Consequently, the number of optimisation iterations that occur between subsequent exchanges between replicas can be calculated as `NUMITER`/`EXCHANGE.FREQ`, which is 200 iterations in this case.

Here we will set the input parameter `STATWINDOW` to have value 50 (its default value is 70). This signifies that the TCU will update the system pseudo temperature once every 50 iterations on each optimisation replica. This guarantees that 4 temperature adjustments will be made if a given replica is involved in two subsequent exchanges (because the number of iterations between exchanges is 200, as explained in the preceding paragraph) as opposed to merely 2 adjustments which would be the case if `STATWINDOW` were left to take its default value and could result in poor agreement between the observed acceptance ratio of the replica in question and the target acceptance ratio. The following line executes Optimus with the above specified inputs:

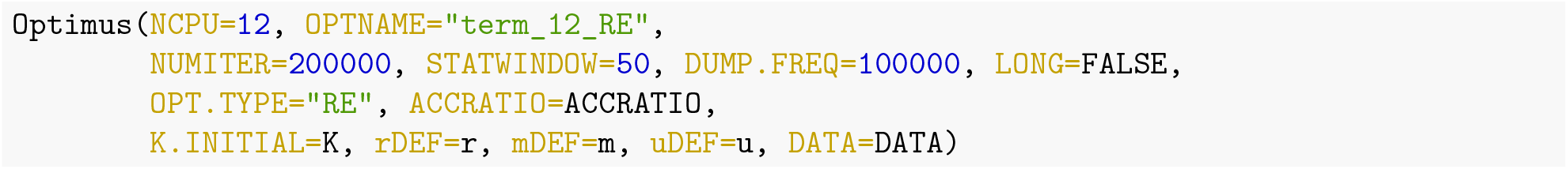

Nine of the optimisation replicas (CPUs 1, 2, 3, 5, 6, 8, 9, 10 and 12) recovered the same solution that was found by the Acceptance Ratio Simulated Annealing Optimus run. Moreover, this solution is better (lower *AIC*) than those recovered by CPUs 4, 7 and 11. Thus, in this example, the Acceptance Ratio Simulated Annealing and Replica Exchange versions produce the same solution.

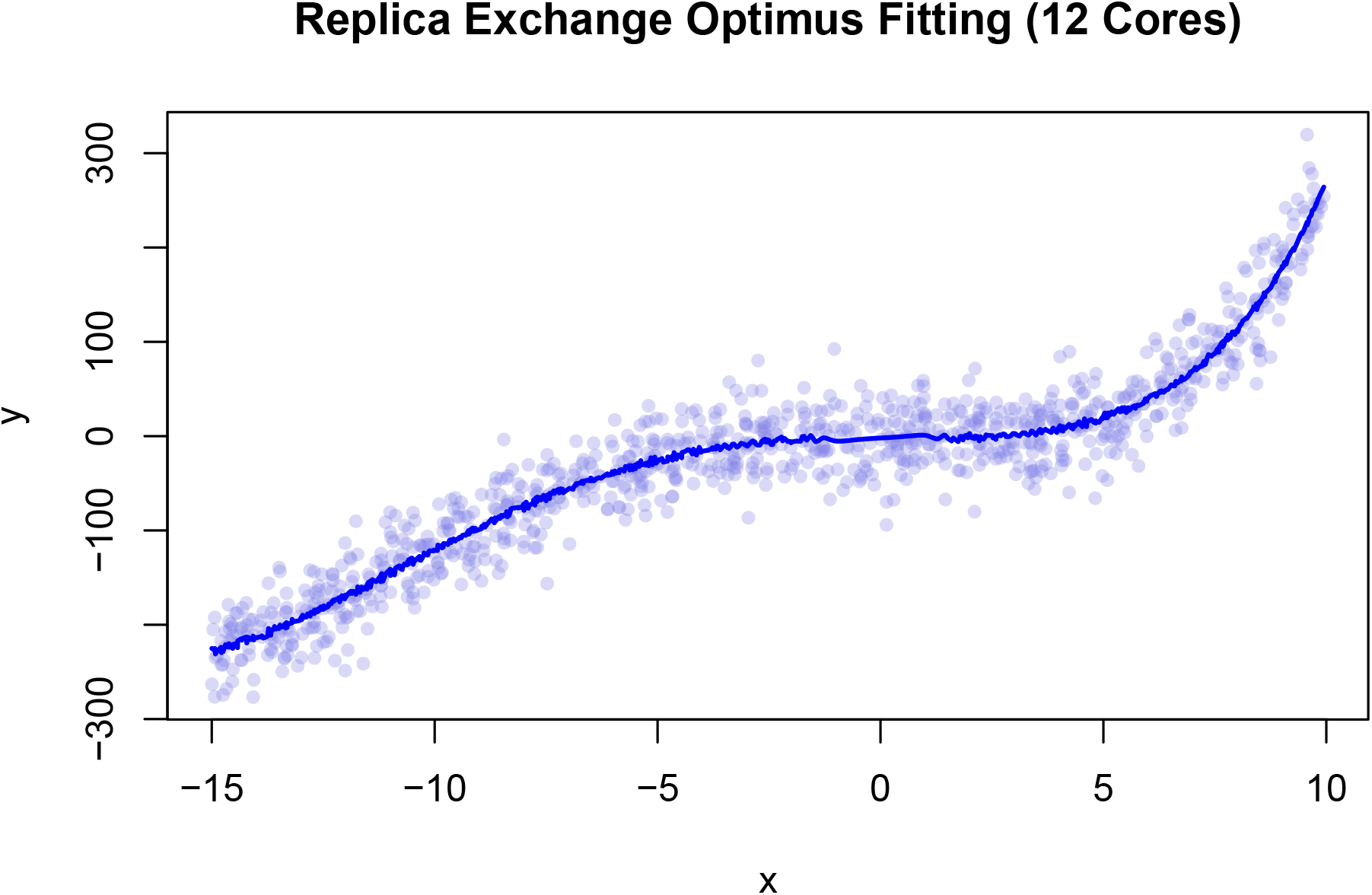

Please note that for convenience, only those replicas that produced a unique solution are listed in the table below (replicas 1, 2, 3, 6, 8, 9, 10 and 12 produced the same solution as replica 5; replica 11 produced the same solution as replica 7).

**Table 5:** 12-core Acceptance Ratio Replica Exchange results from Optimus run.

|  | CPU 4 | CPU 5 | CPU 7 |
| --- | --- | --- | --- |
| Replica Acceptance Ratio | 66.0000 | 58.0000 | 42.0000 |
| E (AIC) | 9.5673 | 9.5672 | 9.5673 |
| Q (RMSD) | 28.6664 | 28.6932 | 28.6951 |
| Term 1 | 0.0000 | 0.0000 | 0.0000 |
| Term 2 | 1.0000 | 1.0000 | 1.0000 |
| Term 3 | 1.0000 | 1.0000 | 1.0000 |
| Term 4 | 1.0000 | 1.0000 | 1.0000 |
| Term 5 | 0.0000 | 0.0000 | 0.0000 |
| Term 6 | 0.0000 | 0.0000 | 0.0000 |
| Term 7 | 0.0000 | 0.0000 | 0.0000 |
| Term 8 | 0.0000 | 0.0000 | 0.0000 |
| Term 9 | 0.0000 | 0.0000 | 0.0000 |
| Term 10 | 0.0000 | 0.0000 | 0.0000 |
| Term 11 | 1.0000 | 1.0000 | 0.0000 |
| Term 12 | 0.0000 | 0.0000 | 0.0000 |
| Term 13 | 1.0000 | 0.0000 | 1.0000 |
| Term 14 | 0.0000 | 0.0000 | 0.0000 |
| Term 15 | 0.0000 | 0.0000 | 0.0000 |
| Term 16 | 0.0000 | 0.0000 | 0.0000 |
| Term 17 | 0.0000 | 0.0000 | 0.0000 |
| Term 18 | 0.0000 | 0.0000 | 0.0000 |
| Term 19 | 0.0000 | 0.0000 | 0.0000 |
| Term 20 | 1.0000 | 1.0000 | 0.0000 |
| Term 21 | 0.0000 | 0.0000 | 0.0000 |
| Term 22 | 0.0000 | 0.0000 | 0.0000 |
| Term 23 | 0.0000 | 0.0000 | 0.0000 |
| Term 24 | 0.0000 | 0.0000 | 0.0000 |
| Term 25 | 0.0000 | 0.0000 | 0.0000 |
| Term 26 | 1.0000 | 1.0000 | 1.0000 |
| Term 27 | 0.0000 | 0.0000 | 0.0000 |
| Term 28 | 0.0000 | 0.0000 | 1.0000 |
| Term 29 | 0.0000 | 0.0000 | 0.0000 |
| Term 30 | 0.0000 | 0.0000 | 0.0000 |

Note that the various replica outcomes illustrate the penalising effects of the *AIC* on models using a greater number of parameters. Consider the solutions found by the 66% acceptance ratio replica and the 58% acceptance ratio replica (CPUs 4 and 5 respectively). Let *y_i_* denote the solution found by CPU *i*. Then, we have:

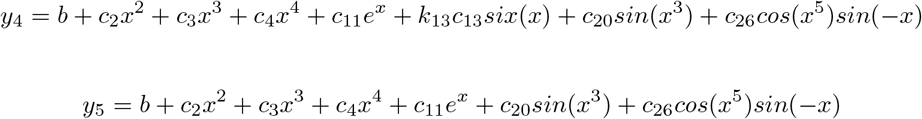

Although the RMSD of *y*_4_, 28.6664, is lower than the RMSD of *y*_5_, 28.6932, *y*_5_ has a lower value for *AIC* because it contains one less term than *y*_4_. Since *AIC* was specified as the objective metric, Optimus (perhaps counterintuitively) selected *y*_5_ as the more optimal solution to reduce overfitting.

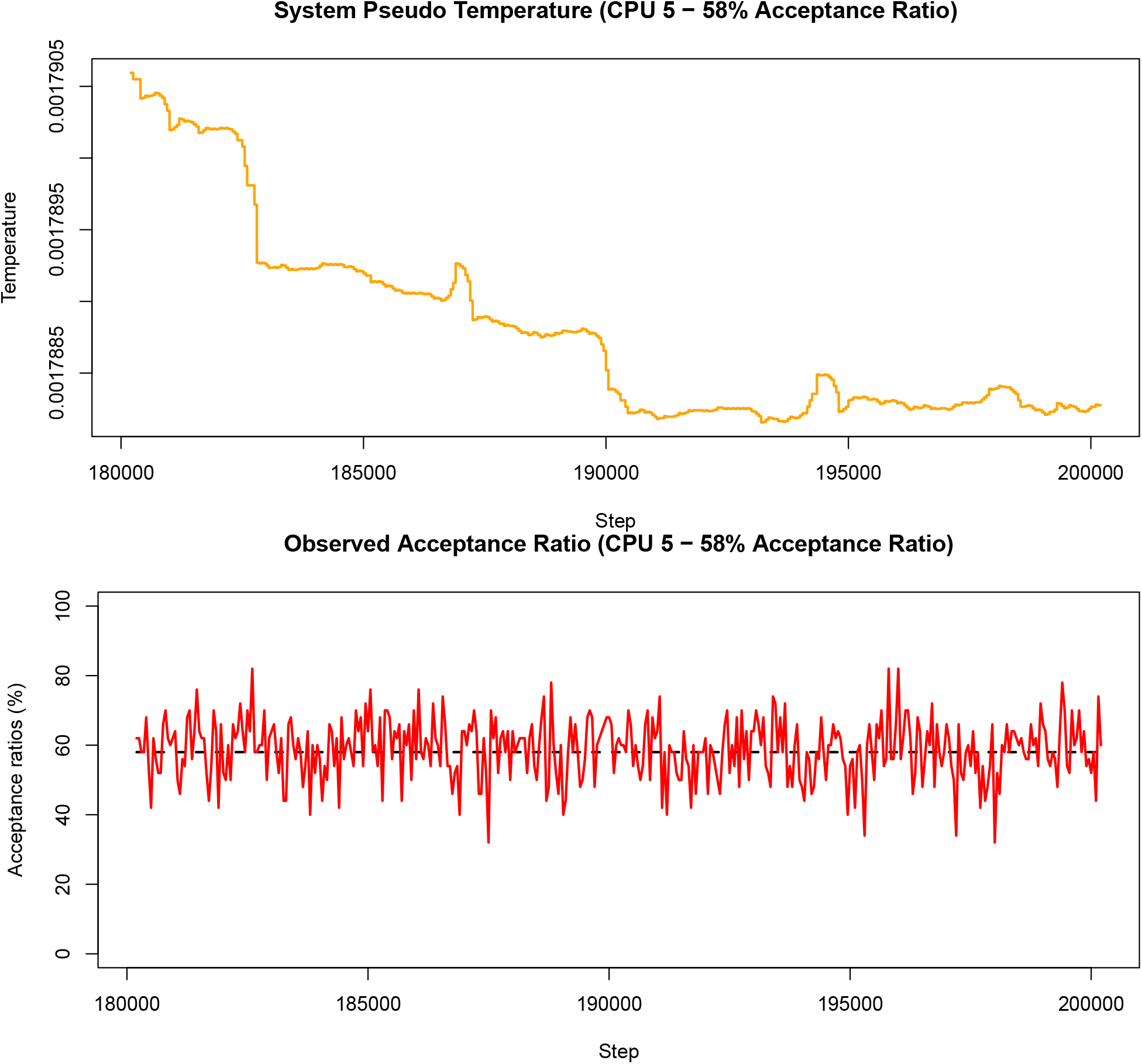

The above graphs are produced using data from the last 20 000 iterations of the 58% acceptance ratio replica (CPU 5). It is clear that the observed acceptance ratio more strongly oscillated around the target acceptance ratio than was the case in the Acceptance Ratio Simulated Annealing run from the previous part of this tutorial. More generally, it should be expected that the observed acceptance ratio fluctuates more significantly around the target acceptance ratio in Replica Exchange than in Simulated Annealing, especially when the objective function is non-smooth as is the case in this example. This is because an exchange between two replicas has the same effect as restarting a Monte Carlo optimisation from a random initial configuration with a temperature that very likely is not conducive to the target acceptance ratio for the given configuration. As such, each time an exchange occurs, significant deviations from the target acceptance ratio may occur and may require several `STATWINDOW`s for the TCU to correct. Despite this challenge, the TCU performs satisfactorily.

#### Summary

We now understand how to employ Optimus to solve a more general problem than was addressed in **Tutorial 1** and one with a non-smooth objective function. Additionally, we have a better understanding of the adaptive thermoregulation. Using the Akaike Information Criterion (*AIC*) as a metric with which to evaluate the performance of a candidate model, taking into account the desire to represent the data while avoiding to overfit the data, both the Acceptance Ratio Simulated Annealing and Replica Exchange versions of Optimus recovered a better functional form to describe the data than the form which was assumed in **Tutorial 1** (based on how the data had been generated), obviously with some overfitting to adapt to the noise while with the stringency used in this example.

**Table 6:** Summary of solutions.

|  | E (AIC) | Q (RMSD) |
| --- | --- | --- |
| Least Squares (Tutorial 1) | 9.5705 | 28.82655 |
| Optimus (AR Simulated Annealing) | 9.5672 | 28.69324 |
| Optimus (AR Replica Exchange) | 9.5672 | 28.69324 |

Least Squares (**Tutorial 1**):

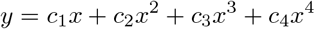

Optimus (Acceptance Ratio Simulated Annealing):

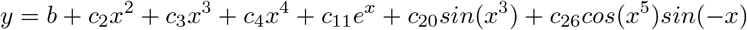

Optimus (Acceptance Ratio Replica Exchange):

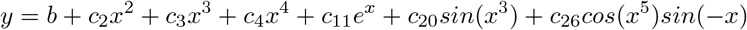

### Tutorial 3: Geometry Optimisation of Vitamin C Molecule

#### Problem Statement

The focus of this tutorial is to depart from problem classes involving the search for functions to represent data, and demonstrate how Optimus can be flexibly applied to arbitrary problem classes provided that they are formulated in accordance with Optimus specifications. Additionally, this tutorial will illustrate that Optimus can act as an optimisation kernel while calling external programs to execute a significant amount of the necessary computation for the optimisation process.

In this tutorial, Optimus will be used, as an illustrative example, to 3D geometry optimise a molecular structure. Specifically, Optimus will be used to determine the optimal values of two dihedral angles in the L-ascorbic acid (Vitamin C) molecule such that the molecule is in its ground state energy conformation. Vitamin C was selected to be the studied molecule because it has more than one freely rotating carbon-carbon bond and the potential for intramolecular hydrogen bonding due to the presence of multiple hydroxyl groups. Moreover, Vitamin C is not a particularly large molecule. Due to these circumstances, Vitamin C can serve as a non-trivial case (as opposed to simpler molecules like ethane for instance), but one that does not require several days or weeks of calculations to arrive to optimal solutions (the optimisation procedures below took roughly 14-18 hours to terminate).

This is the molecular structure of Vitamin C, with the numbering of non-hydrogen atoms provided from the scheme used in geometry specification:

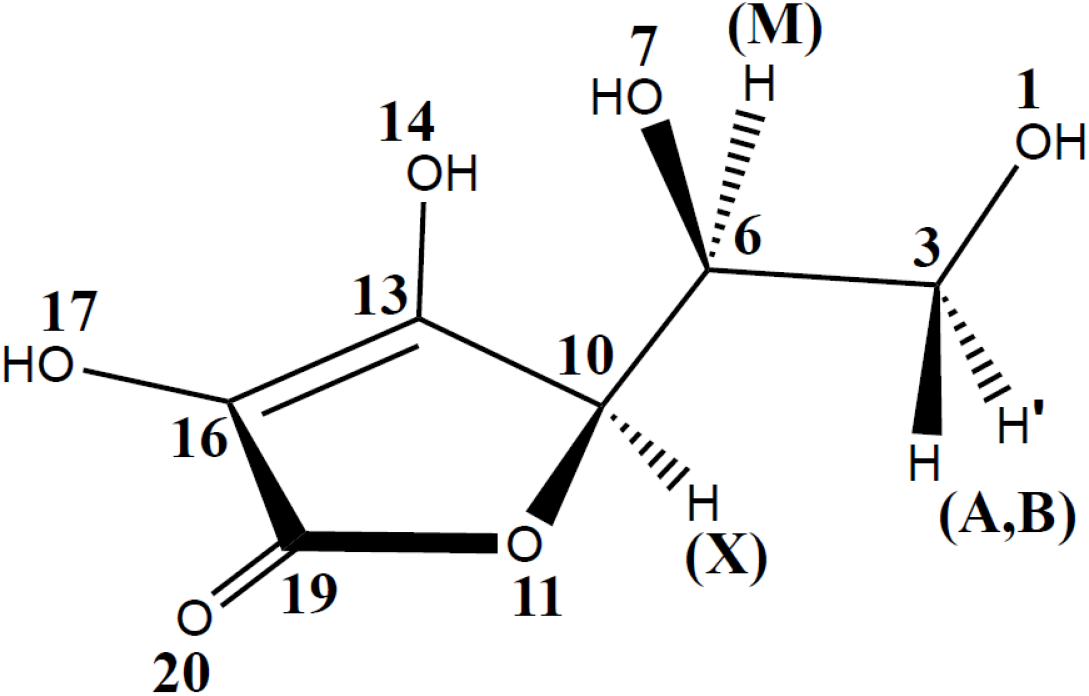

The major geometric features that drive the overall state of the molecule are the two C-C bonds in this structure: the bond joining carbon 3 and 6, and the bond joining carbon 6 and 10. The ground state conformation of Vitamin C will likely be a conformation such that steric clashes are minimised while also allowing for close proximity and right orientation between hydrogen bond donor and acceptor atoms. In the following sections, we formalise this optimisation problem and use Optimus to arrive at the solution.

#### Defining Optimus Inputs

As in the previous tutorials, we must first rigorously define the parameters that we are optimising. Let us begin by defining a dihedral angle as it will be used to our molecular geometry: a dihedral angle is the angle between two intersecting planes, where each plane is specified by 3 atoms of which 2 are common between both planes. Thus, a total of 4 atoms are needed to specify a dihedral angle. The conformation of Vitamin C with respect to its two freely rotating C-C bonds can be specified *via* two dihedral angles. Let *ψ* be the dihedral angle defined by the atoms numbered 1, 3, 6 and 7 and let *ϕ* be the dihedral angle defined by the atoms numbered 7, 6, 10 and 11. Having defined these two angles, we can now define the parameter set `K` as a numeric vector of length 2 whose entries are *ψ* and *ϕ*. We will arbitrarily initialise *ψ* and *ϕ* to have value 180. The corresponding Vitamin C conformation is illustrated below using a 3D structure and Newman projections along the two rotatable carbon-carbon bonds:

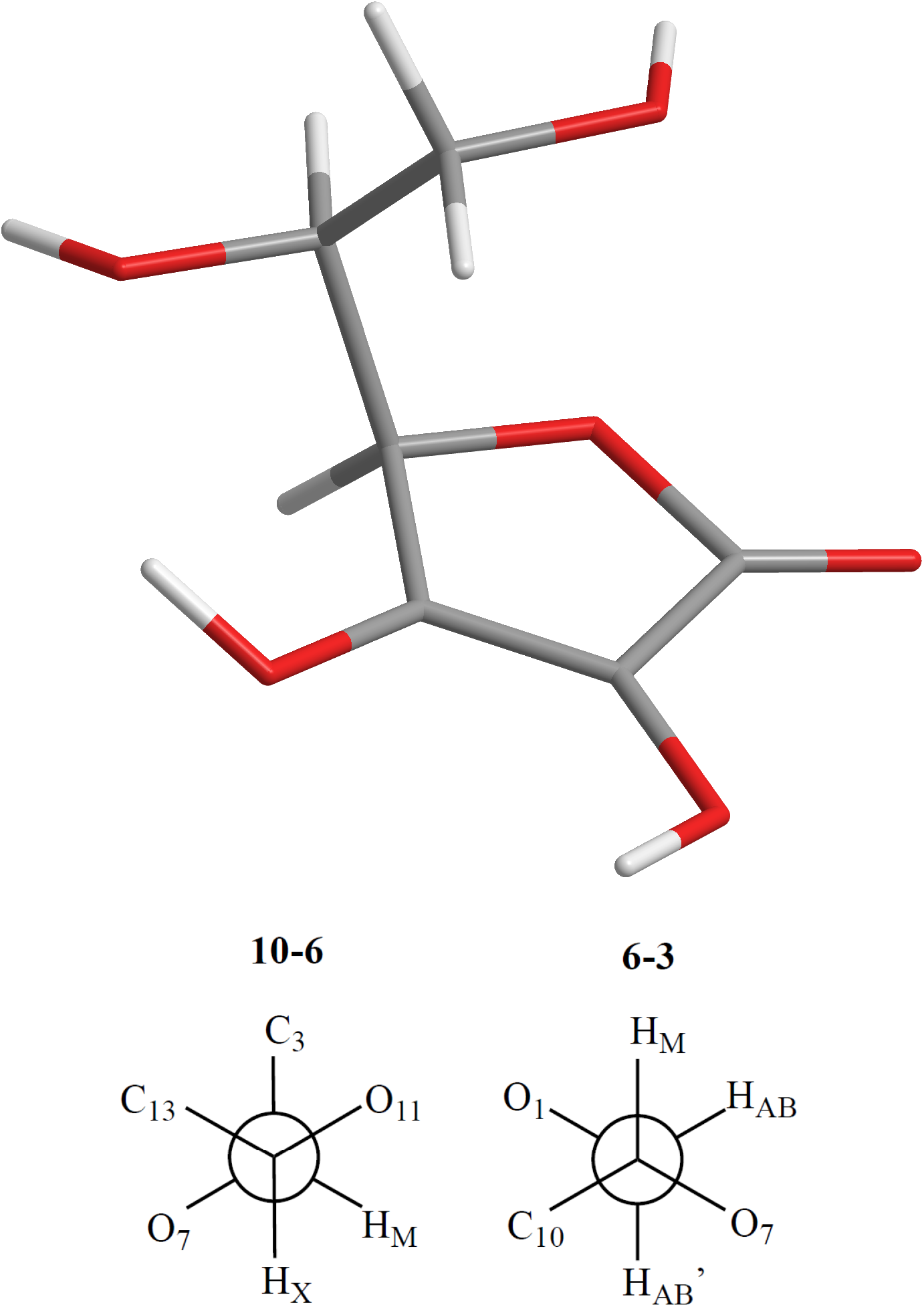

In the 3D structure, grey denotes carbon, red denotes oxygen and white denotes hydrogen atoms.

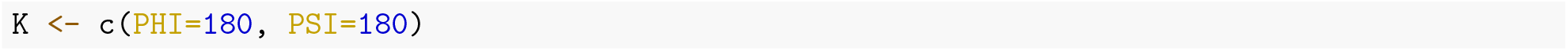

Now we will specify a model function `m()` that will operate on `K`. Starting from an arbitrary molecular conformation, altering the value of `K` will likely cause certain clashes or non-optimal interactions between atoms in the molecule that are not used in the definition of the angles *ψ* and *ϕ*. As such, after receiving an input set of parameters `K`, `m()` will have to alter the 3D location of constituents atoms while holding `K` fixed to arrive at the most stable geometry for the input `K`. Here, unlike in previous tutorials, to accomplish this task `m()` will call an external program MOPAC. MOPAC is a program for semiempirical quantum mechanics (QM) calculations, and can perform constrained and unconstrained geometry optimisations to arrive at a stationary state (note that calling MOPAC for a single initial geometry instance does not guarantee a global minimum will be found). MOPAC takes as input the specification of an initial molecular geometry in addition to an indication of which molecules the program is able to displace (or angles it can alter) and outputs a nearby local minimum molecular conformation with its corresponding energy in kcal/mol. For this optimisation problem, the input to MOPAC will be structured as a Z matrix, a common form for describing a molecular conformation which consists of using lengths, angles and dihedral angles with respect to previously defined atoms to define new atoms in the conformation.

The function `m()` will construct a Z matrix for Vitamin C using the input dihedral angles `K` and default values for the remaining geometric relationships needed to define the molecule. `m()` will then call MOPAC with the newly constructed Z matrix, specifying that all relationships may be altered by QM optimisation, except the input dihedral angles `K`. Finally, `m()` will return the energy calculated by MOPAC *via* PM6 Hamiltonian.

Note that to avoid non-convergence issues when calling MOPAC, `m()` returns a default energy value of −100 kcal/mol if a call to MOPAC does not terminate within 10 seconds (over-simplifications just for the sake of this illustrative example). Also, note that although `m()` requires no additional data on top of `K` to operate, `m()` must still be defined to take an input `DATA` in accordance with Optimus specifications. Lastly, note that a local installation of MOPAC (2016) is required to execute this optimisation procedure. Below is the definition of `m()` :

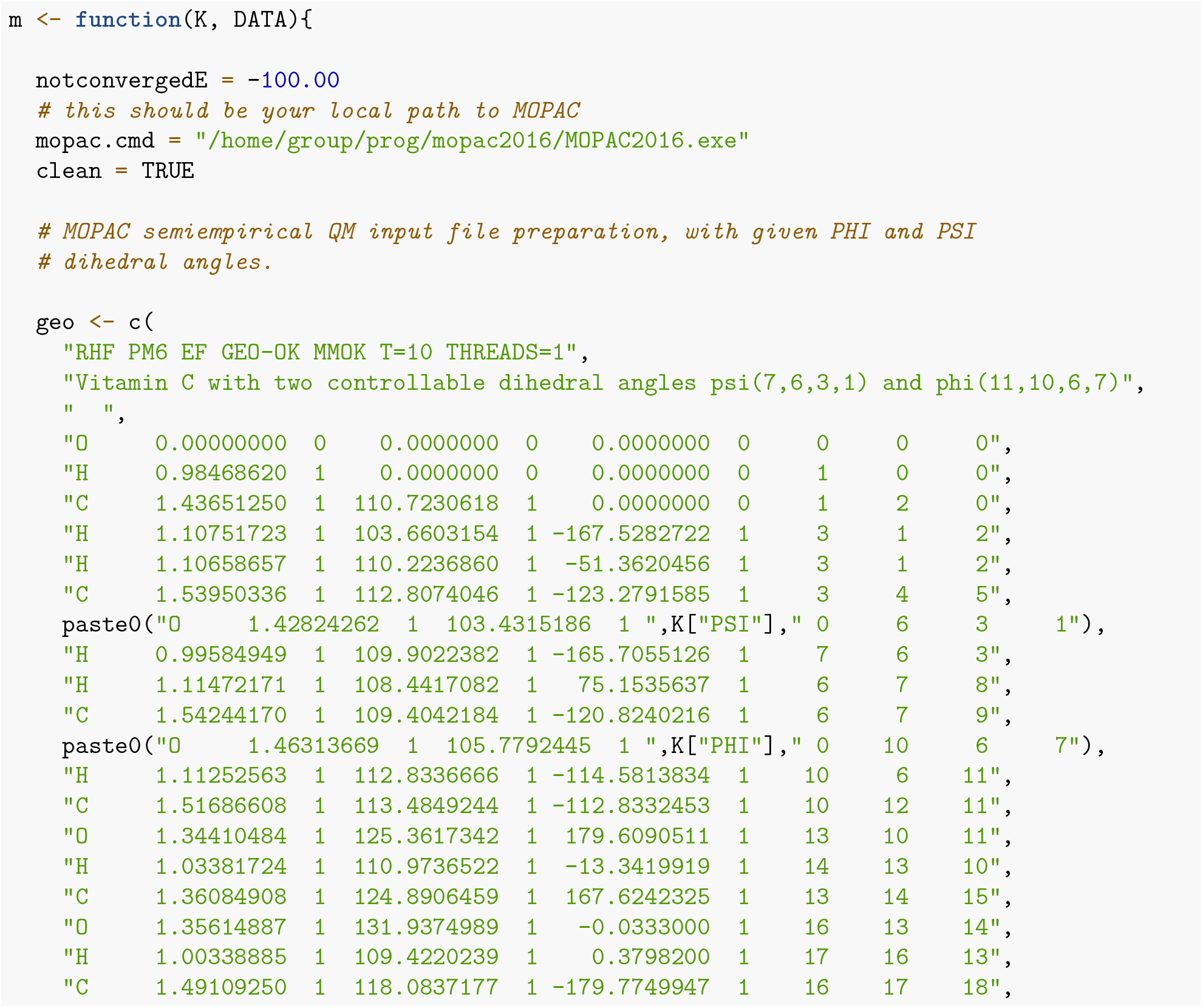
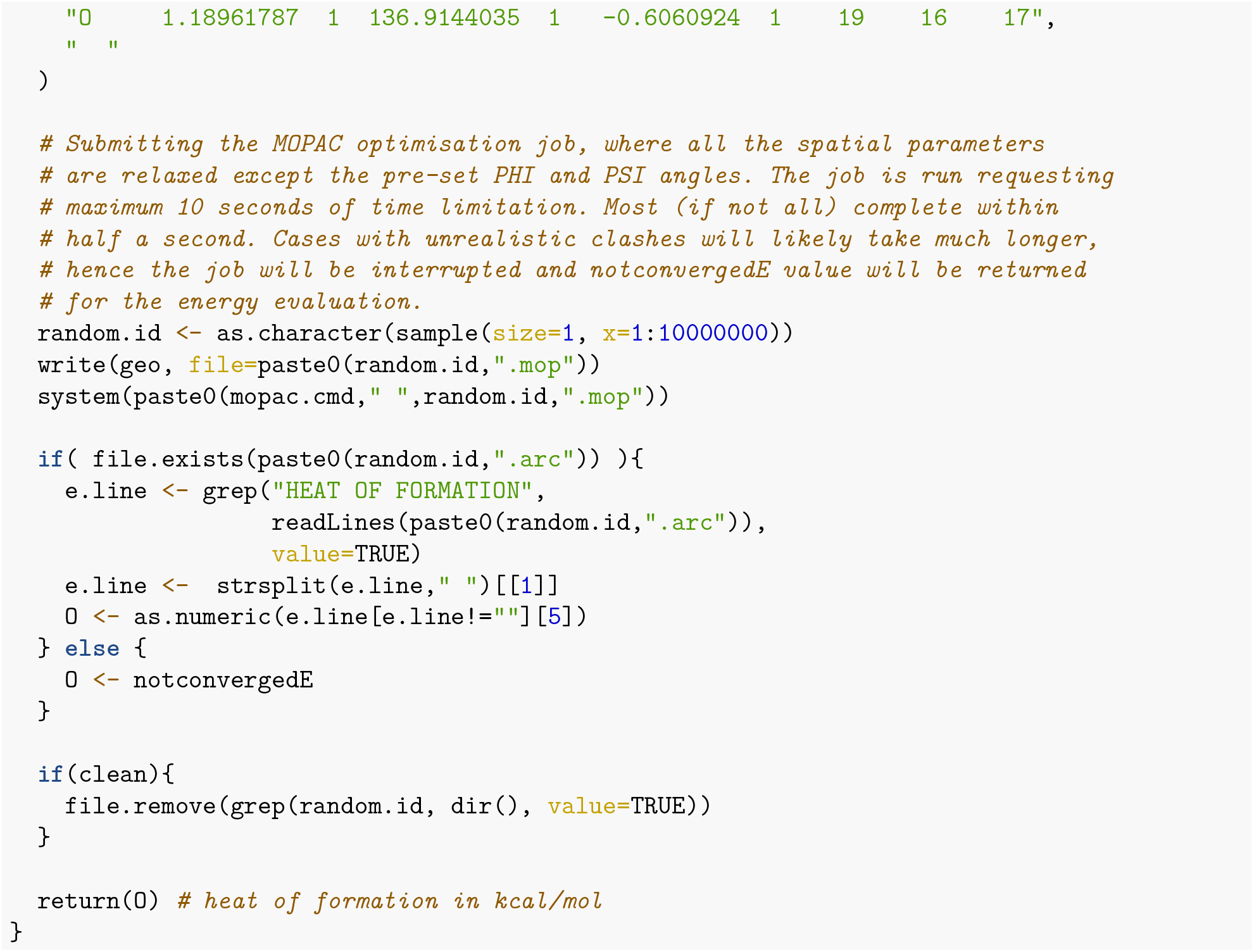

Next, we define the function `u()` which returns an energy `E` and a quality `Q` of the candidate solution. Since the `m()` will already output a value for the physical energy of the candidate Vitamin C conformation, `u()` can simply set `E` to be the same return value of `m()`. We will make `u()` set `Q` to be the negative of the return value of `m()` such that candidate conformations with lower energies produce higher values of quality `Q`. Again, although `u()` does not require any additional data to accomplish this functionality, it must nevertheless be written to optionally accept an input parameter `DATA`.

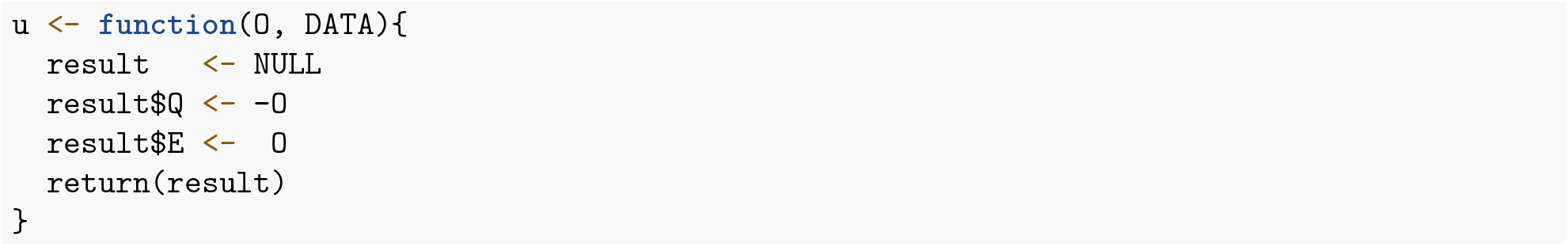

Finally, we define the alteration function `r()`. `r()` will randomly select either *ψ* or *ϕ* to alter. Thereafter, `r()` randomly increases or decreases the selected angle by 2 degrees. `r()` will also ensure that *ψ, ϕ* ∈ [-180.0, 180.0] throughout the optimisation process.

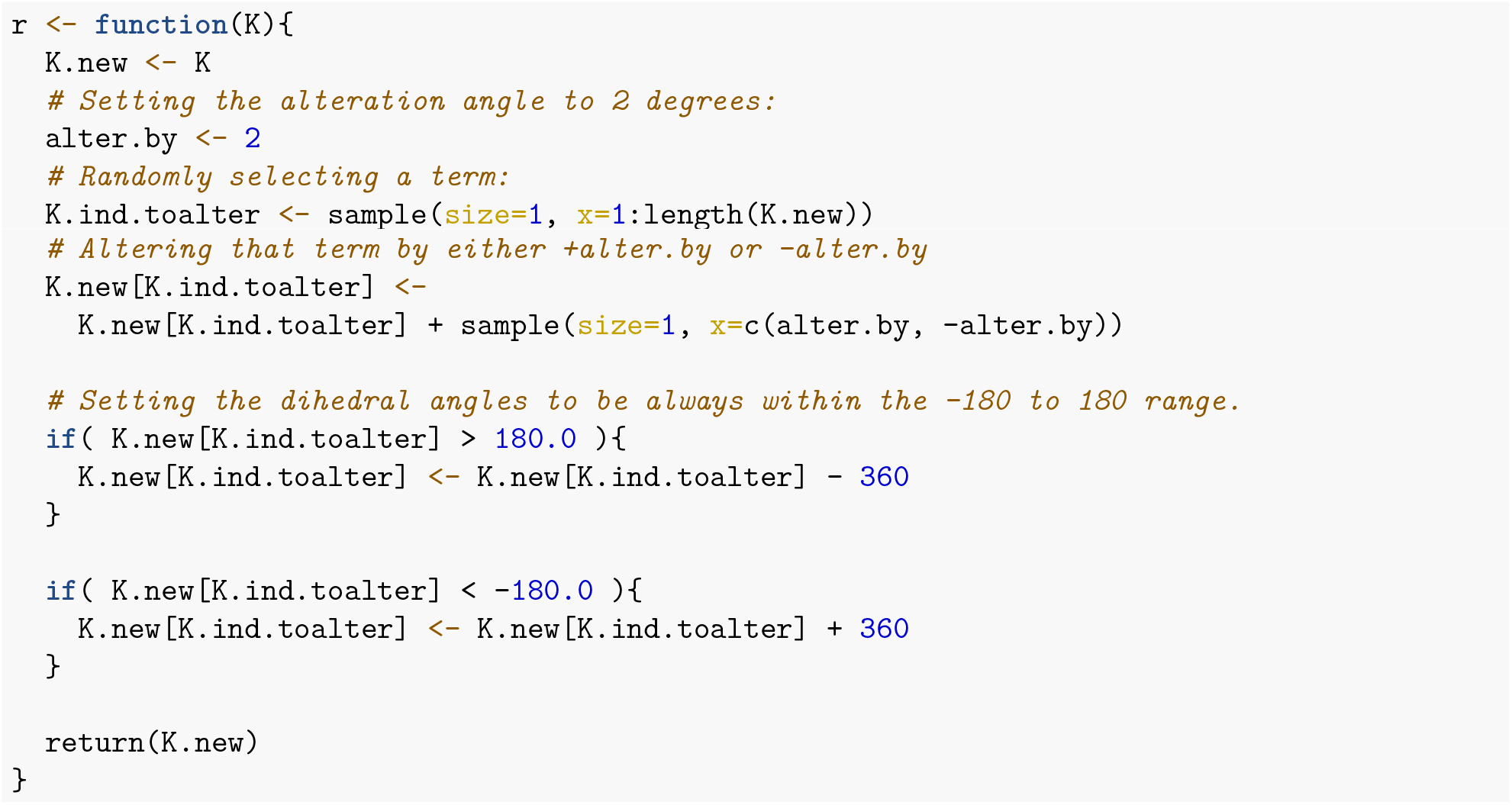

The process of determining the energy of a conformation corresponding to a given set of angles *ψ, ϕ* is the most computationally intensive part of this optimisation formulation. Having defined the necessary inputs for Optimus, it should be apparent that this calculation will entirely be handled by MOPAC. This ability to serve as an optimisation kernel and flexibly be knitted to an external program is one of the many strengths of Optimus.

#### Defining a Benchmark Solution

Before calling Optimus, we have established a benchmark solution to be used to independently evaluate the ability Optimus to arrive to correct *ψ* and *ϕ* combination. In order to explore the energy landscape associated with the parameter space of *ψ* and *ϕ*, a PM6 optimization was performed on 10 conformers picked from the wells pf a more comprehensive potential energy surface scanning through MM2 molecular mechanics force field (the details of PM6 and MM2 are not important for the purposes of this tutorial). This resulted in the identification of 7 energy minima, shown in the table below (listed in increasing order by energy):

**Table 7:** Seven conformational minima calculated for Vitamin C with PM6.

|  | E (kcal/mol) | PHI | PSI |
| --- | --- | --- | --- |
| Conformation 1 | -233.206 | 92.58 | 74.19 |
| Conformation 2 | -232.877 | 50.60 | -172.79 |
| Conformation 3 | -231.800 | -169.67 | -41.23 |
| Conformation 4 | -230.822 | 47.43 | -166.61 |
| Conformation 5 | -230.274 | -172.20 | -54.45 |
| Conformation 6 | -225.214 | -75.69 | -104.44 |
| Conformation 7 | -224.875 | -73.31 | 155.02 |

We will assume that the above listed conformations comprehensively represent most, if not all, of the possible minima in the parameter space of *ψ* and *ϕ*. Under this assumption, Conformation 1 should be considered as the ground state conformation of Vitamin C. The accuracy of the results produced by Optimus can thus be judged by comparing them to the data listed in the above table. It is important to recognize that the “resolution” of *ψ* and *ϕ* when being optimised by Optimus is set to 2 degrees due to the manner in which `r()` was defined. As such, results produced by Optimus that are within plus or minus 2 degrees of a reference conformation should be tolerated.

#### Acceptance Ratio Simulated Annealing Optimus Run

For the Acceptance Ratio Annealing run, we will set `NUMITER` to 100 000 because each optimsation step is more costly due to the relatively computationally expensive calls to MOPAC. Moreover, we will set `CYCLES` to 2. Although this shortens the length of an annealing cycle to 50 000 steps (whereas 100 000 steps per cycle has been kept constant over the previous tutorials), having more than 1 annealing cycle is likely more beneficial than insisting on a cycle lasting 100 000 steps as opposed to 50 000.

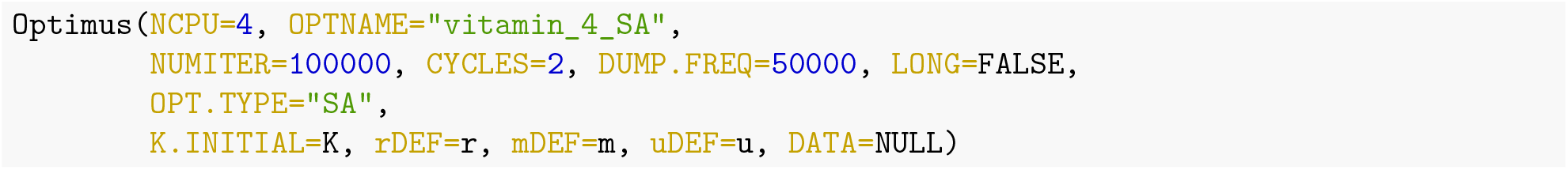

**Table 8:**
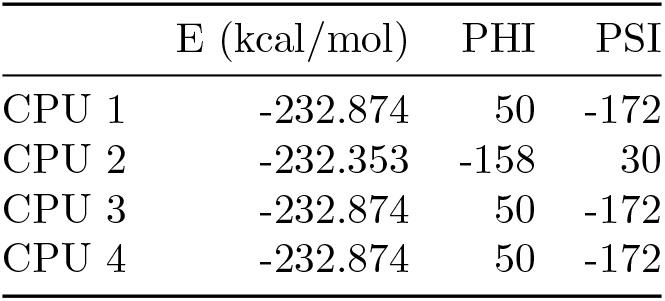
4-core Acceptance Ratio Simulated Annealing results from Optimus runs.

|  | E (kcal/mol) | PHI | PSI |
| --- | --- | --- | --- |
| CPU 1 | -232.874 | 50 | -172 |
| CPU 2 | -232.353 | -158 | 30 |
| CPU 3 | -232.874 | 50 | -172 |
| CPU 4 | -232.874 | 50 | -172 |

CPUs 1, 3 and 4 all arrived at a conformation defined by {*ϕ* = 50, *ψ* = −172}, with an energy of −232.874 kcal/mol. The below 3D structure and Newman projections depict this solution:

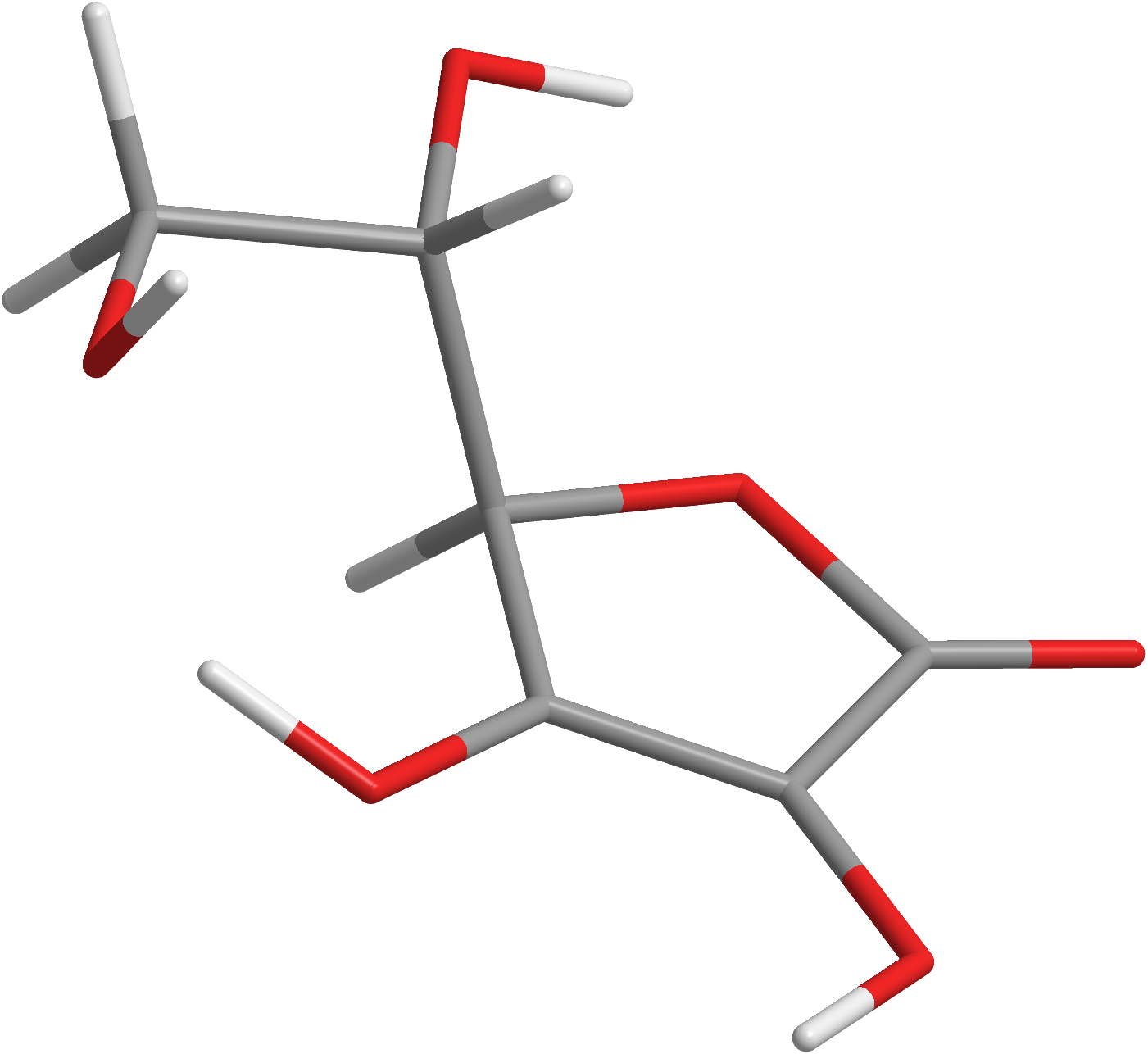
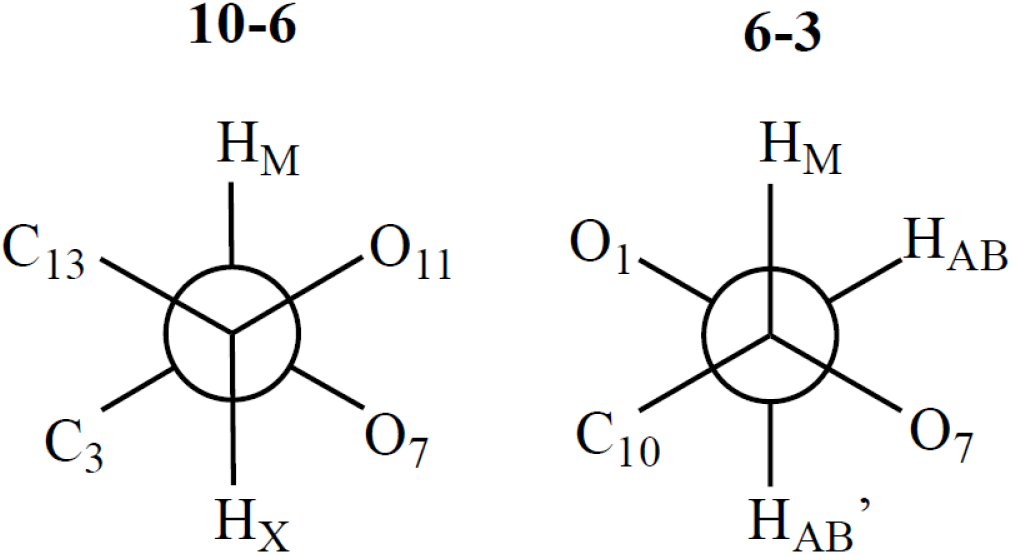

This conformation is equivalent to benchmark Conformation 2. Thus, in this example, Acceptance Ratio Simulated Annealing was able to find the Vitamin C conformation with the second lowest energy in the parameter space. This performance is strong, especially given that the limited steps and cycles executed, and that the energy difference between Conformation 1 and Conformation 2 is only −0.329 kcal/mol.

The graphs below illustrate the system psuedo temperature and observed acceptance ratio for the last 20 000 optimisation iterations executed by CPU 3.

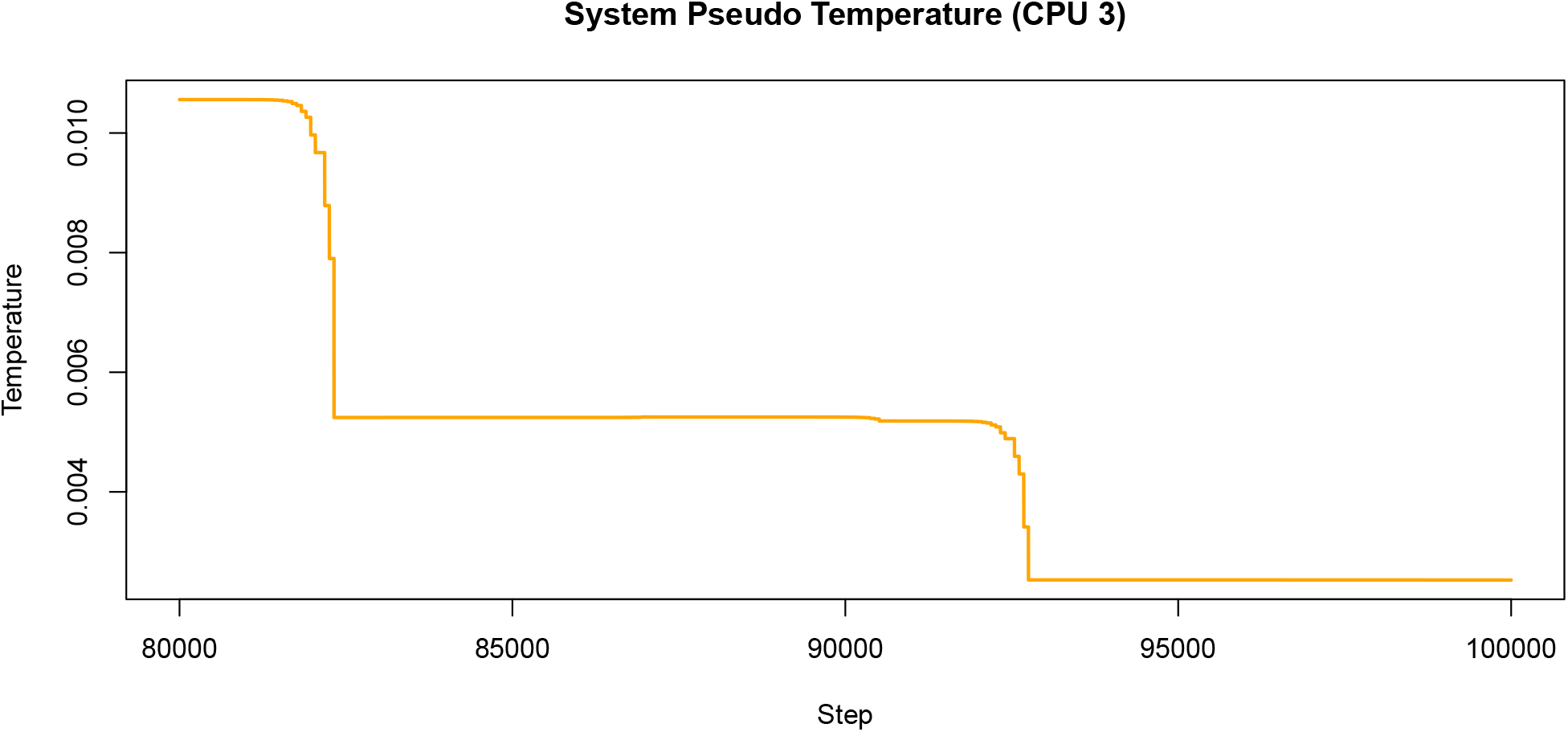
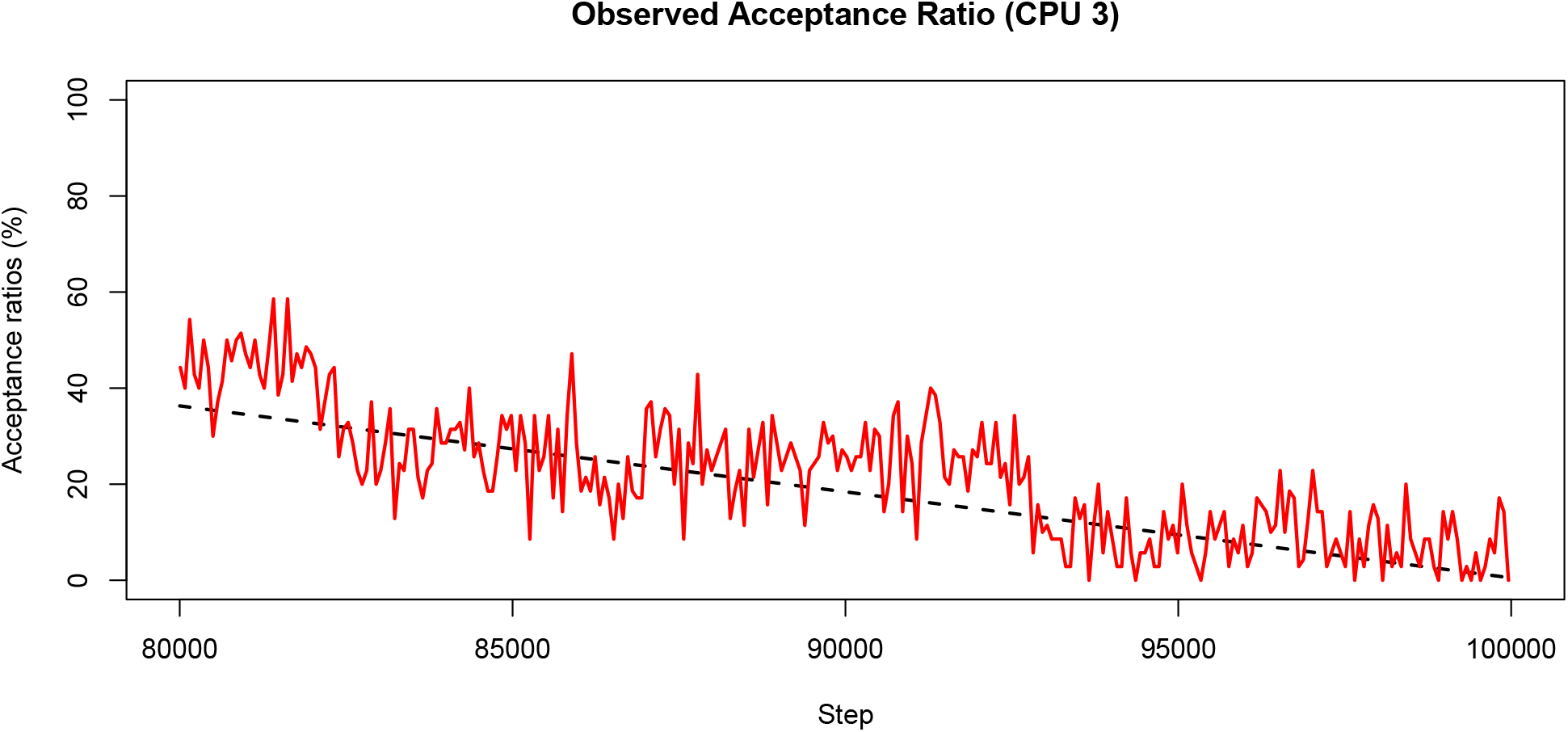

#### Acceptance Ratio Replica Exchange Optimus Run

Let us now consider the Replica Exchange version of Optimus on 12 processors with the variable `ACCRATIO` defined as in the previous tutorials.

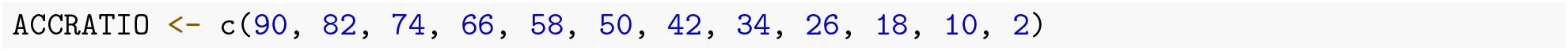

Just as in the Acceptance Ratio Simulated Annealing run, we will set `NUMITER` to 100 000. Furthermore, we will set `EXCHANGE.FREQ` to 500 such that the number of iterations between subsequent exchanges between replicas is 200 as it was in **Tutorial 2**. For the same reasons as in **Tutorial 2**, we will set `STATWINDOW` to 50 for the Replica Exchange run.

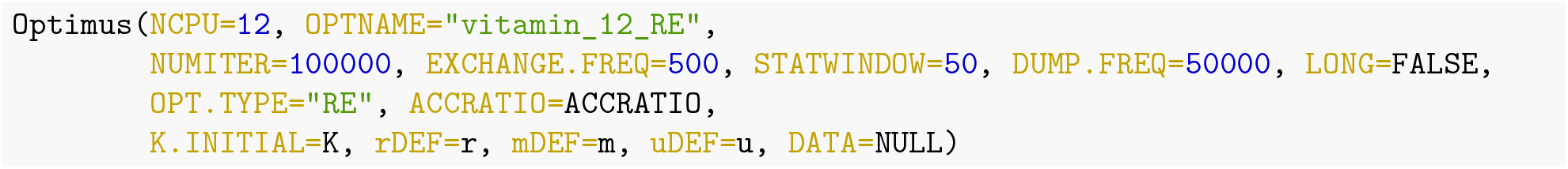

**Table 9:** 12-core Acceptance Ratio Replica Exchange results from Optimus run.

|  | Replica Acceptance Ratio | E (kcal/mol) | PHI | PSI |
| --- | --- | --- | --- | --- |
| CPU 1 | 90 | -229.2359 | -164 | -178 |
| CPU 2 | 82 | -229.2359 | -164 | -178 |
| CPU 3 | 74 | -233.1453 | 82 | 84 |
| CPU 4 | 66 | -233.1979 | 90 | 76 |
| CPU 5 | 58 | -229.2359 | -164 | -178 |
| CPU 6 | 50 | -232.8742 | 50 | -172 |
| CPU 7 | 42 | -233.1947 | 94 | 74 |
| CPU 8 | 34 | -229.2359 | -164 | -178 |
| CPU 9 | 26 | -229.2359 | -164 | -178 |
| CPU 10 | 18 | -227.6394 | 180 | 158 |
| CPU 11 | 10 | -229.2359 | -164 | -178 |
| CPU 12 | 2 | -229.2359 | -164 | -178 |

Of the 12 replicas, CPU 4 recovered the conformation with the lowest energy (−233.1979), defined by {*ϕ* = 90, *ψ* = 76}. The below 3D structure and Newman projections depict this solution:

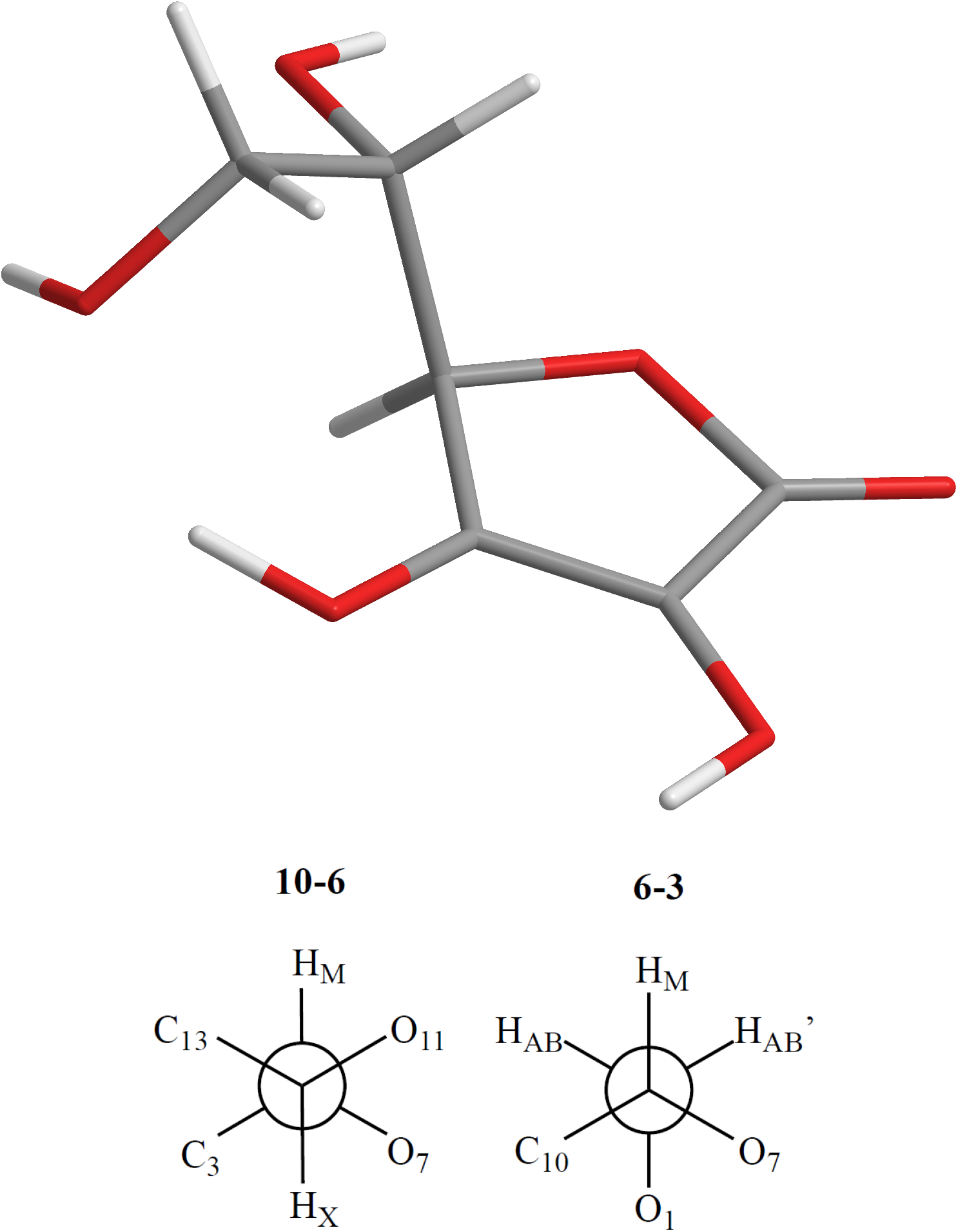

This solution corresponds to reference Conformation 1, the global minimum energy state for Vitamin C. Thus, for this optimisation problem, under the limits of the set number of steps and cycles, the Replica Exchange version of Optimus outperformed Acceptance Ratio Annealing by succeeding in finding the global minimum of the energy landscape.

If we compare the solution found by CPU 4 to benchmark Conformation 1, it is evident that the value for *ϕ* found by Optimus lies slightly outside of the plus or minus 2 degree window that was discussed earlier. Contrarily, CPU 7 finds a solution {*ϕ* = 94, *ψ* = 74} which does lie strictly within the resolution window. Despite this, the solution of CPU 4 has a slightly lower energy (−233.1979) than the solution of CPU 7 (−233.1947) and so represents a better solution. Finally, notice that Replica 6 recovered the same conformation that was identified by Acceptance Ratio Simulated Annealing Optimus.

The below graphs illustrate the system pseudo temperature and observed acceptance ratio for the first 20 000 optimisation iterations executed by CPU 4 (66% acceptance ratio replica).

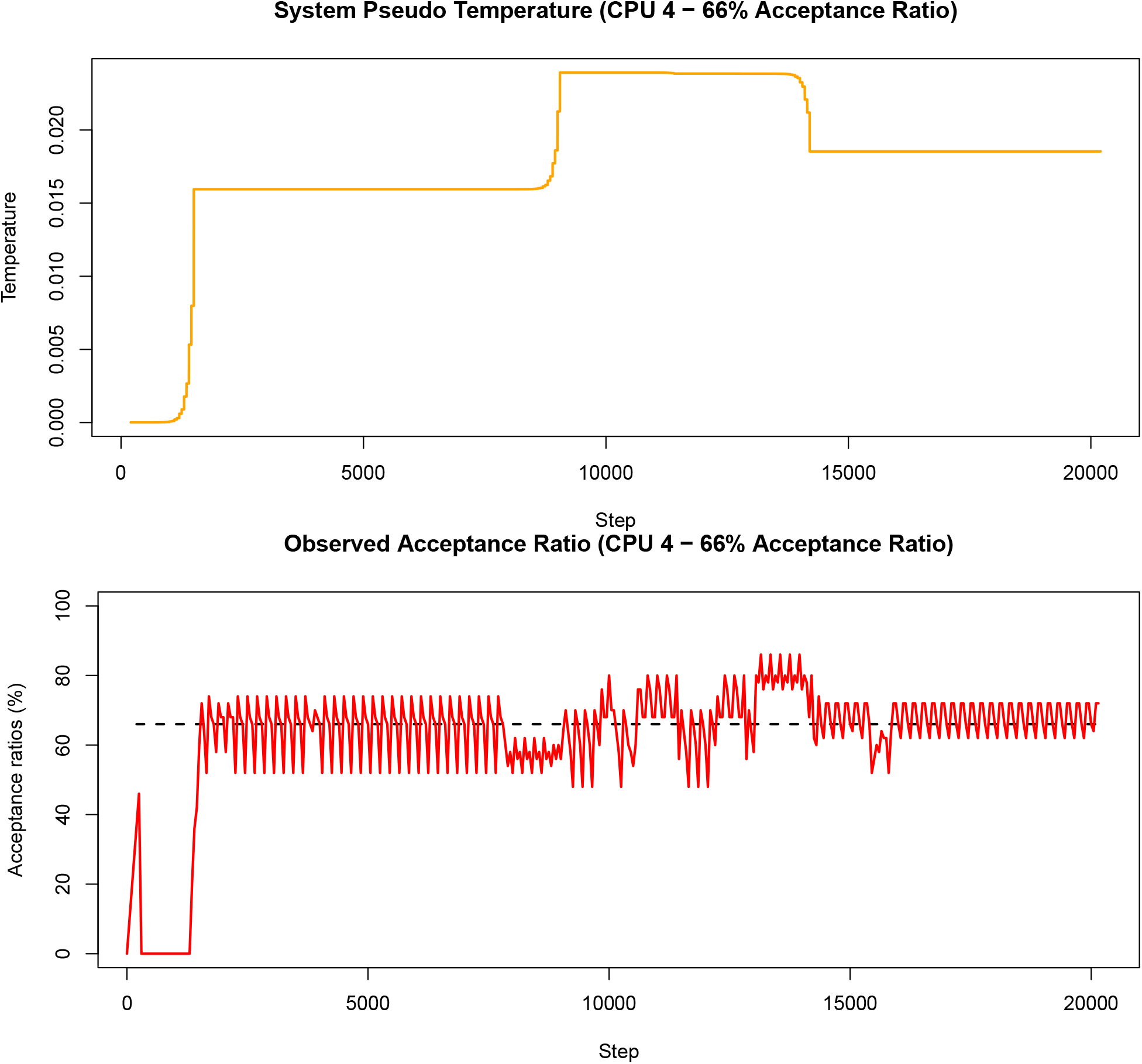

When the optimisation process is first initialised, it is very unlikely that the input initial temperature is conducive to the target acceptance ratio. As such, the adaptive thermoregulation alters the system pseudo-temperature considerably and rapidly to align the observed acceptance ratio with the target acceptance ratio, as can be seen in the above two graphs. Moreover, as stated in the previous tutorial, an exchange between two replicas often has a similar effect of introducing a parameter configuration that is not conducive to the current system pseudo temperature, which necessitates significant temperature adjustments. Accordingly, sharp increases or decreases in the system pseudo temperature and significant changes in the value around which the observed acceptance ratio oscillates in the graph above likely indicate steps at which an exchange involving replica 4 occurred.

#### Summary

We are now familiar with how to structure a more complex optimisation problem, involving an external program, to be solved with Optimus as a kernel. On the particular example of geometry optimisation here, we saw that the Simulated Annealing mode of Optimus was able to find the second lowest local minimum (under restricted number of annealing cycles), while the Replica Exchange mode recovered the global energy minimum.

**Table 10:** Summary of solutions.

|  | Energy (kcal/mol) | PHI | PSI |
| --- | --- | --- | --- |
| Ground State Reference | -233.2060 | 92.58 | 74.19 |
| Optimus (AR Simulated Annealing) | -232.8740 | 50.00 | -172.00 |
| Optimus (AR Replica Exchange) | -233.1979 | 90.00 | 76.00 |

### Tutorial 4: Exploring Coupled ODEs Modelling a Biological System

#### Problem Statement

This Tutorial will demonstrate the use of Optimus to address a problem from yet another problem class. We will employ Optimus to recover the rate constants for a system of coupled ordinary differential equations (ODEs) modelling a biological pathway. Specifically, we will study a phosphorelay system from the high osmolarity glycerol (HOG) pathway in yeast. A phosphorelay system is a network involving multiple proteins in which, after an initial phosphorylation event using ATP (or an alternate phosphate donor), the phosphorylation and dephosphorylation events of proteins in the network proceed without further consumption of ATP (Klipp et al. 2009). The below diagram illustrates the phosphorelay system that will be studied in detail (Klipp et al. 2009):

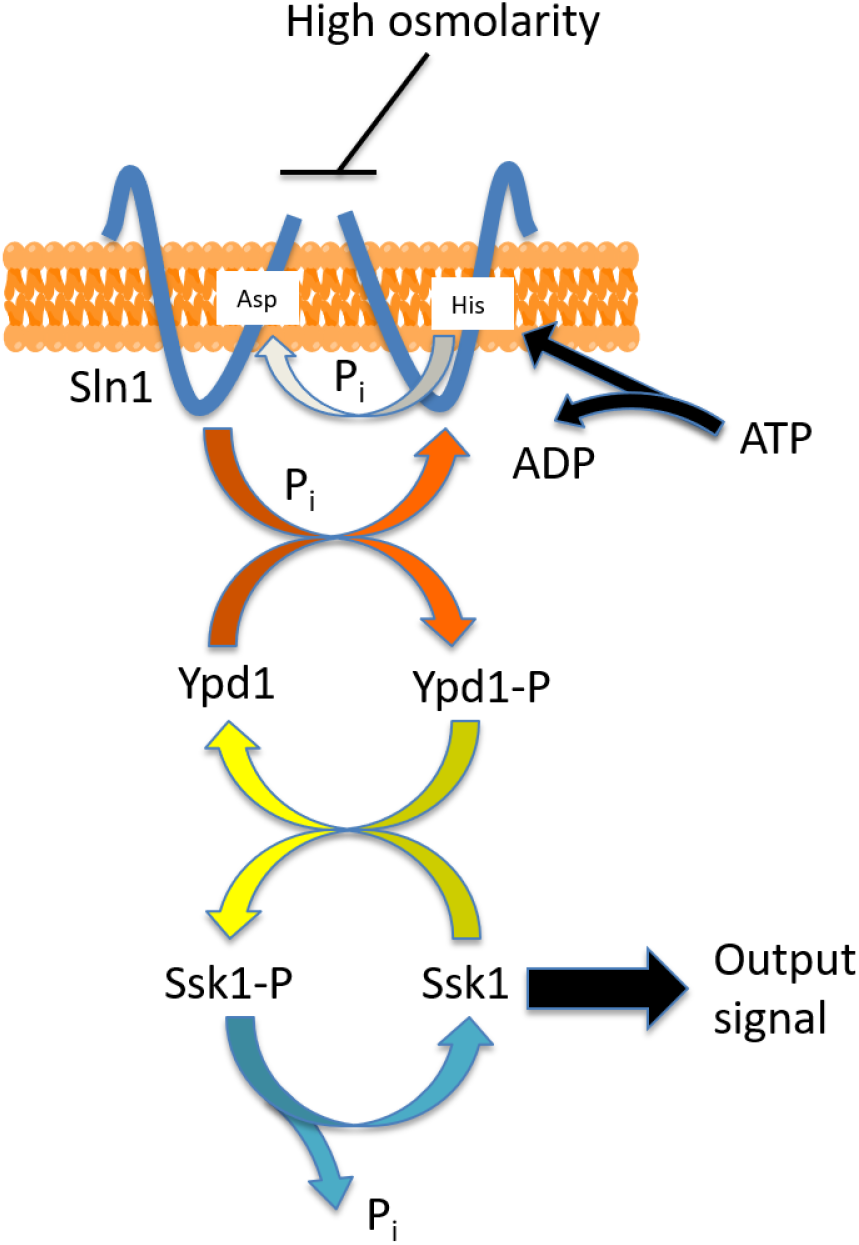

Under normal circumstances, the transmembrane protein Sln1, which is present as a dimer, autophosphorylates at a histidine residue (consuming ATP). The phosphate group is then transferred to an aspartate residue of Sln1. Thereafter, the phosphate is transferred to the protein Ypd1 and finally to the protein Ssk1. Ssk1 is continuously dephosphortylated to give an output signal. The signalling pathway is inhibited by an increase is osmolarity outside of the cell (Klipp et al. 2009). If we let *A* represent Sln1, *B* represent Ypd1, *C* represent Ssk1 and *XP* represent the phophorylated form of protein *X*, then the above network can be represented by the below schematic (Klipp et al. 2009):

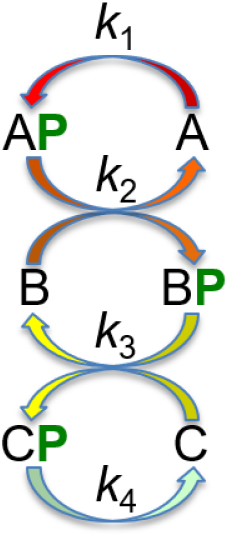

where each *k_i_* represents the rate constant for the relevant phosphorylation/dephosphorylation reaction.

The above graphic allows us to arrive at the following equations to describe the temporal behavior of the phosphorelay system:

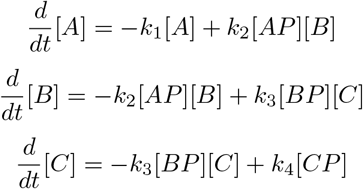

Moreover, under the generally accepted assumption that the degradation and production of proteins occurs on a time scale that far exceeds that of phosphorylation events, we have the following conservation relationships (Klipp et al. 2009):

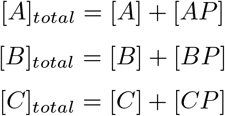

where [*A*]_*total*_, [*B*]_*total*_ and [*C*]_*total*_ are constants. Differentiating, we have:

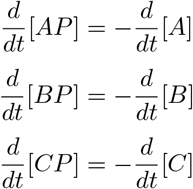

Given this model of the phosphorelay system, the question we desire to answer is as follows: given initial concentrations of the three proteins {[*A*]*_i_*, [*B*]_*i*_, [*C*]*_i_*} and target concentrations of the three proteins {[*A*]_*t*_, [*B*]_*t*_, [*C*]_*t*_}, what are the values {*k*_1_, *k*_2_, *k*_3_, *k*_4_} that result in the proteins having the target concentrations at steady state when the system is allowed to equilibrate from the initial concentrations? This formulation assumes that no information is known about the rate constants and that initial and target concentrations can be determined experimentally, which is often the case in practice (Raue et al. 2013). The problem formulation could be altered depending on the information that is known or that can be determined experimentally.

#### Defining Optimus Inputs

Having outlined how the behaviour of the phosphorelay system can be modelled using a system of differential equations, we can now proceed with defining input parameters for Optimus. We will create a variable state that will be a numeric vector holding the names and initial concentrations of all species in the network. For this tutorial, we will choose [*A*]*_i_* = [*B*]_*i*_ = [*C*]*_i_* = 100 and [*AP*]*_i_* = [*BP*]*_i_* = [*CP*]_*i*_ = 0. Note that the units are arbitrary and that the total sum of units across this vector will remain constant throughout the simulation of the dynamics of the phosphorelay system.

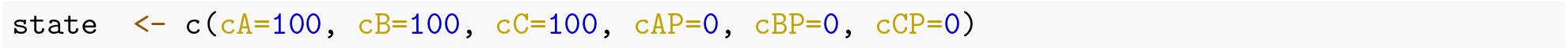

Next, we will create a variable target which will be a numeric vector holding the names and target concentration of all species in the network. We will arbitrarily choose target values of [*A*]_*t*_ = 90, [*B*]_*t*_ = 20, [*C*]_*t*_ = 70, [*AP*]_*t*_ = 10, [*BP*]_*t*_ = 80 and [*CP*]_*t*_ = 30. Note that the chosen target values must be consistent with the above defined conservation equations, meaning we must have [*X*]_*i*_ + [*XP*]_*i*_ = [*X*]_*t*_ + [*XP*]_*t*_, ∀ *X* ∈ {*A, B, C*}.

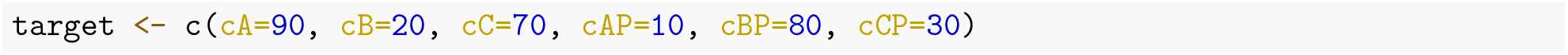

In order to determine the steady state behavior of the ODE system, we will employ the function `ode()` from the R library `deSolve` (this function interfaces with the Fortran library typically used to solve systems of differential equations). This function requires as input a function that describes the dynamics of the ODE system. We will call this function `model()`. At a high level, `model()` will simply define the equations derived in the previous section that describe the network. It should contain equations that use the objects with the names specified within state above, and should have equations that assign the outcomes to new objects that have the same order and names as specified in state, but with “d” at the beginning (a more detailed description of the requirements of `model()` can be found in the R documentation of `ode()`).

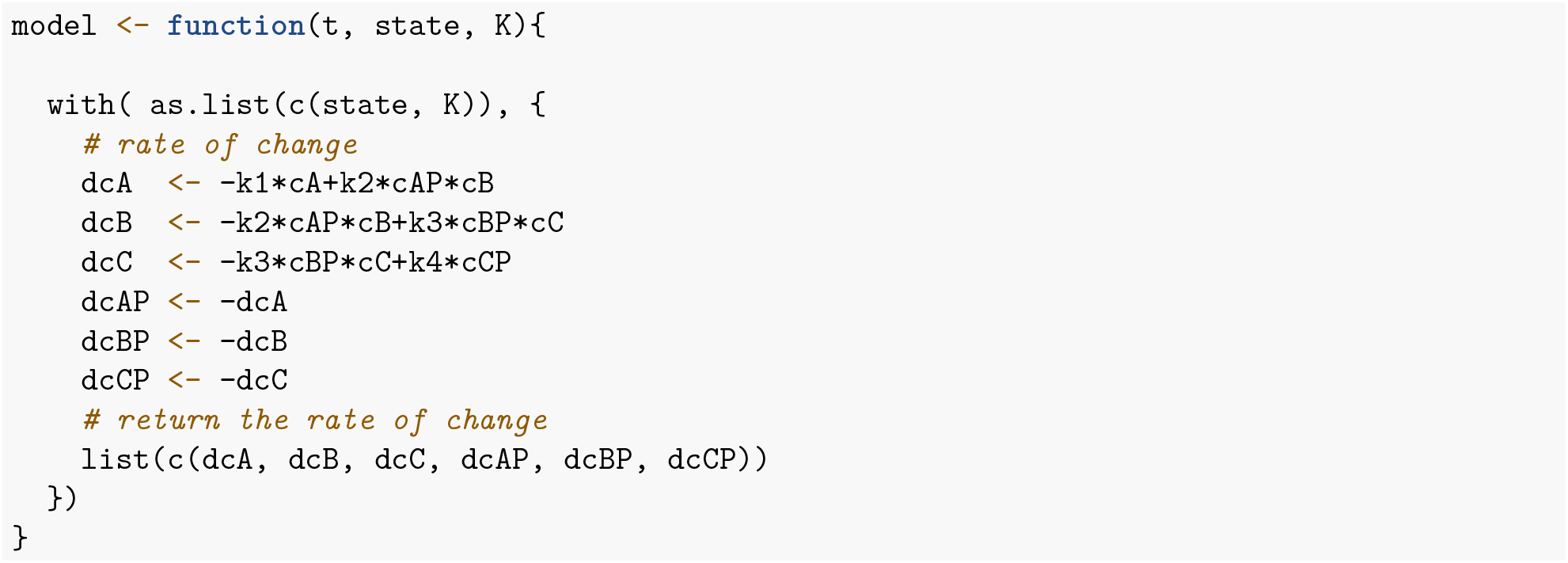

The variables `state` and `target`, and the function `model()` should be stored as entries in a list `DATA` which will be given to the functions `m()` and `u()` as inputs.

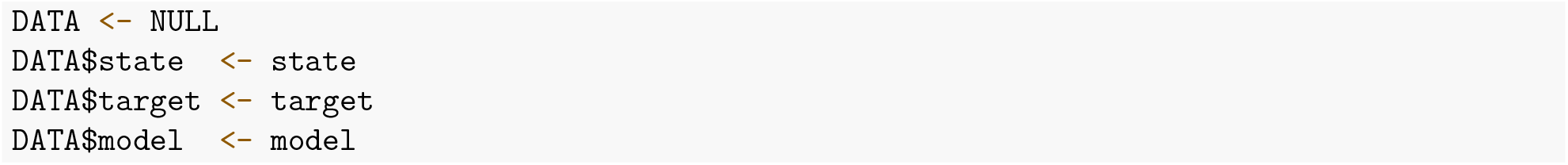

We will make `K` be a numeric vector holding the set of rate constants {*k*_1_, *k*_2_, *k*_3_, *k*_4_}. We will (arbitrarily) initialize each rate constant to have value 1.0.

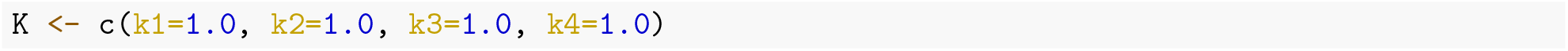

The function `m()` will take as input the vector `K` of rate constants and the list `DATA`. It will return an object `O` that contains the concentrations of the six species in the network when the system is simulated from the initial state specified in `DATA` using the `K` rate constants for 10 time steps. Note that it is not necessarily guaranteed that the system will reach a steady state after 10 time steps; the number of time steps was chosen such that the optimisation procedure would terminate within 1-2 hours in this example. `m()` will call the function `ode()` from the library `deSolve`, so we must first ensure that `deSolve` is installed.

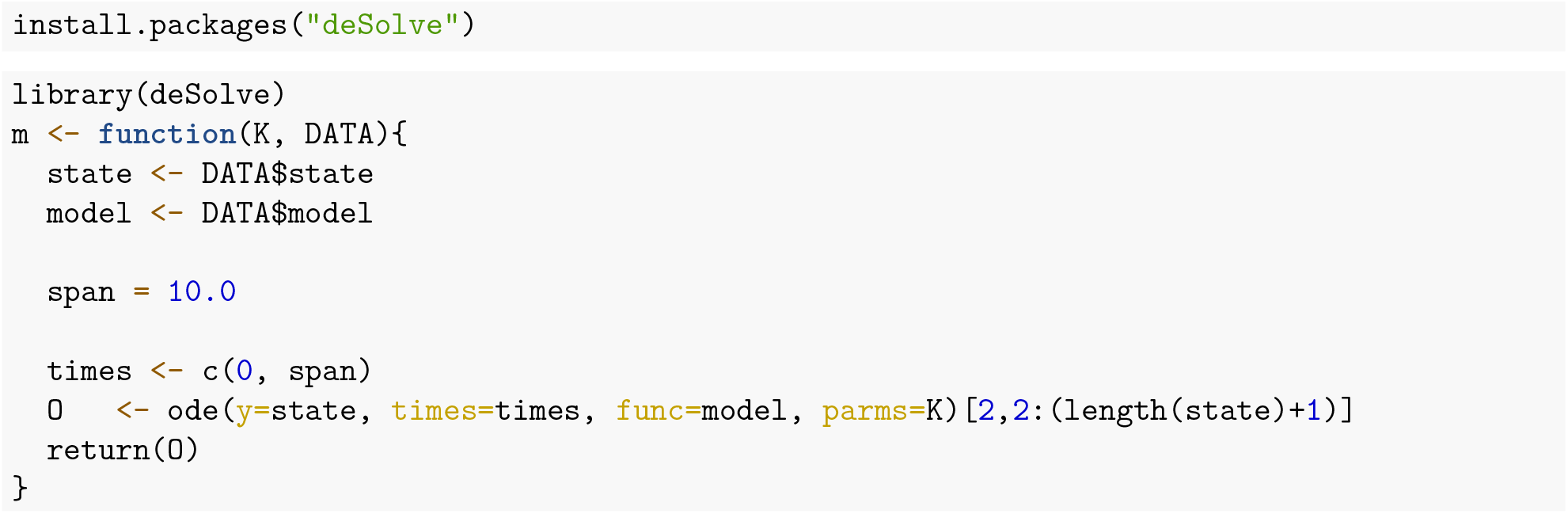

Recall that the function `u()` must return an energy `E` and a quality `Q` of the candidate solution. Here, `u()` will set both `E` and `Q` to be the RMSD between the steady state concentrations of the network corresponding to the current set of rate constants `K`, as determined by `m()`, and the target concentrations.

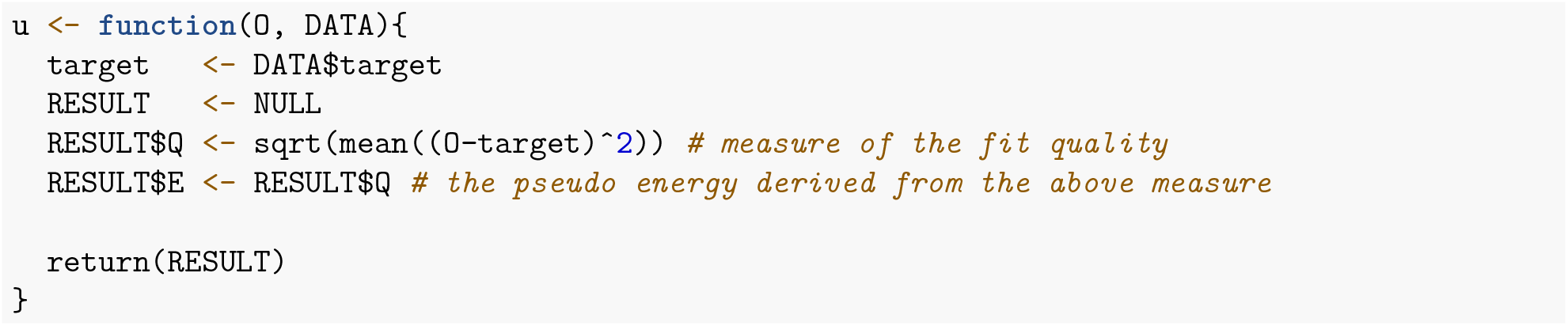

The final mandatory input to Optimus that must be defined is the alteration function `r()`. Just as in **Tutorial 1**, for each snapshot of `K`, we shall randomly select one of its four coefficients, then either increment or decrement (chosen randomly) it by 0.0002, returning the altered set of coefficients. Since we are dealing with rate constants in this case, if ever `r()` were to make an entry in *K* negative, that entry will automatically be set to 0.

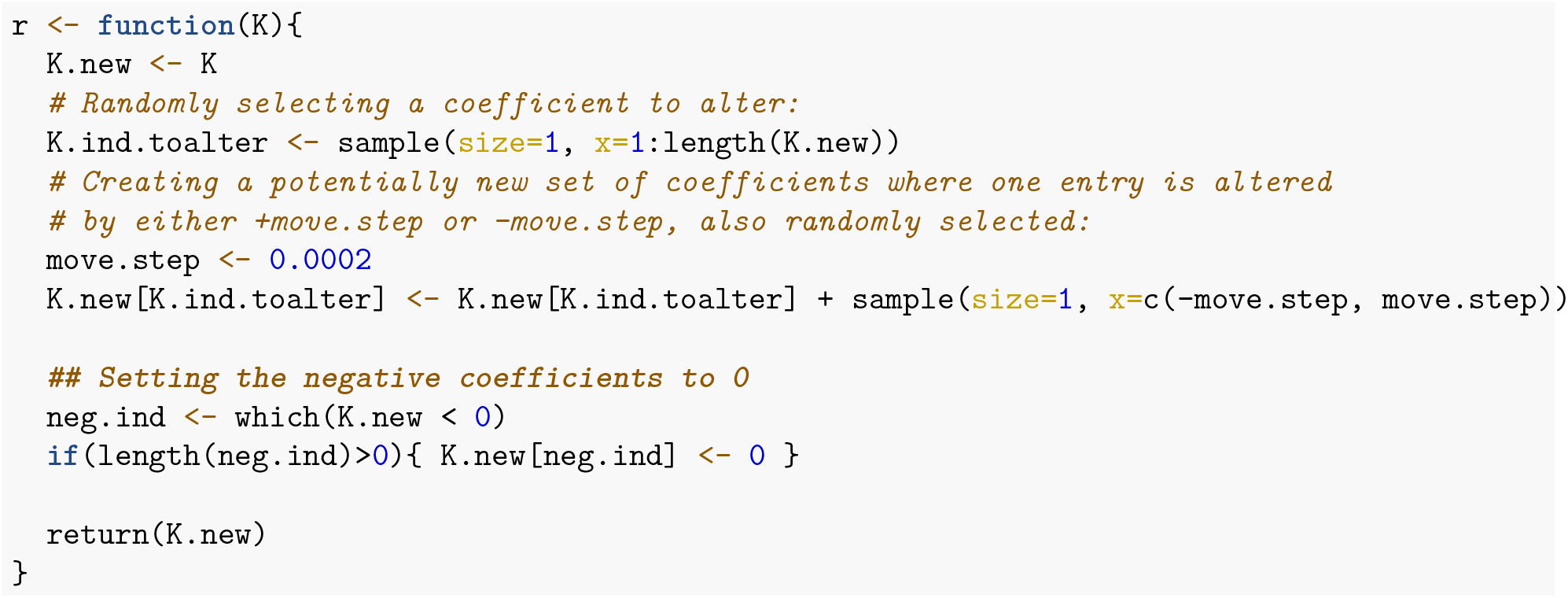

#### Exploring the System Dynamics

Before calling Optimus to solve this problem, let us first simulate the system of ODEs from the chosen initial state using a few sets of arbitrary rate constants to become familiar with how the system evolves. The below graphs illustrate the evolution of the system for 50 time steps for the rate constants {*k*_1_ = 1.0, *k*_2_ = 1.0, *k*_3_ = 1.0, *k*_4_ = 1.0}:

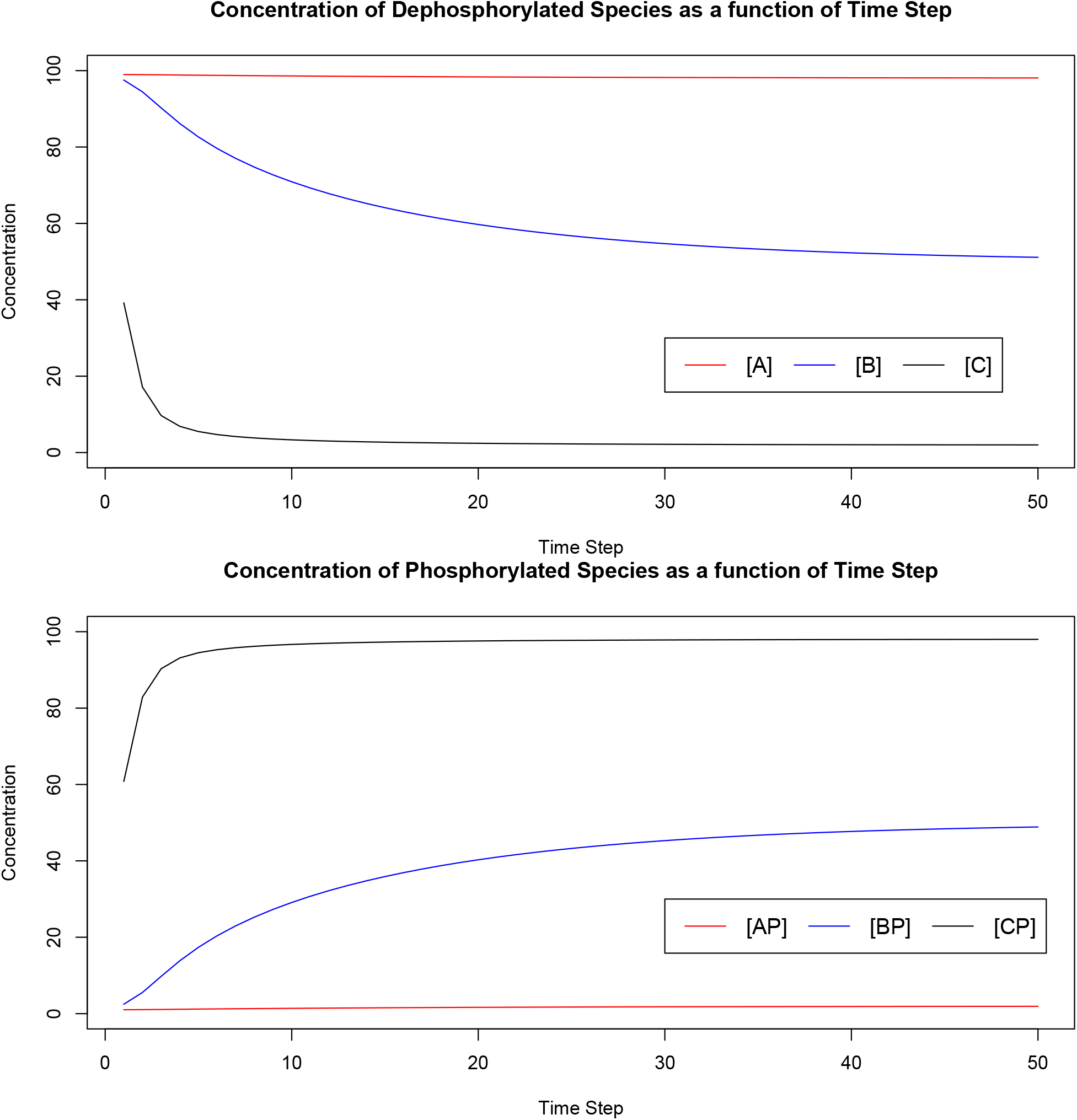

The table below summarises the initial and final concentrations of the various species when the system is simulated for 50 time steps using the rate constants {*k*_1_ = 1.0, *k*_2_ = 1.0, *k*_3_ = 1.0, *k*_4_ = 1.0}:

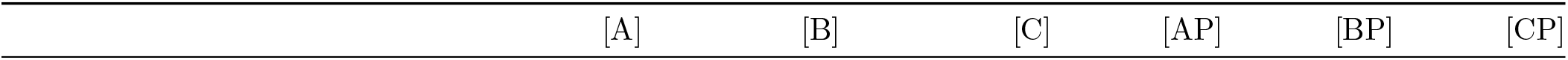

**Table 11:** System summary for k1 = k2 = k3 = k4 = 1.0.

|  | [A] | [B] | [C] | [AP] | [BP] | [CP] |
| --- | --- | --- | --- | --- | --- | --- |
| Initial | 100.00000 | 100.00000 | 100.000000 | 0.000000 | 0.00000 | 0.00000 |
| Final (after 50 time steps) | 98.08145 | 51.12118 | 2.004924 | 1.918549 | 48.87882 | 97.99508 |

If instead we use the set of rate constants {*k*_1_ = 1.5, *k*_2_ = 0.5, *k*_3_ = 1.0, *k*_4_ = 1.0}, the system evolves as follows:

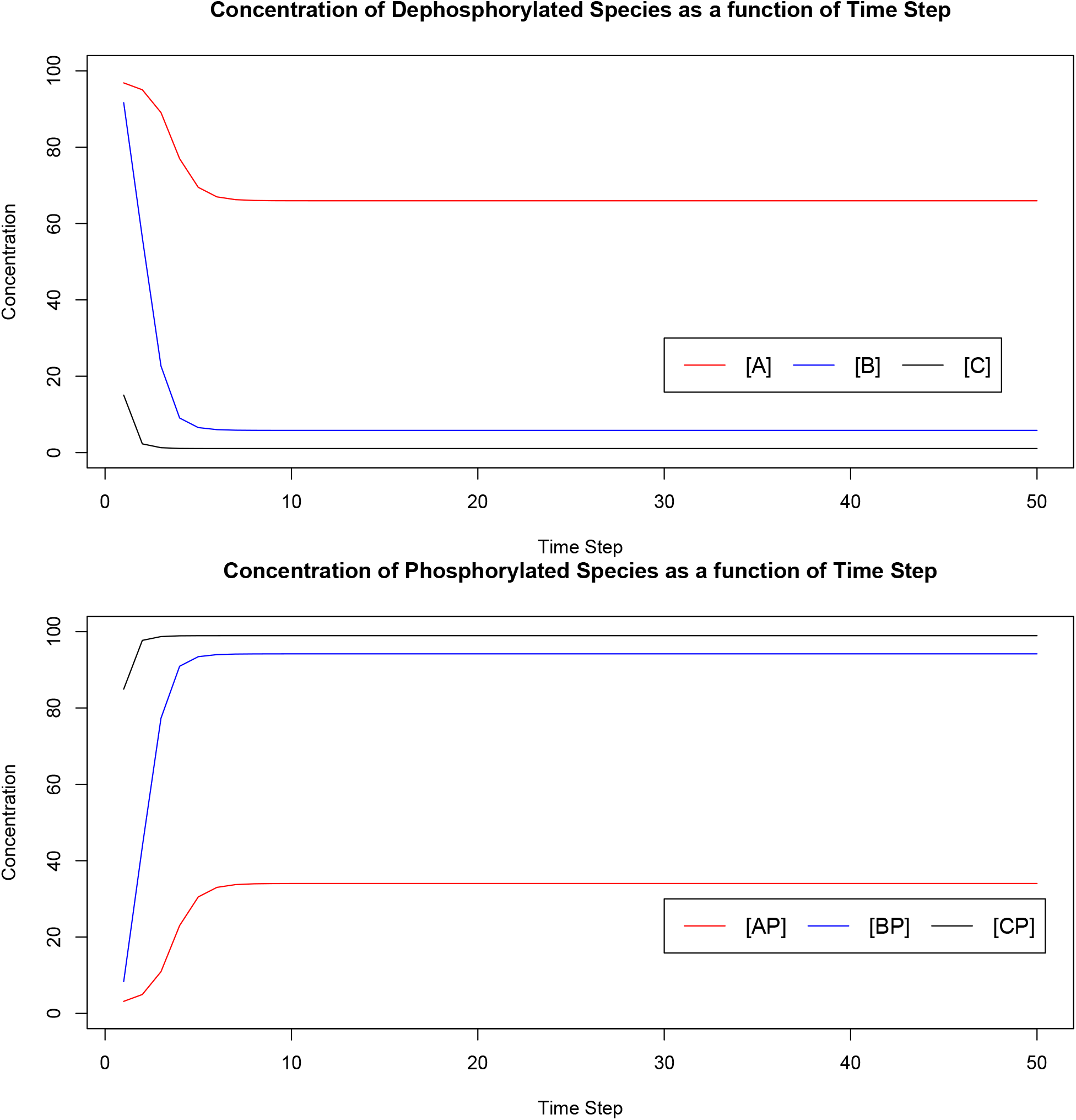

The table below summarizes the initial and final concentrations of the various species when the system is simulated for 50 time steps using the rate constants {*k*_1_ = 1.5, *k*_2_ = 0.5, *k*_3_ = 1.0, *k*_4_ = 1.0}:

**Table 12:** System summary for k1 = 1.5, k2 = 0.5, k3 = k4 = 1.0.

|  | [A] | [B] | [C] | [AP] | [BP] | [CP] |
| --- | --- | --- | --- | --- | --- | --- |
| Initial | 100.00000 | 100.000000 | 100.000000 | 0.00000 | 0.00000 | 0.00000 |
| Final (after 50 time steps) | 65.96628 | 5.814787 | 1.050583 | 34.03372 | 94.18521 | 98.94942 |

#### Acceptance Ratio Simulated Annealing Optimus Run

We will now call Acceptance Ratio Simulated Annealing Optimus to solve our problem. Similarly to **Tutorial 2**, we will execute 200 000 optimisation iterations and perform 2 annealing cycles. We will set `DUMP.FREQ` to have a value of 100 000.

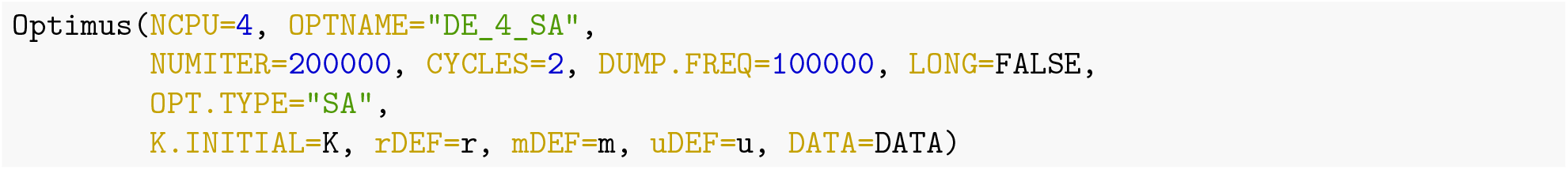

**Table 13:** 4-core Acceptance Ratio Simulated Annealing results from Optimus run.

|  | E (RMSD) | K1 | K2 | K3 | K4 |
| --- | --- | --- | --- | --- | --- |
| CPU 1 | 0.0012516 | 0.7974 | 0.3586 | 0.0128 | 2.3886 |
| CPU 2 | 0.0017556 | 0.7850 | 0.3532 | 0.0126 | 2.3512 |
| CPU 3 | 0.0013625 | 0.8098 | 0.3642 | 0.0130 | 2.4262 |
| CPU 4 | 0.0012516 | 0.7974 | 0.3586 | 0.0128 | 2.3886 |

Of the 4 optimisation replicas, CPU 1 and 4 find the best set of rate constants, {*k*_1_ = 0.7974, *k*_2_ = 0.3586, *k*_3_ = 0.0128, *k*_4_ = 2.3886}. This set of rate constants results in an RMSD (after 10 iterations) of 0.0012516. Let us simulate how the system evolves according to these rate constants for 10 time steps:

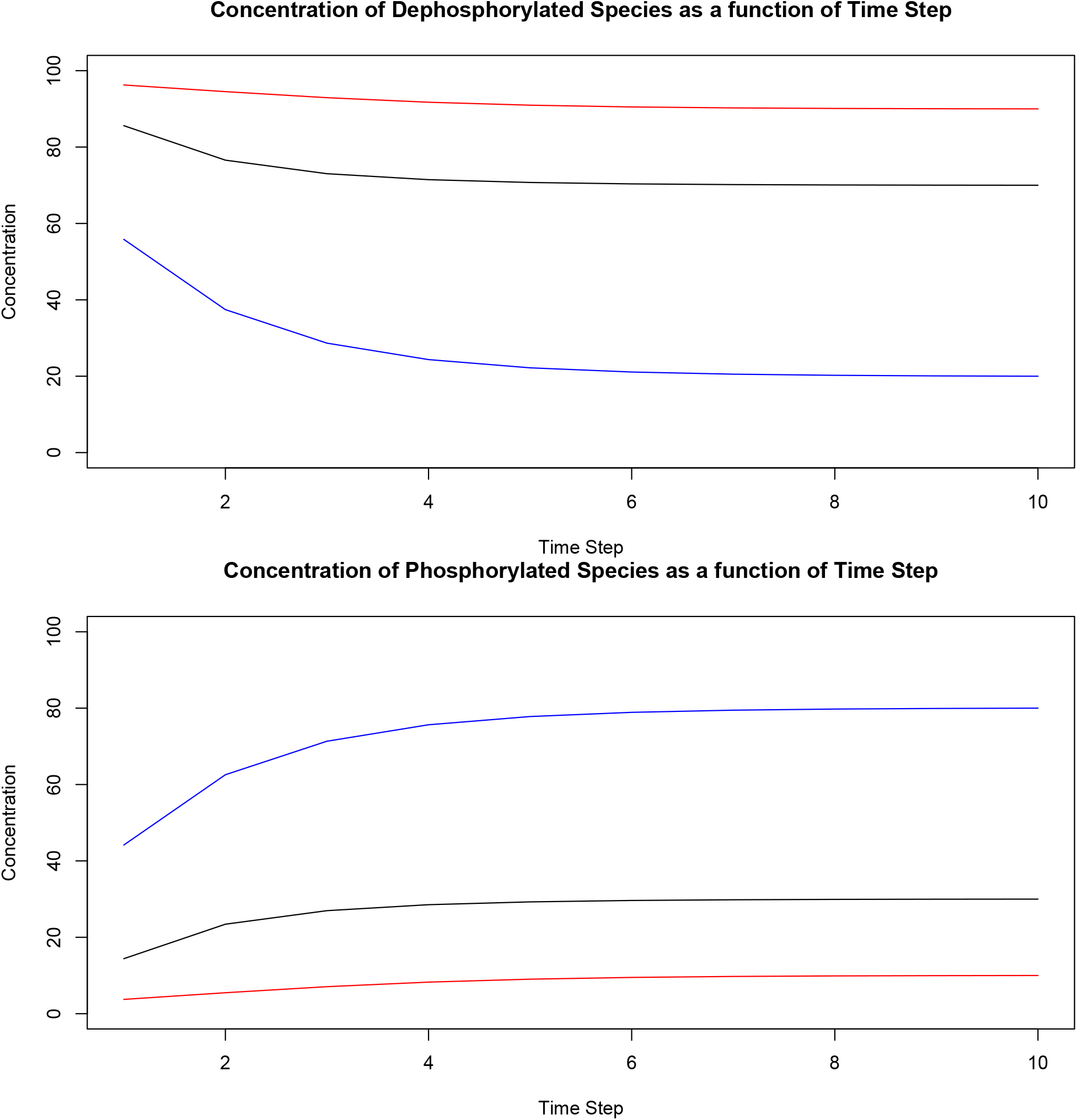

The table below summarizes the initial and final concentrations of the various species when the system is simulated for 10 time steps using the rate constants {*k*_1_ = 0.7974, *k*_2_ = 0.3586, *k*_3_ = 0.0128, *k*_4_ = 2.3886}:

**Table 14:** System summary for k1 = 0.7974, k2 = 0.3586, k3 = 0.0128, k4 = 2.3886.

|  | [A] | [B] | [C] | [AP] | [BP] | [CP] |
| --- | --- | --- | --- | --- | --- | --- |
| Initial | 100.00000 | 100.00000 | 100.00000 | 0.00000 | 0.00000 | 0.00000 |
| Final (after 10 time steps) | 89.99814 | 20.00081 | 69.99924 | 10.00186 | 79.99919 | 30.00076 |

As can be seen in the above table, the concentration of the species in the system after 10 time steps are very close to the target values [*A*]_*t*_ = 90, [*B*]_*t*_ = 20, [*C*]_*t*_ = 70, [*AP*]_*t*_ = 10, [*BP*]_*t*_ = 80 and [*CP*]_*t*_ = 30. As alluded to earlier, it is possible that the system has not reached a steady state after 10 time steps, however the above graphs suggest that the concentrations after 10 steps are already extremely close to, if not equal to, steady state values. The optimisation process could be re-executed after increasing the value of the parameter `span` in the function `m()` to simulate the system for a larger number of time steps.

#### Acceptance Ratio Replica Exchange Optimus Run

Let us now examine how replica exchange Optimus using 12 cores performs on this task. We will use 200 000 optimisation iterations and set `STATWINDOW` to 50, similarly to **Tutorials 2** and **3**.

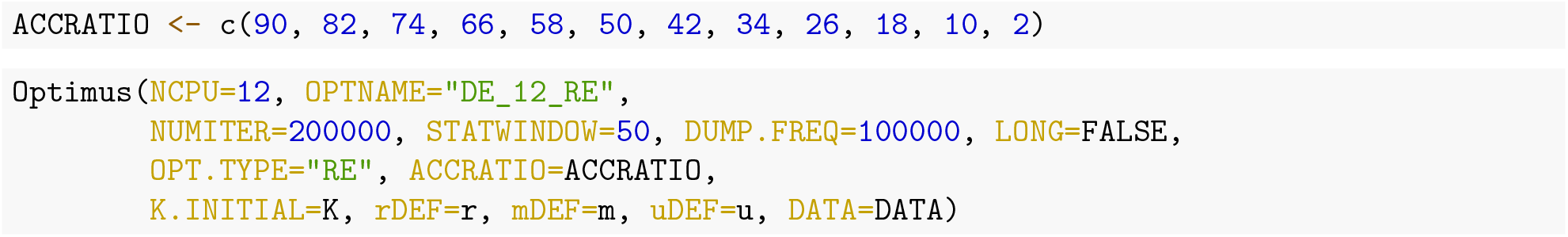

**Table 15:**
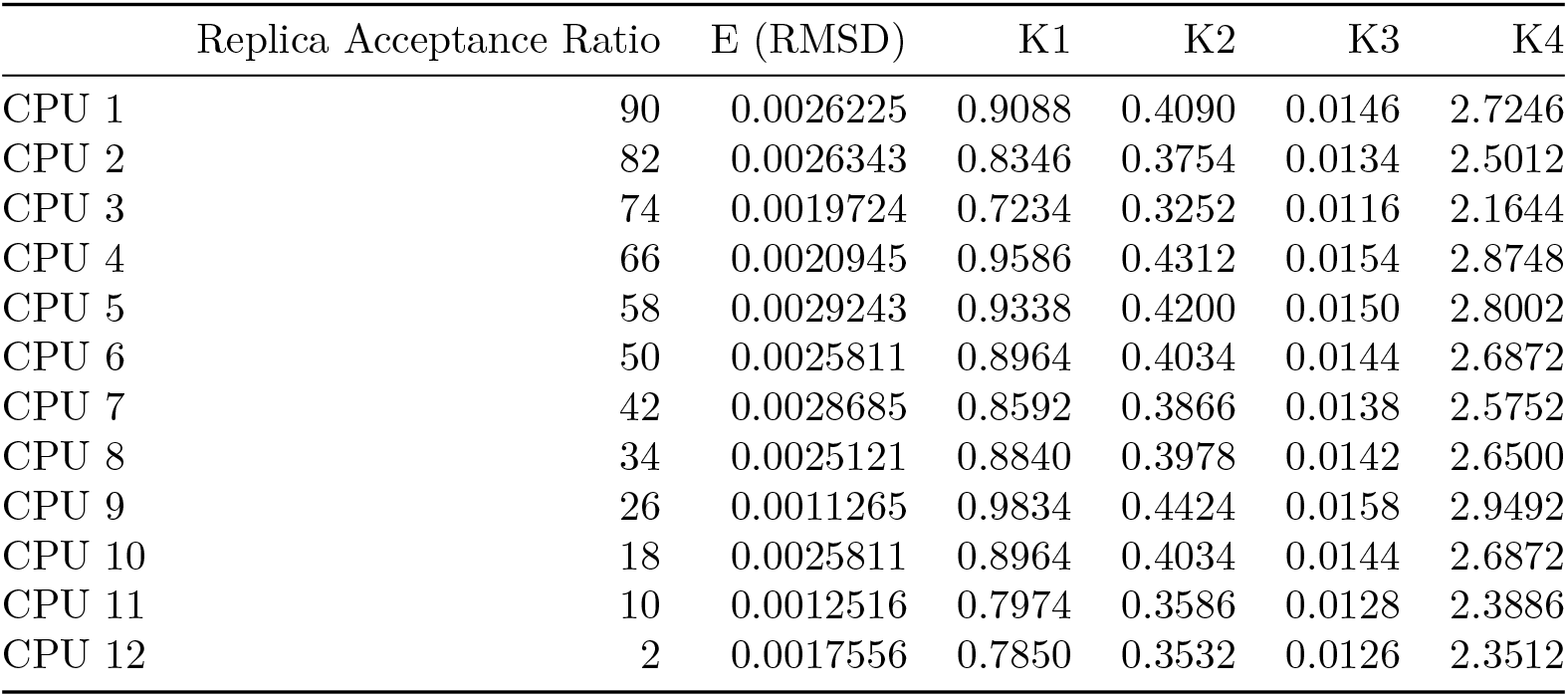
12-core Acceptance Ratio Replica Exchange results from Optimus run.

Of the 12 optimisation replicas, CPU 9 (26% acceptance ratio) finds the best set of rate constants, {*k*_1_ = 0.9834, *k*_2_ = 0.4424, *k*_3_ = 0.0158, *k*_4_ = 2.9492}. This set of rate constants results in an RMSD (after crude 10 iteration limit) of 0.0011265, which is lower than the RMSD of the solution found by Acceptance Ratio Simulated Annealing Optimus (0.0012516) done with the use of more modest computational resources. Let us simulate how the system evolves according to these rate constants for 10 time steps:

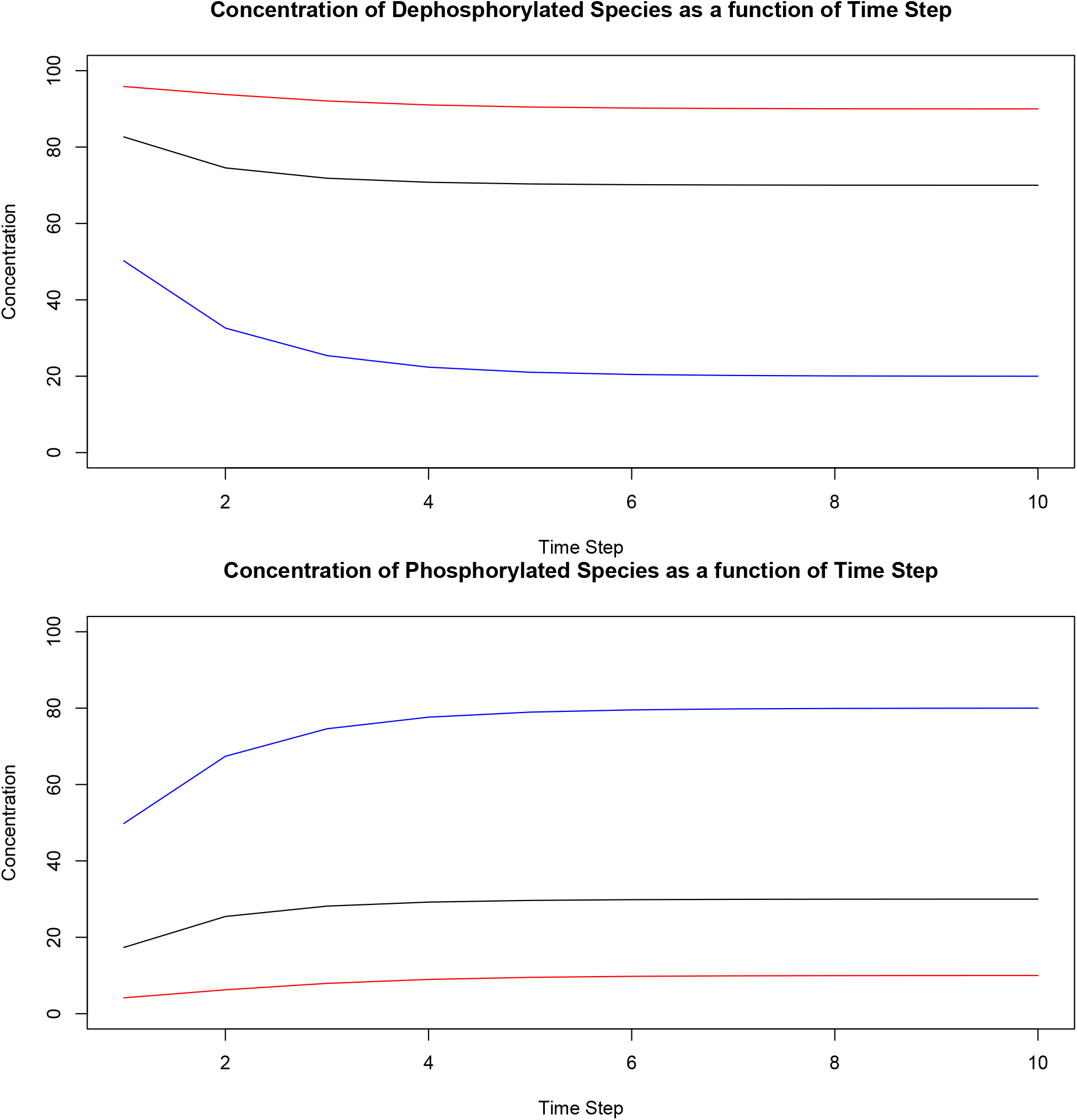

The table below summarises the initial and final concentrations of the various protein species when the system is simulated for 10 time steps using the rate constants {*k*_1_ = 0.9834, *k*_2_ = 0.4424, *k*_3_ = 0.0158, *k*_4_ = 2.9492}:

**Table 16:** System summary for k1 = 0.9834, k2 = 0.4424, k3 = 0.0158, k4 = 2.9492.

|  | [A] | [B] | [C] | [AP] | [BP] | [CP] |
| --- | --- | --- | --- | --- | --- | --- |
| Initial | 100.00000 | 100.00000 | 100.00000 | 0.00000 | 0.00000 | 0.00000 |
| Final (after 10 time steps) | 89.99807 | 19.99983 | 70.00022 | 10.00193 | 80.00017 | 29.99978 |

Here again, we see that the protein concentrations after 10 time steps are remarkably close to the target values [*A*]_*t*_ = 90, [*B*]_*t*_ = 20, [*C*]_*t*_ = 70, [*AP*]_*t*_ = 10, [*BP*]_*t*_ = 80 and [*CP*]_*t*_ = 30 and the graphs suggests that these concentrations have either converged or are very close to converging to the steady state.

#### Summary

We have seen how Optimus can be employed to recover rate constants for a system of coupled ODEs that describes a biological pathway. Given an initial state of the system and a target state, both Acceptance Ratio Simulated Annealing (SA) Optimus and Replica Exchange (RE) Optimus found a set of rate constants that resulted in the desired system behaviour upon simulation of the system. In the current setup in this tutorial, RE slightly outperformed SA, both however are not directly comparable unless their allocated resources and times are equalised.

**Table 17:** Summary of protein concentrations after 10 time steps.

|  | [A] | [B] | [C] | [AP] | [BP] | [CP] |
| --- | --- | --- | --- | --- | --- | --- |
| Target | 90.00000 | 20.00000 | 70.00000 | 10.00000 | 80.00000 | 30.00000 |
| Optimus (AR Simulated Annealing) | 89.99814 | 20.00081 | 69.99924 | 10.00186 | 79.99919 | 30.00076 |
| Optimus (AR Replica Exchange) | 89.99807 | 19.99983 | 70.00022 | 10.00193 | 80.00017 | 29.99978 |

**Table 18:** Summary of recovered rate constants.

|  | E (RMSD) | K1 | K2 | K3 | K4 |
| --- | --- | --- | --- | --- | --- |
| Optimus (AR Simulated Annealing) | 0.0012516 | 0.7974 | 0.3586 | 0.0128 | 2.3886 |
| Optimus (AR Replica Exchange) | 0.0011265 | 0.9834 | 0.4424 | 0.0158 | 2.9492 |

### Tutorial 5: Constrained Shuffling of Genomic Contacts to Form a Control Set

#### Problem Statement

Here, we shall consider a problem from the field of 3D genome organisation, to exemplify how Optimus can optimise a specific constrained shuffling task given a certain set of conditions.

Mammalian genomes are packaged into a complex three-dimensional (3D) structure inside the nucleus. They are efficiently folded bringing DNA regions near or far from each other into contact, facilitating the tight gene regulation.

Here, we take a set of 734 pairs of *i* and *j* 40-kilobase DNA regions that are known to be in contact inside a cell. In our set, a 40-kb DNA region is represented as a single integer that is equal to its end position divided by the length of the region, which is 40 kb. For instance, a region, with start and end positions at the 1^*st*^ and 40000^*th*^ nucleotides, respectively, is denoted as 1 (40000^*th*^ base /40000 bases = 1). This simplifies the notation for a contact to a pair of positive integers as shown below:

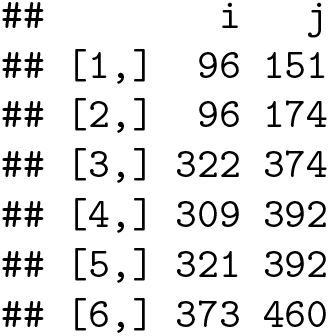

where *i* ∈ *Z*^+^ and *j* ∈ *Z*^+^.

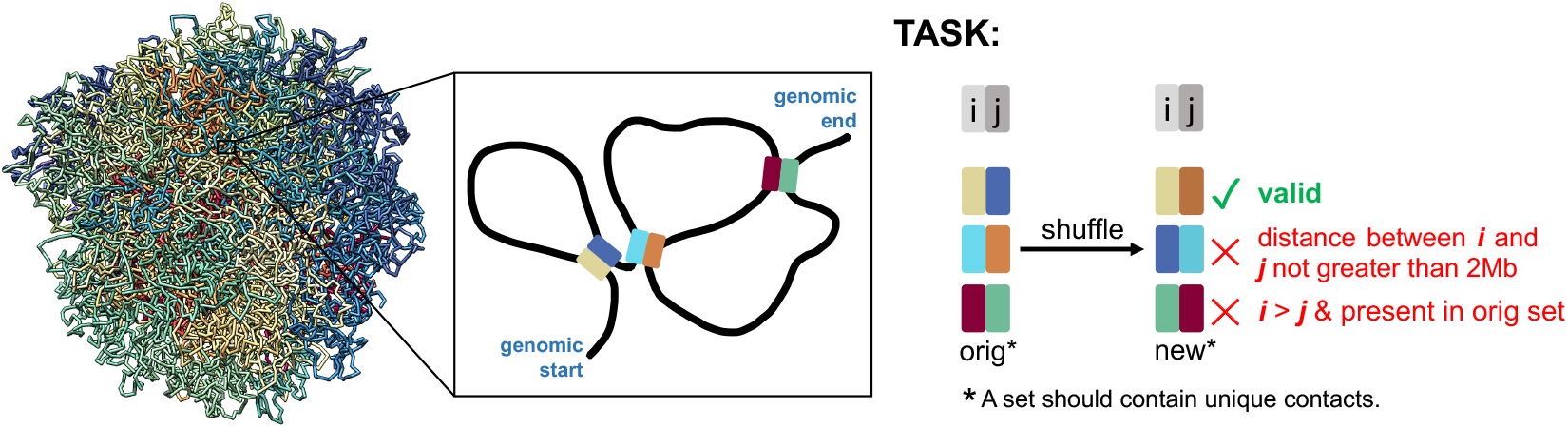

Take note that our set of contacts comply with the following criteria.

1. The set only contains **unique, long-range contacts**, with *i* and *j* regions always separated by greater than 2 megabases (Mb). In terms of the integer identifiers, the threshold is 50 (2*Mb*/40*kb* = 50), such that (*j* – *i*) > 50.
2. *i* is set to be always less than *j*. This avoids duplicate contacts in the set definition, such as

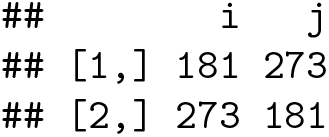
3. A region can take part in more than one contact (as in region 96 in rows 1 and 2 in the table above), given that the contacts it forms comply with the two aforementioned criteria.

The task herein is to take the DNA regions from the original set (shown above) and to specifically shuffle them to produce a new set of contacts. This is useful for a research on genomic 3D contacts requiring good quality of negative controls that still have the constraints that the original true data possess. The challenge here is to maximise the number of new control contacts that we can devise and that are valid, meaning that they A) still satisfy the aforementioned 3 criteria, and B) are not present in the original set.

#### Defining Optimus Inputs

We can solve this highly constrained shuffling problem using Optimus by doing the following preparations. Keeping the original order of *i*’s in the loaded real set of contacts (referred hereafter as `IJ.ORIG`), we can randomly shuffle all *j*’s in the set once, without replacement. The output will be the object `K` to be supplied to Optimus for the optimisation.

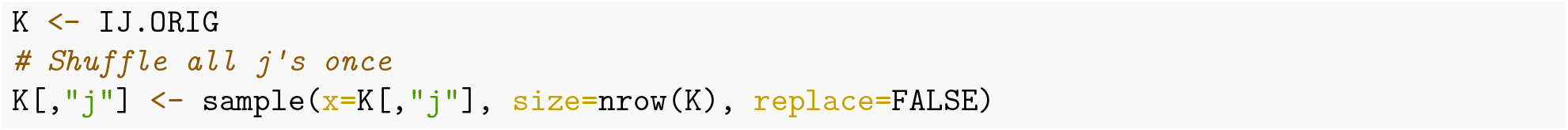

The `m()` function should then operate on the object `K`, which, to reiterate, is derived from the original set of contacts with all *j*’s shuffled once (see above). It can also accept a supplementary object `DATA` that holds data necessary for the model calculations and outcome comparison. Output of `m()` is the observable object `O` that is the percentage of contacts not satisfying the aforementioned criteria or are already in the original set. To get O, `m()` will require a) the original set of contacts (`IJ.ORIG`), b) number of contacts, and c) gap threshold set between contacting regions in terms of positive integer (50). These are first stored in the `DATA` object then passed to `m()`.

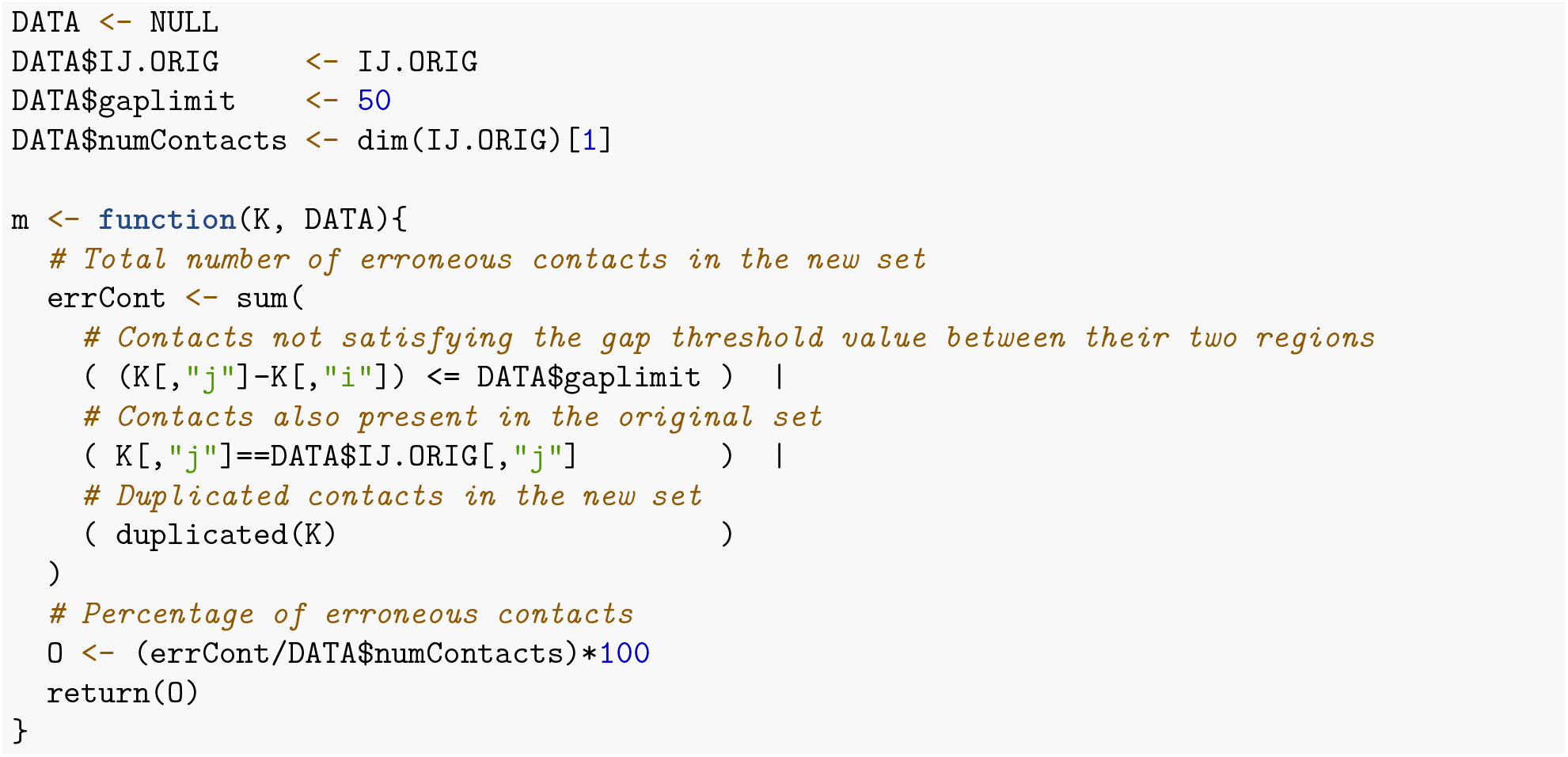

Dictated by the `u()` function, the error percentage `O`, will be directly used as the pseudo energy `E`, since we do want to minimise this value. Meanwhile, the quality `Q` for the given snapshot of `K`, which we want to increase, can be the negative of `O`.

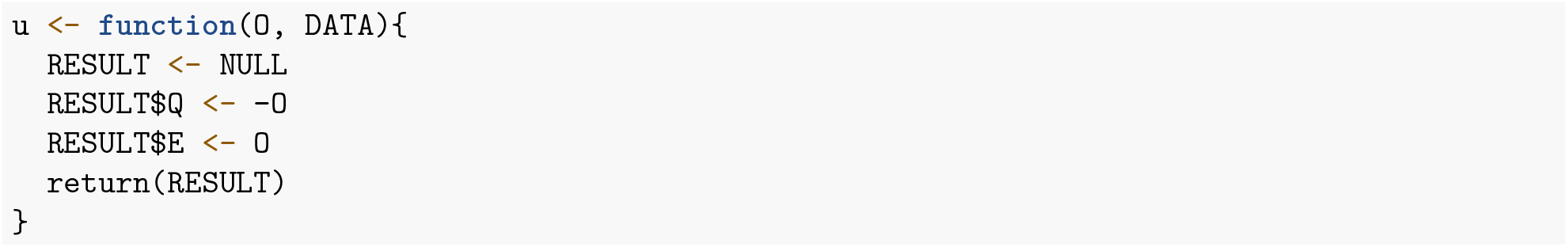

Finally, `r()`, which defines how object `K` will be altered at every step, takes two *j*’s and then swaps them, without changing the order of their partner *i*’s.

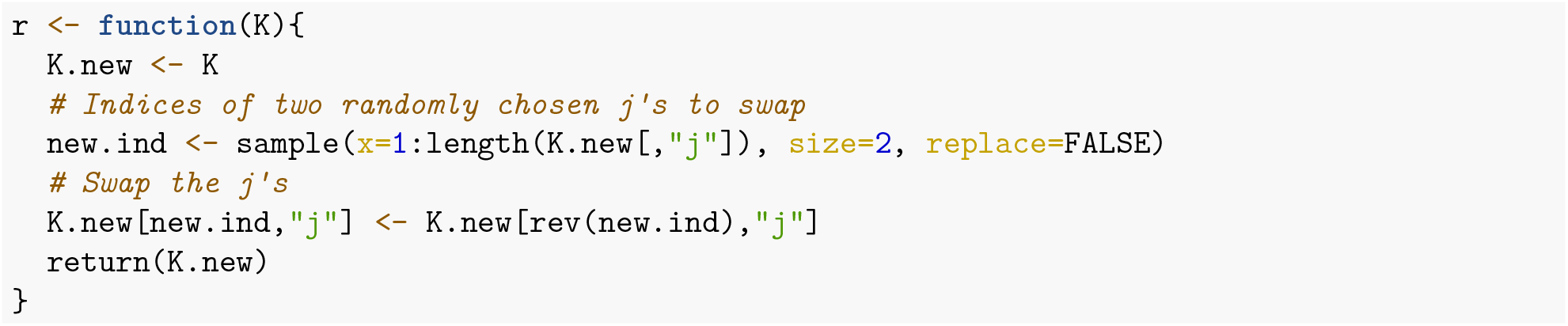

#### Acceptance Ratio Simulated Annealing Optimus Run

We first use the Acceptance Ratio Simulated Annealing scheme in Optimus to solve the problem. As in **Tutorials 2** and **4**, Optimus is set to do 200 000 steps, 2 annealing cycles, and is to produce 4 independent replicas of the same simulation.

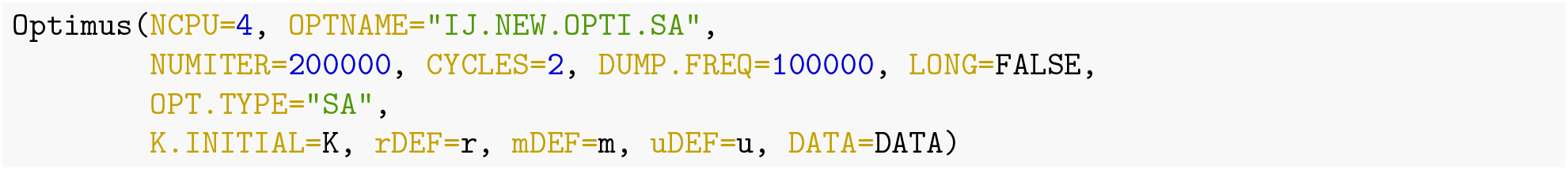

Note that if additional data is required by `m()` or `u()`, that object should be assigned to `DATA` argument of `Optimus()`, otherwise `DATA` defaults to `NULL`.

The 4 new sets obtained as optimums from each of the 4 replicates of independent simulated annealing runs have the following error percentages:

**Table 19:**
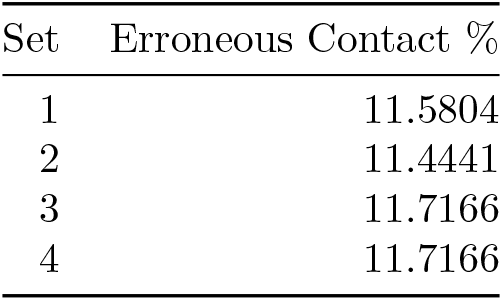
4-core Acceptance Ratio Simulated Annealing results from Optimus run.

| Set | Erroneous Contact % |
| --- | --- |
| 1 | 11.5804 |
| 2 | 11.4441 |
| 3 | 11.7166 |
| 4 | 11.7166 |

#### Acceptance Ratio Replica Exchange Optimus Run

Now, we can use the Acceptance Ratio Replica Exchange mode of Optimus using 12 cores and the same number of optimisation steps (200 000) as in the Acceptance Ratio Simulated Annealing Optimus run. Similar to **Tutorials 2** to **4**, `STATWINDOW` is set to 50 and the acceptance ratios are as follows:

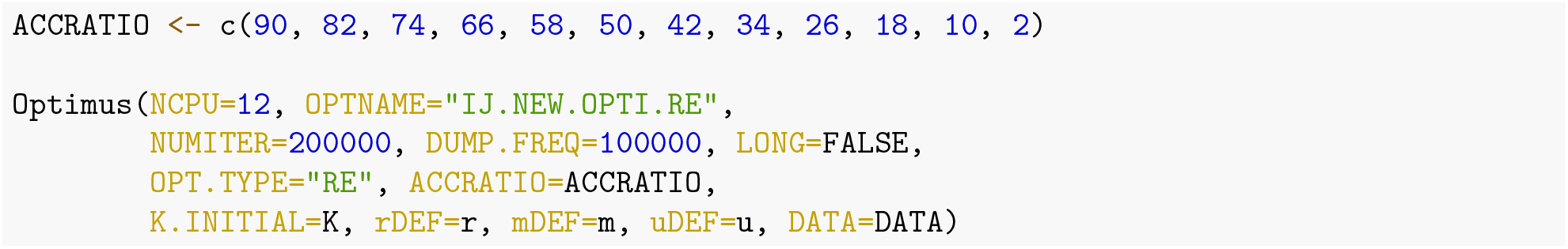

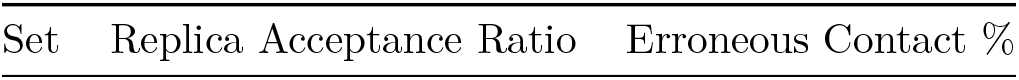

**Table 20:**
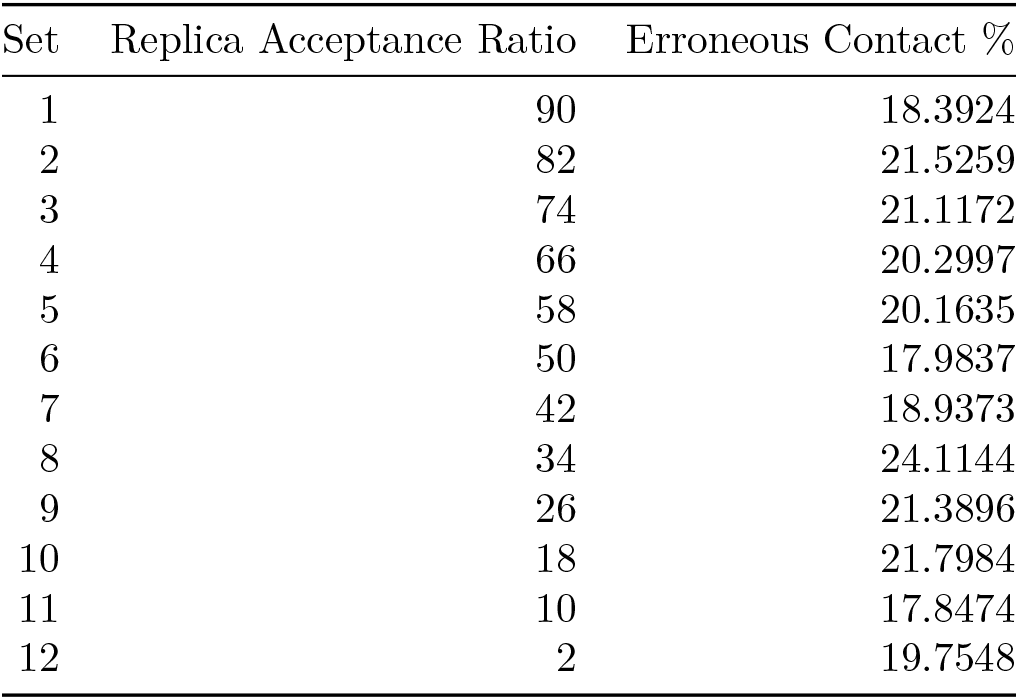
12-core Acceptance Ratio Replica Exchange results from Optimus run.

Notably, in this setup, none of the 12 new sets of contacts from the replica exchange run are better than the sets produced by the Acceptance Ratio Simulated Annealing mode of Optimus.

#### Summary

We have added yet another one to the diverse applications of Optimus, this time by performing a constrained shuffling of pairs of positive integers that, in this case, represent contacts between DNA regions, to form a new set of control contacts. Additionally, Optimus showed to be a platform, where certain restrictions or conditions can easily be incorporated to a given shuffling task, which usually is required for specific research objectives. The table below shows the % of erroneous contacts of the best outputs from the acceptance ratio annealing and replica exchange Optimus runs, with the former producing the better set with 650 valid control contacts out of the 734 total number of the original real contacts.

**Table 21:**
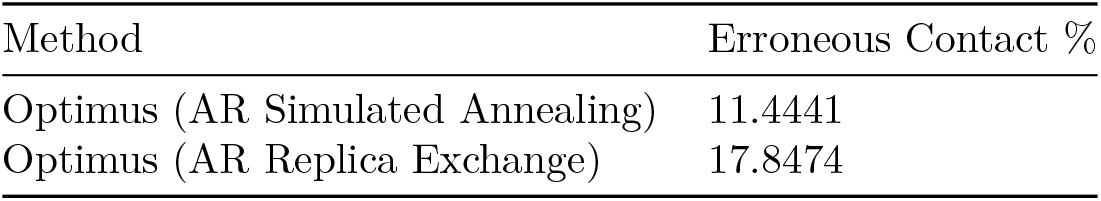
Summary of solutions.

| Method | Erroneous Contact % |
| --- | --- |
| Optimus (AR Simulated Annealing) | 11.4441 |
| Optimus (AR Replica Exchange) | 17.8474 |

### Advanced User Manual

This section will document all possible non-experimental input arguments to Optimus and will describe the output format of Optimus. The code snippet below is the first part of the definition of Optimus, which lists all non-experimental input arguments. We will refer to input arguments without default values as mandatory input arguments and those with default values as optional (with the exception of `K.INITIAL`, which will be considered a mandatory input argument).

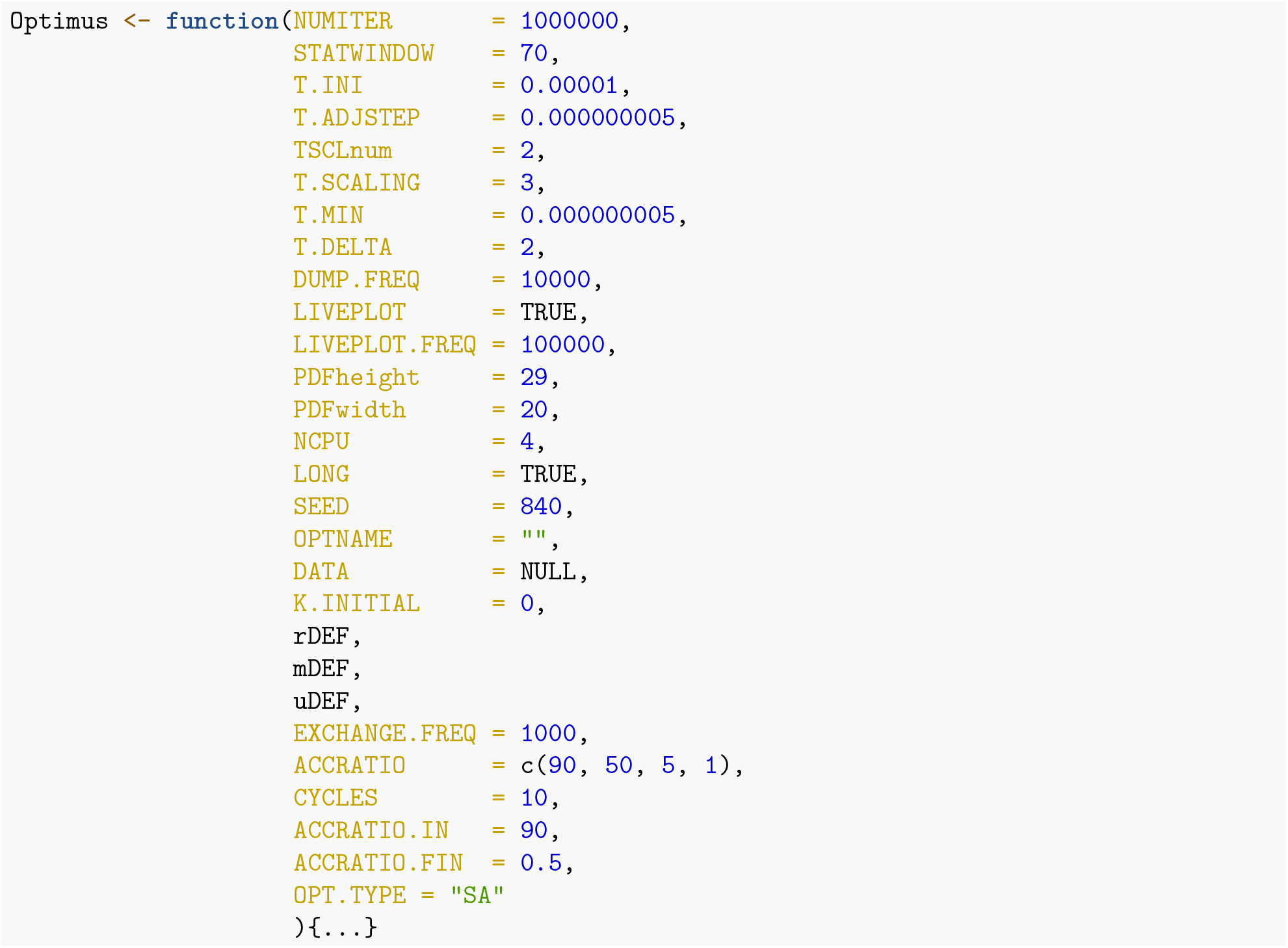

#### Mandatory Input Arguments

All mandatory input arguments have necessarily already been defined in the tutorials. However, we will reiterate these definitions in this section. Optimus has four mandatory inputs, `mDEF, uDEF, rDEF` (referred to as `m()`, `u()` and `r()` respectively in the tutorials) and `K.INITIAL`.

##### mDEF

`mDEF` must be of type closure. `mDEF` should be a function designed to operate on the whole set of parameter snapshot `K` and return the corresponding observable object `O`. Please note, that the size and shape of `K` and `O` are not necessarily to match, depending on the nature of the model used. `mDEF` must necessarily take `K` and `DATA` as input variables, and it must necessarily operate on `K` to produce `O` (operating on `DATA` in optional, see **Tutorial 3** for an illustration).

##### uDEF

`uDEF` must be of type closure. `uDEF` should be a function designed to evaluate the performance of a given snapshot of coefficients `K`. `uDEF` should necessarily take as inputs `O` (the output of `mDEF`) and the variable `DATA`. The output of `uDEF` should have two components, `Q` holding a single number of the quality of the `K` coefficients (also used for plotting), and `E` holding a (pseudo)energy for the given snapshot `K`. It is important that the returned (pseudo)energy value is lower for better performance/version of `K`, never vice versa. The `Q` component of the `uDEF` function output is only used for plotting the optimisation process, and, if desired, can just repeat the value of the `E` component.

##### rDEF

`rDEF` must be of type closure. `rDEF` should be a function that defines a rule by which the `K` coefficient vector is to be altered from one step to another. `rDEF` must accept `K` as an input and return an object equivalent to `K`, but with some alteration(s).

##### K.INITIAL

`K.INITIAL` is an object of any type, which stores the initial values for the parameter(s) to be optimised. The only requirement for `K` is that it should be something alterable *via* `rDEF` and something that influences the outcome of `mDEF`.

#### Optional Input Arguments

##### NUMITER

`NUMITER` is a variable of type double that is the number of steps of the optimisation process per processing core. It has a default value of 1 000 000.

##### STATWINDOW

`STATWINDOW` is a variable of type double that is the number of steps executed between subsequent temperature adjustments executed by the Temperature Control Unit. `STATWINDOW` is also the number of steps used to calculate the observed acceptance ratio). It has a default value of 70.

##### T.INI

`T.INI` is a variable of type double that represents the initial system pseudo temperature at the beginning of the optimisation procedure. It has a default value of 0.00001.

##### T.ADJSTEP

`T.ADJSTEP` is a variable of type double that represents the baseline temperature change step-size for temperature auto-adjustment. It has a default value of 0.000000005.

##### TSCLnum

`TSCLnum` is a variable of type double that indicates the maximum number of `STATWINDOWS` for which the observed acceptance ratio can sequentially be below or above the ideal acceptance ratio before `T.ADJSTEP` is scaled by `T.SCALING`. It has a default value of 2.

##### T.SCALING

`T.SCALING` is a variable of type double that represents the value by which `T.ADJSTEP` is scaled in accordance with the condition specified by `TSCLnum`. It has a default value of 3.

##### T.MIN

`T.MIN` is a variable of type double and is the value that the system pseudo temperature is automatically set to if at any time, the Temperature Control Unit attempts to make the pseudo temperature a negative value. It has a default value of 0.000000005.

##### T.DELTA

`T.DELTA` is a variable of type double. If after a `STATWINDOW`, the observed acceptance ratio is within `T.DELTA` of the ideal acceptance ratio, the Temperature Control Unit will make no change to the system pseudo temperature. It has a default value of 2.

##### DUMP.FREQ

`DUMP.FREQ` is a variable of type double. It is the frequency in steps of printing the best found model to the working directory. It has a default value of 10 000.

##### LIVEPLOT

`LIVEPLOT` is a variable of type logical that indicates whether the optimisation process will be plotted in a PDF file in the working directory. It has a default value of `TRUE`.

##### LIVEPLOT.FREQ

`LIVEPLOT.FREQ` is a variable of type double that indicates the frequency in steps of printing the optimisation process in a PDF (this variable is only relevant if `LIVEPLOT = TRUE`). It has a default value of 100 000.

##### PDFheight

`PDFheight` is a variable of type double that indicates the height of the PDF that is produced (if `LIVEPLOT = TRUE`). It has a default value of 29.

##### PDFwidth

`PDFwidth` is a variable of type double that indicates the width of the PDF that is produced (if `LIVEPLOT = TRUE`). It has a default value of 20.

##### NCPU

`NCPU` is a variable of type double that indicates the number of optimisation replicas to execute. If calling the Acceptance Ratio Replica Exchange mode of Optimus, `NCPU` must be greater than 1. `NCPU` has a default value of 4.

##### LONG

`LONG` is a variable of type logical. If `LONG = TRUE`, a memory-friendly version of Optimus will be activated (in anticipation of a long simulation), and only data from the optimal explored parameter configuration and the last 10 000 optimisation iterations will be stored. `LONG` has a default value of TRUE.

##### SEED

`SEED` is a variable of type double which sets the seed for the random number generator. It has a default value of 840.

##### OPTNAME

`OPTNAME` is a variable of type character that can be thought of as the name of the optimisation process. `OPTNAME` is used when creating the file names of the Optimus output. `OPTNAME = “”` is the default value.

##### DATA

`DATA` is a variable of type list holding any additional data that must be accessed by `mDEF` and `uDEF`. The default value for `DATA` is `NULL`.

##### OPT.TYPE

`OPT.TYPE` is a variable of type character, which specifies the mode of Optimus to execute and should always be equal to “SA” or “RE.” If equal to “SA” (for Simulated Annealing), the Acceptance Ratio Simulated Annealing version of Optimus will be executed. If equal to “RE” (for Replica Exchange), the Acceptance Ratio Replica Exchange version of Optimus will be executed. The default value of `OPT.TYPE` is “SA.”

##### DIR

`DIR` is a variable of type character, which specifies the directory to save the Optimus outputs. The default value for `DIR` is “.” that means the current directory while running Optimus.

##### starcore

`starcore` is an experimental variable of type list, holding some parameters for in-lab starcore use only. For example, `starcore` can be `c(“MAXCOEFORDER”=4)`, which will specify the maximum coefficient order. The default value for `starcore` is `NULL`, i.e. not activated.

#### Optional Inputs Specific to Acceptance Ratio Simulated Annealing

The below optional input arguments only impact the Acceptance Ratio Simulated Annealing mode of Optimus.

##### CYCLES

`CYCLES` is a variable of type double that specifies the number of annealing cycles to execute per core during the optimisation process. It has a default value of 10.

##### ACCRATIO.IN

`ACCRATIO.IN` is a variable of type double that specifies the initial target acceptance ratio for the annealing schedule in each annealing cycle. It has a default value of 90.

##### ACCRATIO.FIN

`ACCRATIO.FIN` is a variable of type double that specifies the final target acceptance ratio for the annealing schedule in each annealing cycle. It has a default value of 0.5.

#### Optional Inputs Specific to Acceptance Ratio Replica Exchange Optimisation

The below optional input arguments only impact the Acceptance Ratio Replica Exchange mode of Optimus.

##### EXCHANGE.FREQ

`EXCHANGE.FREQ` is a variable of type double that specifies the number of optimisation iterations to execute between subsequent parameter configuration exchanges between adjacent replicas. It has a default value of 1000.

##### ACCRATIO

`ACCRATIO` is a vector of doubles that must have length equal to the value of `NCPU`. `ACCRATIO` specifies the target acceptance ratio for each optimisation replica. It has a default value of `c(90, 50, 5, 1)`.

#### Optimus Output

Optimus creates multiple output files in a user’s working directory during an optimisation run. Each core used will generate 4 or 5 output files (depending on the value of `LIVEPLOT`):

1. [OPTNAME][Core number]_model_ALL
2. [OPTNAME][Core number]_model_K
3. [OPTNAME][Core number]_model_O
4. [OPTNAME][Core number]_model_QE
5. [OPTNAME][Core number]

Note that the 5*^th^* file is only produced if `LIVEPLOT=TRUE`. As an illustration, for the Acceptance Ratio Simulated Annealing run from **Tutorial 1**, Optimus generates 20 output files with the following names: poly_4_SA1_model_ALL, poly_4_SA1_model_K, poly_4_SA1_model_O, poly_4_SA1_model_QE, poly_4_SA1, poly_4_SA2_model_ALL, poly_4_SA2_model_K, poly_4_SA2_model_O, poly_4_SA2_model_QE, poly_4_SA2, poly_4_SA3_model_ALL, poly_4_SA3_model_K, poly_4_SA3_model_O, poly_4_SA3_model_QE, poly_4_SA3,poly_4_SA4_model_ALL, poly_4_SA4_model_K, poly_4_SA4_model_O, poly_4_SA4_model_QE and poly_4_SA4.

##### [OPTNAME][Core number]__model__ALL

This is the most important output file in that essentially all contents from the other output files is contained in this file. The *_model_ALL file is an R workspace that contains a variable `OUTPUT` of type list. For Acceptance Ratio Simulated Annealing, `OUTPUT` has 12 fields (numbered 1-12 below), while for Acceptance Ratio Replica Exchange, `OUTPUT` contains an additional 14 fields (numbered 13-26 below). Note that the additional fields of `OUTPUT` in Replica Exchange were included to facilitate the writing of the code; they can largely be ignored by the user.

1. `K.stored`
2. `O.stored`
3. `STEP`
4. `PROB.VEC`
5. `T.DE.FACTO`
6. `IDEAL.ACC.VEC`
7. `ACC.VEC.DE.FACTO`
8. `STEP4ACC.VEC.DE.FACTO`
9. `ENERGY.DE.FACTO`
10. `Q.STRG`
11. `ACCEPTANCE`
12. `STEP.STORED`
13. `E.stored`
14. `E.old`
15. `Q.old`
16. `K`
17. `T`
18. `Step.stored`
19. ENERGY.TRIAL.VEC
20. `STEP.add`
21. `NumofAccRatSMIdeal`
22. `NumofAccRatGRIdeal`
23. `t.adjstep`
24. `AccR.category`
25. `new.T.INI`
26. `instanceOFswitch`

`K.stored` holds the optimal parameter configuration found by the given processor. `O.stored` holds the object `O` generated by `mDEF` from the optimal parameter configuration `K.stored`. `STEP` is a double that holds the current optimisation iteration number. `PROB.VEC` is a vector that holds the acceptance probability for each optimisation step. `T.DE.FACTO` is a vector that holds the system pseudo temperature during each optimisation iteration. `IDEAL.ACC.VEC` is a vector that holds the target acceptance ratio for each optimisation iteration in the case of Acceptance Ratio Simulated Annealing. In the case of Acceptance Ratio Replica Exchange, `IDEAL.ACC.VEC` holds the same value as the input variable `ACCRATIO. ACC.VEC.DE.FACTO` is a vector that holds the observed acceptance ratio at the end of each `STATWINDOW. STEP4ACC.VEC.DE.FACTO` is a vector that holds the optimisation step numbers that correspond to the end of a `STATWINDOW. ENERGY.DE.FACTO` is a vector that holds the actual system energy *E* at each optimisation step. `Q.STRG` is a vector that holds the system quality *Q* at each optimisation step. `ACCEPTANCE` is a vector of binary variables whose *i^th^* entry is 1 if the candidate parameter configuration was accepted on the *i^th^* optimisation step and 0 otherwise. `STEP.STORED` is a vector storing the number of each optimisation step.

`E.stored` is a double that stores the energy *E* associated with the optimal parameter configuration `K.stored`. `E.old` is a double that stores the energy *E* associated with the most recently considered parameter configuration. `Q.old` is a double that stores the quality *Q* associated with the most recently considered parameter configuration. `K` holds the most recently considered parameter configuration. `T` is a double that holds the current system temperature. `Step.stored` is a double that holds the optimisation step number on which the optimal parameter configuration `K.stored` was discovered. ENERGY.TRIAL.VEC is a vector that holds the energy *E* of the candidate parameter configuration at every optimisation step. `STEP.add` is a double that facilitates indexing into output vectors. `NumofAccRatSMIdeal` is a double that represents the number of times the observed acceptance ratio has sequentially been smaller than the target acceptance ratio. `NumofAccRatGRIdeal` is a double that represents the number of times the observed acceptance ratio has been sequentially greater than the target acceptance ratio. `t.adjstep` is a double holding the current value by which the system pseudo temperature is increased or decreased each `STATWINDOW` by the Temperature Control Unit. `AccR.category` is a character that indicates whether the observed acceptance ratio was above or below the target acceptance ratio during the previous `STATWINDOW. new.T.INI` is a double that stores an estimate for the ideal initial system pseudo temperature deduced by the Temperature Control Unit. Finally, `instanceOFswitch` is a double that tracks the number of times the observed acceptance ratio has transitioned from being less than the target ratio to greater than the target ratio or vice versa.

##### [OPTNAME][Core number]_model_K

The *_model_K file is an R workspace that contains a variable `K.stored` of the same type as `K.INITIAL`. It holds the optimal parameter configuration found by the given processor.

##### [OPTNAME][Core number]_model_O

The *_model_O file is an R workspace that contains a variable `O.stored` that holds the object `O` generated by `mDEF` from the optimal parameter configuration `K.stored`.

##### [OPTNAME][Core number]_model_QE

The *_model_QE file is a text file that stores the values of *E* and *Q* that are produced from the optimal parameter configuration `K.stored` and, in the case of the Acceptance Ratio Replica Exchange mode of Optimus, stores the target acceptance ratio associated with the given replica.

##### [OPTNAME][Core number]

The * file is a pdf file that includes 5 plots:

1. acceptance probability as a function of optimisation step;
2. system psuedo temperature as a function of optimisation step;
3. observed (red solid line) and target (black dashed line) acceptance ratio as a function of optimisation step;
4. energy *E* as a function of optimisation step;
5. quality *Q* as a function of optimisation step.

